# PPARγ mediated enhanced lipid biogenesis fuels *Mycobacterium tuberculosis* growth in a drug-tolerant hepatocyte environment

**DOI:** 10.1101/2024.02.02.578554

**Authors:** Binayak Sarkar, Jyotsna, Mohit Yadav, Priya Sharma, Raman Deep Sharma, Shweta Singh, Aakash Chandramouli, Kritee Mehdiratta, Ashwani Kumar, Siddhesh S. Kamat, Devram S. Ghorpade, Debasisa Mohanty, Dhiraj Kumar, Rajesh S. Gokhale

## Abstract

*Mycobacterium tuberculosis* (Mtb) infection of the lungs, besides producing prolonged cough with mucus, also causes progressive fatigue and cachexia with debilitating loss of muscle mass. While anti-tuberculosis (TB) drug therapy is directed toward eliminating bacilli, the treatment regimen ignores the systemic pathogenic derailments that probably dictate TB-associated mortality and morbidity. Presently, it is not understood whether Mtb spreads to metabolic organs and brings about these impairments. Here we show that Mtb creates a replication-conducive milieu of lipid droplets in hepatocytes by upregulating transcription factor PPARγ and scavenging lipids from the host cells. In hepatocytes, Mtb shields itself against the common anti-TB drugs by inducing drug-metabolizing enzymes. Infection of the hepatocytes in the *in vivo* aerosol mice model can be consistently observed post-week 4 along with enhanced expression of PPARγ and drug-metabolizing enzymes. Moreover, histopathological analysis indeed shows the presence of Mtb in hepatocytes along with granuloma-like structures in human biopsied liver sections. Hepatotropism of Mtb during the chronic infectious cycle results in immuno-metabolic dysregulation that could magnify local and systemic pathogenicity, altering clinical presentations.

## Introduction

*Mycobacterium tuberculosis* (Mtb), the causative agent of human tuberculosis, remains the leading infectious killer globally with an estimated death of 1.3 million in 2022[1]. Despite progressive work on designing new anti-TB therapeutics and implementing vaccination programs in TB-endemic countries, it has a high global case fatality rate and a poor treatment success rate, along with a rising number of drug-resistant infections[2, 3]. Emerging paradigms in infectious diseases advocate tackling pathogen-driven ailments as one-dimensional problems, where the entire emphasis is given to pathogen elimination. A holistic understanding of how the host systems respond to the infection, vaccination, and treatment is key to TB management programs[4]. Recent widespread and severe physiological derangements associated with COVID-19 patients, even after the elimination of the virus, have brought back the focus on identifying novel strategies that are inclusive of modulating the host immune-metabolic axis[5, 6].

The clinical symptoms of pulmonary TB encompass localized manifestations like prolonged cough with mucus, pleuritic chest pain, hemoptysis, and lung damage. Besides, systemic outcomes like cachexia, progressive fatigue, oxidative stress, altered microbiota and glucose intolerance result in organ-wide disruptions[7, 8]. Pulmonary TB patients often suffer from progressive and debilitating loss of muscle mass and function with severe weight loss, this TB-associated cachexia cannot be reversed by conventional nutritional support[9, 10]. Besides, numerous epidemiological studies indicate that hyperglycemia may occur during active tuberculosis, which can compromise insulin resistance and glucose tolerance, although the mechanisms are unclear[7, 11, 12]. Both the localized and systemic pathophysiology of TB infection indicate an alteration in the host immuno-metabolic axis. It is somewhat bewildering that the engagement of the liver during the Mtb infection cycle is not considered, despite its central role in balancing the immune and metabolic functions of the body[13]. The crosstalk between the liver and lung has been largely overlooked in tuberculosis (TB), even though acute phase proteins (APPs) are used as predictive biomarkers in pulmonary tuberculosis[14]. A robust hepatic APR response in mice, mediated by key hepatocyte transcription factors, STAT3 and NF-κB/RelA, has been known to trigger pulmonary host defenses for survival during pneumonia and sepsis[15]. In TB, the active phase of the disease is associated with heightened expression of various genes that modulate flux in the lipid metabolic pathways[16-18].

The liver is involved in a wide variety of functions – synthesis of plasma proteins, secretion of various hepatokines, degradation of xenobiotic compounds, and storage of lipids, glucose, vitamins, and minerals[19, 20]. De-novo lipogenesis, secretion of acute phase proteins, hepatokine production, etc. are all directly or indirectly controlled by the hepatocytes, thereby communicating with almost all the organs of the body[19, 21]. Moreover, the liver is actively involved in triacylglycerol synthesis and storage under the intricate regulation of various hormones like insulin, glucagon, and thyroid hormone[22]. All these functions uniquely position the liver as the central regulator of lipid metabolism.

To avert organ damage, the liver maintains tolerogenic properties rendering it an attractive target for various pathogenic microorganisms. Although several studies have indicated the role of both -Mtb virulence components and host factors (immune activation, nitric oxide, IFNγ, intracellular pH and hypoxia) in the generation of Mtb drug tolerance, the role of liver, being the principal center for xenobiotic metabolism needs careful investigation[23-27]. Liver is the hub of both phase I and phase II drug modifying enzymes (DMEs). The levels as well as the activity of both type of DMEs play significant roles in determining the pharmacokinetics and efficacy of various drugs across multiple diseases, from infection to malignancy[28].

In this study, we demonstrate the active involvement of the liver in a murine aerosol TB infection model during the chronic phase and establish hepatocytes as a new replicative niche for Mtb. Using a variety of *in vivo*, *ex vivo*, and *in vitro* techniques we show how Mtb perturbs biological functions within hepatocytes remodeling intracellular growth, localization, and drug sensitivity. Cellular and mass spectrometric studies demonstrate Mtb infection-mediated enhanced fatty acid biogenesis and TAG biosynthesis in the hepatocytes regulated by PPARγ. We propose that infection of hepatocytes by Mtb during the chronic phase can contribute to significant changes in disease progression, TB treatment, and development of infection-induced metabolic diseases.

## Results

### Human miliary tuberculosis patients harbor Mtb in the liver

Mtb infects lungs, and other organs like lymph nodes, pleura, bones, and meninges. There are also isolated case reports of hepatic TB, without providing many pathophysiological consequences[29, 30]. To gain further insights into the involvement of the liver in Mtb infections, we acquired human autopsied liver samples from individuals with miliary tuberculosis and analyzed them for the presence of Mtb bacilli. Hematoxylin and eosin (H and E) staining showed the presence of distinct immune cell infiltration and granuloma-like structures in the infected samples **(Fig 1A and Fig S1C).** We examined the liver specimens, with Mtb specific Ziehl-Nielsen (Z-N) acid-fast and auramine O-rhodamine B stain (**Fig 1B, C**). Both these stains showed distinct positive signals with characteristic rod-shaped bacilli that could be visualized by the acid-fast staining (as indicated by arrows) **(Fig 1B).** We further corroborated our findings by performing fluorescence in-situ hybridization (FISH) using Mtb-specific 16s rRNA probes, where specific signals were observed **(Fig 1D).** The specific staining in FISH eliminates the possibility of non-tuberculous mycobacteria (NTMs) in the tissue specimen and confirms the presence of Mtb infection in the liver. Similar staining in the uninfected liver sections from other individuals did not show any signal. **(Fig S1 A, B**). Although hepatic granuloma is the characteristic histological feature for both local and miliary forms of hepatic TB, the precise involvement of the different cells like Kupffer cells, hepatocytes, stellate cell, liver sinusoidal endothelial cells, hepatic stellate cells, and other cell types have not been studied in detail[31]. Multiplex immunostaining with β-actin antibody and Mtb-specific Ag85B antibody shows the presence of Ag85B signals within the human hepatocytes further confirming the presence of Mtb within hepatocytes (**Fig 1E and Fig S1D)**, indicated with yellow arrows). Hepatocytes are morphologically distinct, large polygonal cells (20-30 µm) with round nuclei, many of which are double-nucleated and mainly positioned at the center of the cytoplasm[32]. The presence of Ag85B signals near the nucleus, as depicted in (**Fig 1E and S1 D)**, further supports our assertion. Moreover, hepatic granulomas in the human samples showed localized clustering of the immune cells **(Fig S1D).** The corresponding Ag85B staining in the uninfected liver biopsy samples did not show any signal (Fig 1F). Furthermore, to establish a correlation between liver Mtb load and Mtb burden at the primary infection site, the lung, we conducted H&E staining and acid-fast staining on lung sections from the same individuals. H&E staining revealed distinct granulomas, while acid-fast staining confirmed an elevated bacterial load, both indicative of a high degree of infection **(Fig S1E, F).** Using auramine O-rhodamine B, Ziehl-Nielsen acid-fast staining, and FISH, Mtb presence was detected in several lung specimens **(Fig S1G).** These results indicate Mtb infection in the liver of human subjects and suggest the localization of Mtb within hepatocytes. These findings are quite intriguing considering very few studies have discussed the lung-liver cross talk or the involvement of liver in pulmonary TB.

**Figure 1:**
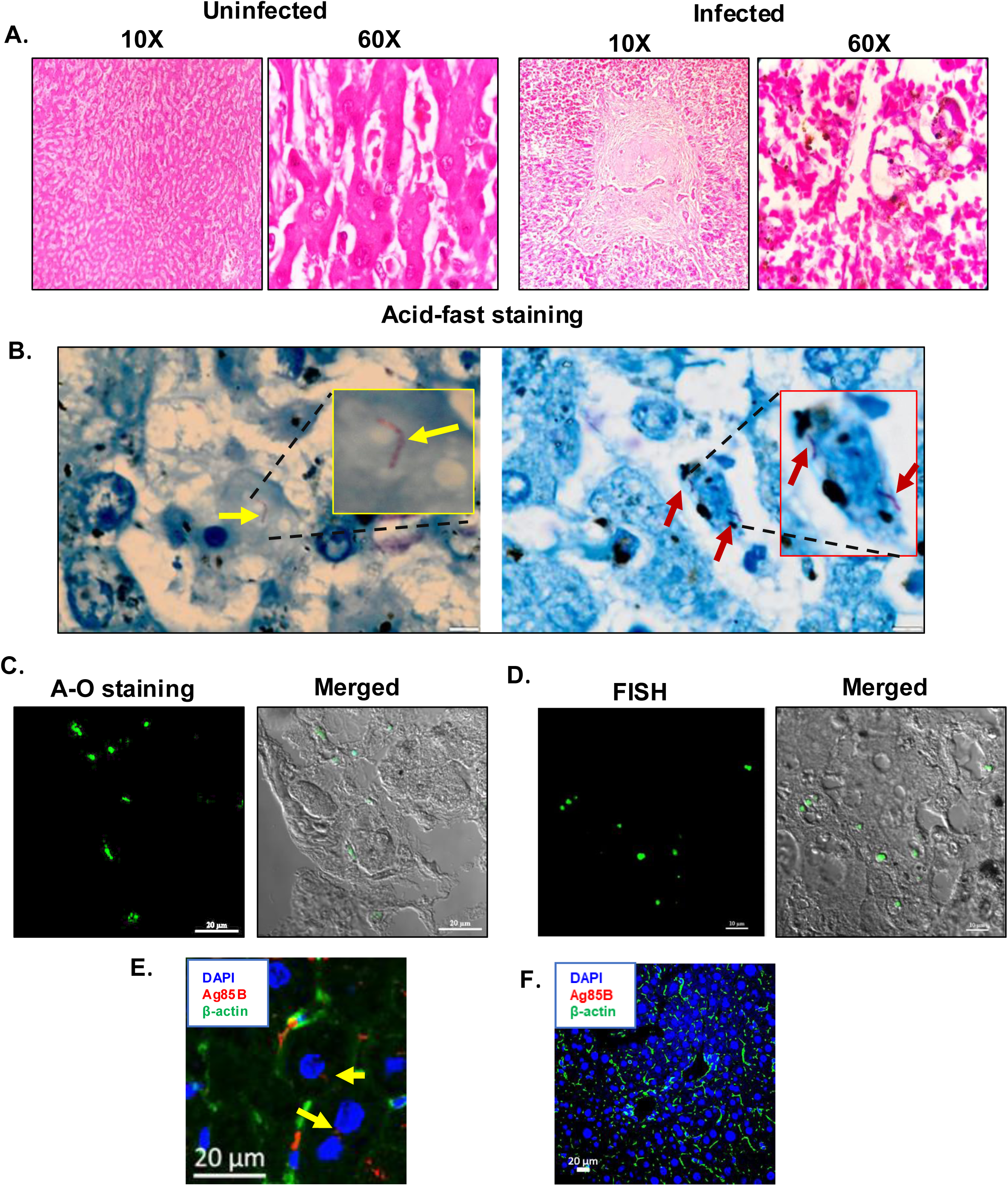
Presence of Mtb in human miliary tuberculosis patients: **(A)** Hematoxylin and eosin (H and E) staining of the human autopsied liver tissue sections showing the presence of granuloma-like structure in the Mtb infected liver. **(B)** Acid-fast staining shows the presence of Mtb in the liver section of miliary TB patients. (arrows point to the presence of Mtb in the enlarged image) **(C-D)** Auramine O-Rhodamine B (A-O) staining and fluorescence in-situ hybridization (FISH) with Mtb specific 16s rRNA primer further confirms the presence of Mtb in human liver tissue sections. **(E).** Dual staining of β-actin (green) and Ag85B (red) using respective antibodies shows the presence of Mtb in hepatocytes of human biopsied liver sections. **(F).** Dual staining of β-actin (green) and Ag85B (red) using respective antibodies shows no distinct signal of Ag85B in the uninfected liver biopsied sample (control).

### Mtb infection of mice via the aerosol route leads to significant infection of the liver and primary hepatocytes

To investigate whether the liver harbours Mtb during mice aerosol infection, we infected C57BL/6 mice with 200 CFU of Mtb *H37Rv* and scored for the bacterial load in the conventional niche-the lung as well as in the liver. Mtb could be detected in the liver consistently across several experiments at 4 weeks post-infection and the bacterial load increased till week 10 **(Fig 2A and B).** Consistent with the Mtb burden, phalloidin and hematoxylin and eosin (H and E) staining of the infected liver at 8 weeks post-infection showed localized cellular aggregation forming ectopic granulomas forming a granuloma-like structure **(Fig 2C and S2C).** Further staining the liver sections with CD45.2 antibody shows clustering of immune cells in Mtb infected mice liver **(Fig S2D).** To assess whether liver infection leads to deranged liver function in the aerosol model, we analyzed the levels of liver functional enzymes like albumin aspartate transaminase (AST), alanine transaminase (ALT), and gamma-glutamyl transpeptidase (GGT) in the sera. Till week 10 post-infection, sera levels of these markers showed significant alterations as the infection progressed, especially in the levels of AST with changes also in the levels of albumin and GGT **(Fig S2B).** Since hepatocytes, the principal parenchymal cells of the liver constitute 70-80 percent of the liver by weight, we hypothesized whether hepatocytes could harbour Mtb in the murine model of infection. To this end, we isolated primary hepatocytes from the mice at different time points post-infection. The purity of the isolated hepatocytes was validated by multiple methods-morphologically hepatocytes can be identified by their distinct hexagonal architecture, round nucleus, some of which are binucleated as observed in Alexa fluor phalloidin 488 and DAPI stained hepatocytes **(Fig 2E).** Flow cytometry staining with antibody against hepatocyte specific marker asialoglycoprotein receptor 1 (ASGR1) protein, confirmed that almost 100 percent of the isolated cells are primary hepatocytes as seen in **(Fig 2F)**. It was further validated by confocal microscopy as all the cultured cells specifically stained for ASGR1 protein, although with variable levels of expression **(Fig 2G).** After thorough confirmation of the identity and the purity of the hepatocytes, the cells were lysed and plated. Like the whole liver CFU, Mtb infected primary hepatocytes (PHCs) at 4 weeks post-infection with substantial bacterial load at week 6 and week 8 **(Fig 2H).** Staining with antibody for Mtb-specific Ag85B protein in both the infected tissue sections and cultured hepatocytes isolated from the *in vivo* infected mice revealed the presence of Mtb within hepatocytes **(Fig 2D and 2I)**. To further strengthen our data, we co-stained the isolated primary hepatocytes from the Mtb infected mice with anti-albumin antibody (albumin is a robust and widely used hepatocyte marker) and Ag85B. As depicted in (Fig 2J), distinct Ag85B signals can be observed in the albumin rich hepatocytes. Besides the aerosol route, infection intraperitoneally also led to hepatocyte infection, also led to hepatocyte infection as early as day 10 **(Fig S2A).** Further, to prove the spread of Mtb to the liver in other model organisms, we infected guinea pigs with 200 CFU of Mtb and analyzed the bacterial load in the lung, liver, and spleen at week 4 and week 8 post-infection **(Fig S3A)**. At both the time points we could observe robust liver infection with granulomatous structure in the liver, proving that the dissemination of Mtb to the liver occurs across multiple model animals **(Fig S3B).**

**Figure 2:**
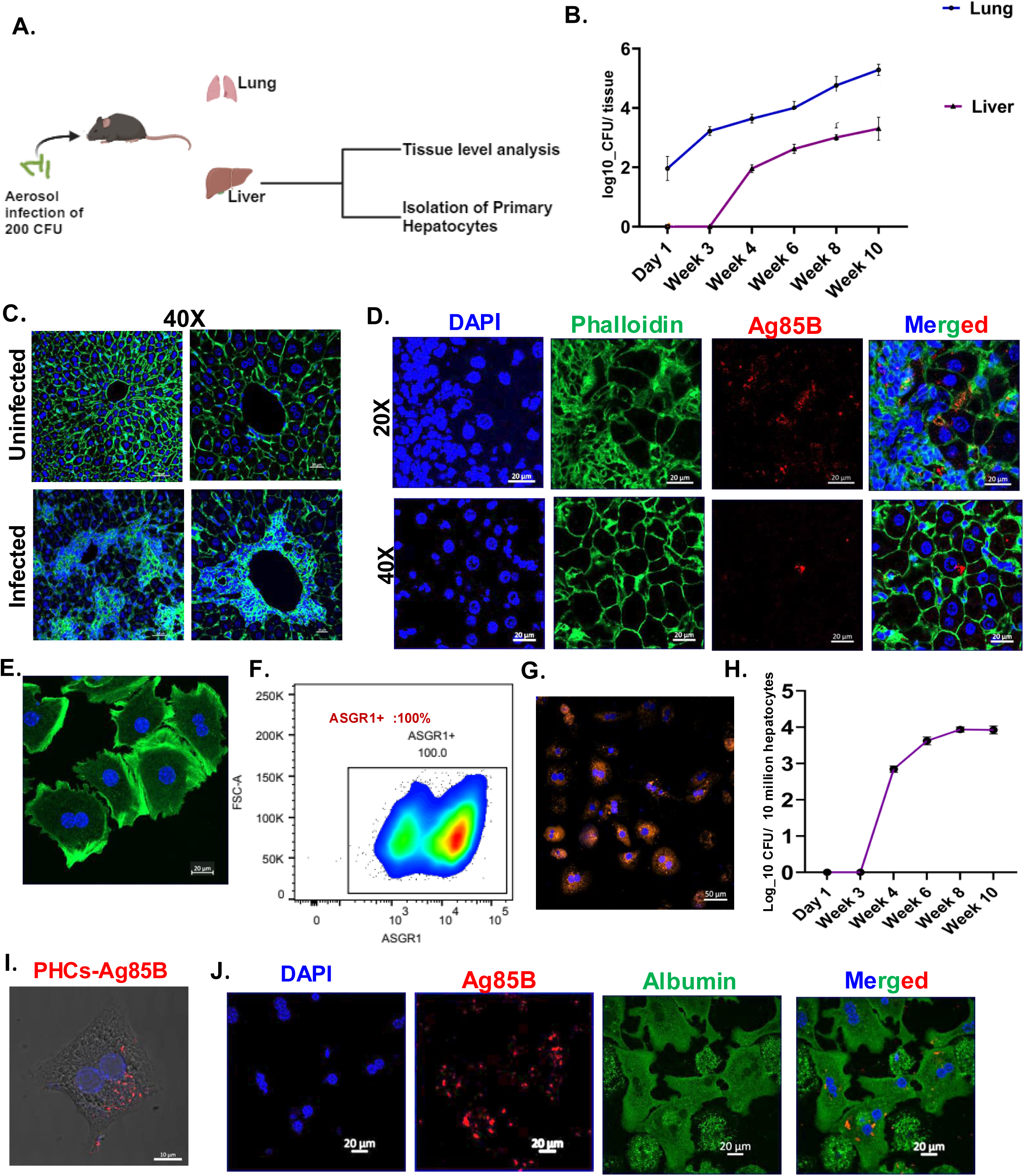
Involvement of the liver and hepatocytes in a mouse aerosol model of Mtb infection. **(A).** Schematic denoting the flow of experimental set up of studying the liver at tissue level and cellular level (generated by Biorender.com). **(B)** C57BL/6 mice were infected with 200 CFU of H37Rv through the aerosol route and the bacterial burden of the lung and liver was enumerated at different time points post-infection in lung and spleen **(C).** Alexa Fluor 488 Phalloidin and DAPI staining of uninfected and infected lungs at 8 weeks post-infection. Images were taken in 40 X magnification as mentioned (scale bar is 20 μm). **(D).** Immunofluorescence staining of Alexa Fluor 488 Phalloidin, DAPI, and Ag85B in the infected mice liver at 8 weeks post-infection shows the presence of Ag85B signals within hepatocytes (magnified image) (scale bar is 20 μm). **(E).** Isolated primary hepatocytes stained with phalloidin green and DAPI show the typical polygonal shape with binucleated morphology. **(F).** Anti-asialoglycoprotein receptor (ASGR1) antibody staining specifically stains primary hepatocytes isolated from infected mice as validated by the contour plot in flow cytometry. **(G).** Antiasialoglycoprotein receptor (ASGR1) antibody staining specifically stains primary hepatocytes isolated from mice as visualized through confocal microscopy. **(H).** Primary hepatocytes were isolated from the infected mice, lysed and CFU enumeration was done at the mentioned time points. **(I).** Ag85B staining of cultured primary hepatocytes, isolated from mice at 8 weeks post Mtb infection. (scale bar is 20 μm). **(J).** Dual immunostaining of Ag85B (red) and albumin (green) shows distinct signals of Mtb within the isolated hepatocytes from infected mice at 8 weeks post-infection. (n)= 6-7 mice in each group.

### Hepatocytes provide a replicative niche to Mtb

Consistent data from human TB patients and aerosol murine model prompted us to develop an in vitro model of Mtb and hepatocyte infection. We examined Mtb infection in several hepatocyte cell lines of mouse and human origin along with primary murine hepatocytes. Infection studies with fluorescently labeled Mtb *H37Rv* were carried out with primary murine hepatocyte cells (PHCs), HepG2, Huh-7, and AML-12 **(Fig 3A, B, and C).** The multiplicity of infection was titrated both in PHCs and HepG2 with the MOI of 1, 2.5, 5, and 10. MOI 10 consistently showed an infectivity of more than 60% in both HepG2 and PHCs **(Fig S4 A, B, and C).** While we acknowledge that MOI 10 is moderately high, several well-cited studies have used MOI 10 for various conventional phagocytes and other non-conventional cell types like mesenchymal stem cells, adipocytes and human lymphatic endothelial cells[33-37]. Hence MOI 10 was selected for further experiments. Macrophage cell lines RAW 264.7, THP-1, and mouse bone marrow-derived macrophages (BMDMs) were used as positive controls. Even though PHCs, HepG2, Huh-7, and AML-12 are not considered to be classical phagocytic cells, all cells showed infectivity of more than 60 percent after 24 hours, comparable to RAW 264.7, THP-1 and BMDMs **(Fig 2A, B).** Analysis of bacterial load in the PHCs post-infection showed colony-forming units (CFU) like RAW 264.7, THP-1, and BMDMs, supporting microscopic observations **(Fig 3C).** Mtb in macrophages is known to remodel the intracellular environment to survive within phagosomes. We studied Mtb growth kinetics within hepatocytes using GFP-labeled Mtb *H37Rv* in PHCs and HepG2 **(Fig 3D and F).** Mean fluorescent intensity measurements showed a consistent increase in GFP intensity in both PHCs and HepG2 with increasing time **(Fig 3E and G).** Fold change in replication dynamics with respect to 5 hours post infection (HPI) showed that, while bacterial growth in macrophages plateaus after 48 hours post-infection, Mtb continues to grow in both HepG2 and PHCs **(Fig 3H**). Relative CFU numbers in the PHCs and HepG2 further supports our data (Fig S4 D-H). Our studies thus establish that hepatocytes, besides being robustly infected by Mtb, also provide a favourable replicative niche for Mtb.

**Figure 3:**
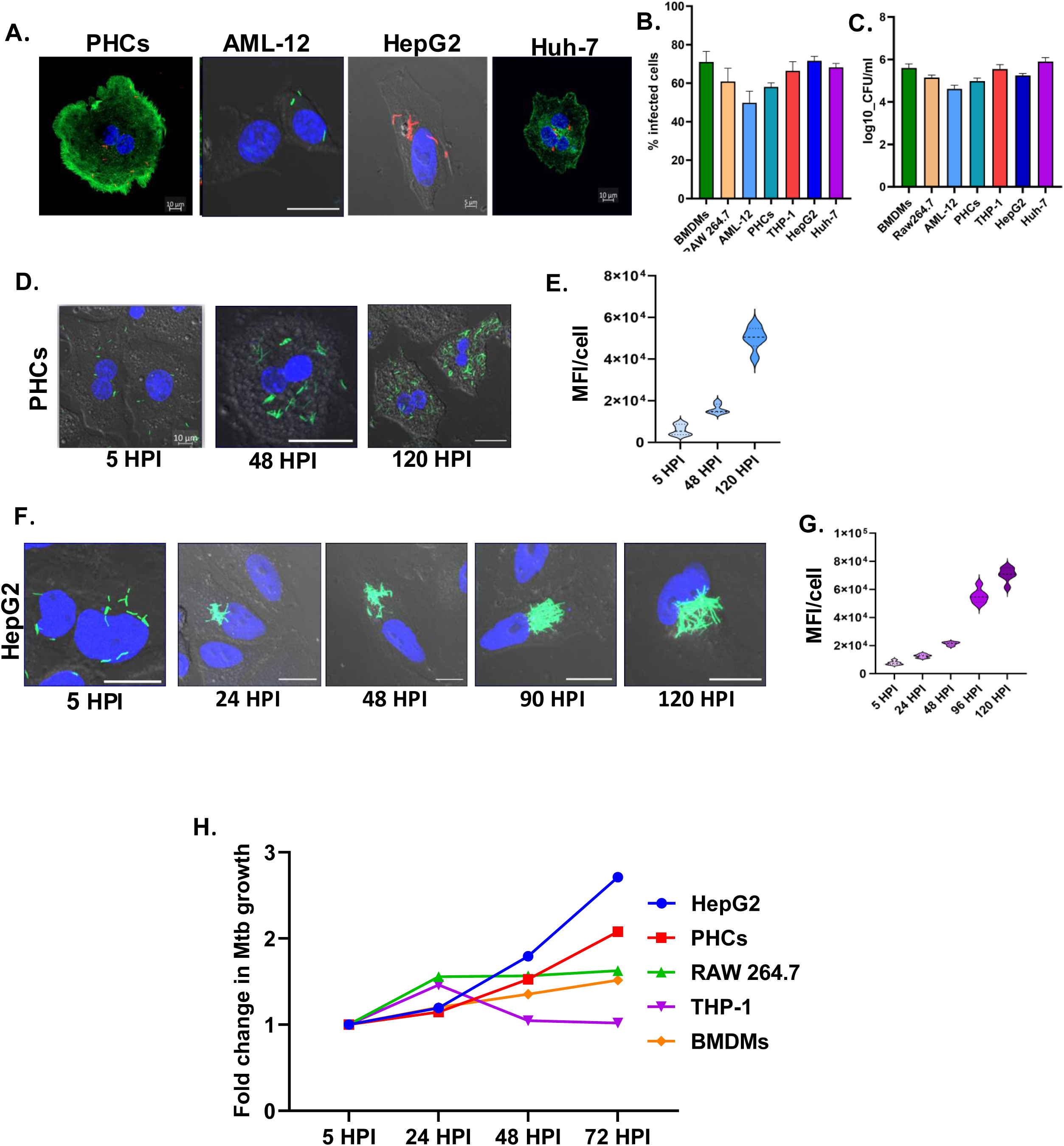
Mtb uses hepatocytes as a replicative niche:. **(A and B)** Representative microscopic images showing the infection of primary hepatocytes (PHCs), AML-12, HepG2, and Huh-7 with labelled Mtb-H37Rv strains and subsequent quantification of percentage infectivity in the respective cell types. RAW 264.7, THP-1 and murine BMDMs were used as macrophage controls. The scale bar in all images is 10 µm except in HepG2, which is 5 µm **(C).** CFU enumeration of Mtb-H37Rv in different hepatic cell lines. **(D and E).** Representative confocal microscopy images of Mtb-GFP-H37Rv infected PHCs at the respective time points post-infection and bar blot depicting mean fluorescent intensity (MFI) / cell at the respective time points. **(F and G).** Representative confocal microscopy images of MtbH37Rv-GFP infected HepG2 at the respective time points post-infection and bar blot depicting MFI/cell at the respective time points, n= 4 independent experiments, with each dot representing 5 fields of 30-60 cells. Scale bar 10 µm. (**H).** Fold change of growth rate of Mtb within HepG2, PHCs, RAW 264.7, THP-1 and BMDMs at the mentioned time points post-infection.

### Transcriptomics of infected hepatocytes reveal significant changes in key metabolic pathways

To understand Mtb-induced changes in the hepatocytes and the underlying mechanisms of how hepatocytes provide a favourable environment to the pathogen, we performed transcriptomic analysis of the infected and sorted HepG2 cells at 0 hours (5 hours post incubation) and 48 hours post-infection. Sorting before RNA isolation specifically enriches the infected cellular population thus eliminating cellular RNA from uninfected cells **(Fig 4A**). Unsupervised clustering segregated the data into 4 distinct groups on the PC1 with a variance of 27 percent, showing good concordance within the replicates **(Fig 4B).** The close spatial clustering for the two 0-hour time points corresponding to uninfected and infected is indicative of relatively less transcriptomic changes. On the other hand, the spatial segregation of the 48 hours datasets suggests clear differences between the RNA transcripts of uninfected and infected cells. The differentially expressed genes were calculated using DE seqR with fold change >0.5 and a false discovery rate of <0.2. Gene ontology (GO) enrichment analysis for the differentially regulated pathways at both the early (0 hours) and the late (48 hours) infection time points is shown in **(Fig 4C).** At 0 hours post-infection, the immediate stress response pathway of the cell, involving ROS generation, intracellular receptor signaling pathways, and response to xenobiotic stresses got activated, while at a late time point, Mtb modulated some of the key immuno-metabolic pathways like macroautophagy, cellular respiration, proteasomal degradation pathway, response to type I interferon, IκB kinase/NF-κB signaling, etc **(Fig 4D).** Major alterations in the vacuolar and vesicular transport at 48 hours are indicative of the dynamic changes in the phagosome maturation pathway **(Fig 4D).** Volcano plot analysis showed greater relative changes in the gene expression pattern at 48 hours compared to 0 hours, with many genes like *CXCL10, CXCL11, IDO, CCL5* etc being greatly upregulated **(Fig 4E, 4F).** Interestingly, our RNA sequencing data indicated major changes in various facets of lipid metabolic pathways like fatty acid biosynthesis pathway, glycerolipid and glycerophospholipid metabolism, cholesterol biosynthesis pathways, etc. Several key genes like *FASN, DGAT1, DGAT2*, *HMGCR*, etc. were upregulated, indicating the possibility of greater synthesis of neutral lipids. Additionally, we have added a supplementary excel file with top 1000 DEGs at both 0 hours and 48 hours post-infection. Thus, transcriptomic studies shed light on several of the key Mtb-induced changes in the hepatocytes.

**Figure 4:**
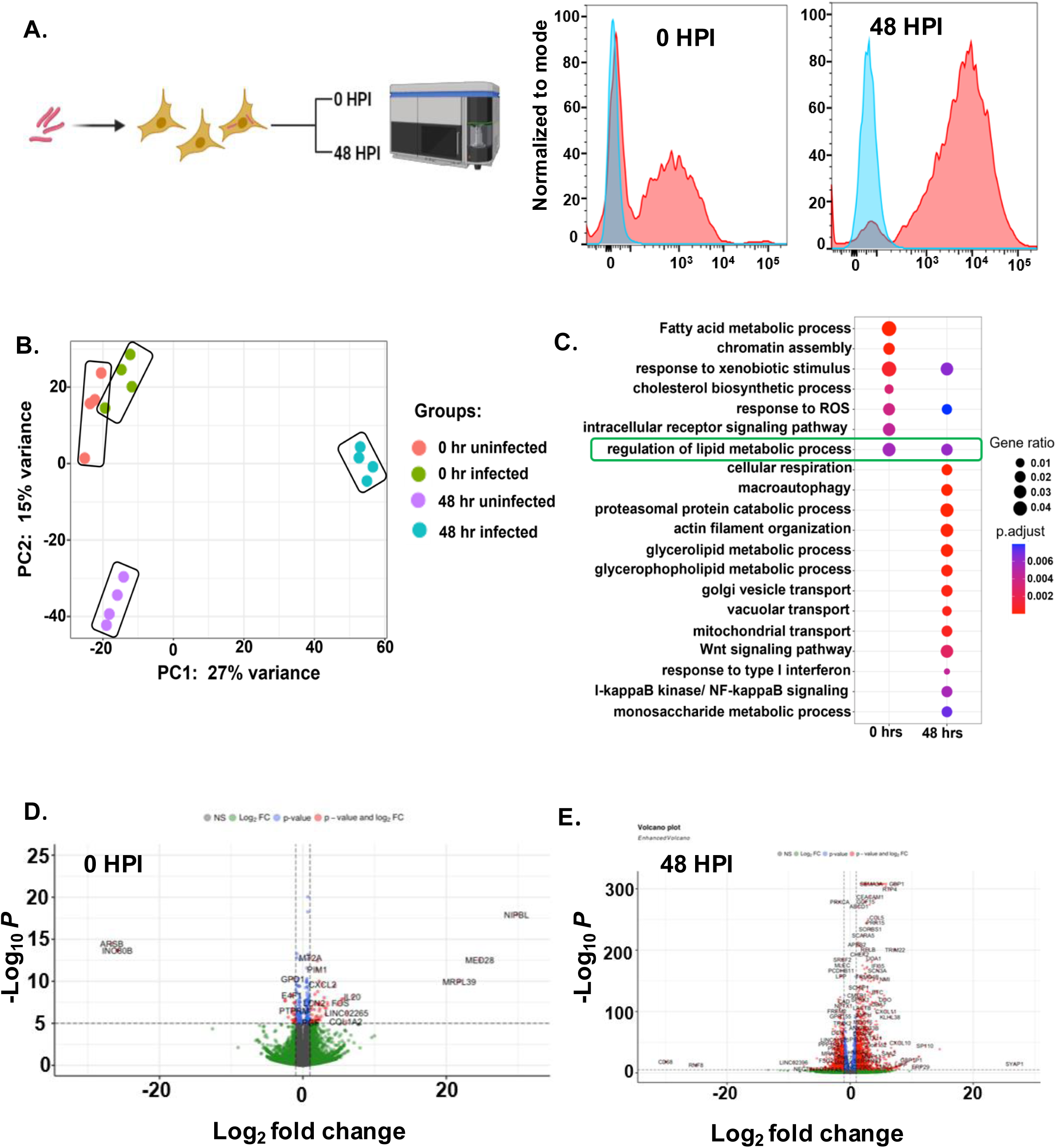
RNA sequence analysis of infected and sorted hepatocytes at 0 hours and 48 hours post-infection: **(A).** Experimental setup for infection of HepG2 with Mb-H37Rv-mCherry at 10 MOI with histogram of mCherry signals at 0- and 48-hours post-infection (Schematic depicting experimental set up generated with Biorender.com) **(B).** Principal component analysis (PCA) plot illustrating the HepG2 transcriptome, identified through global transcriptomic analysis of 0 hours uninfected and infected and 48 hours uninfected and infected. **(C).** GO pathway enrichment analysis was done for DEGs with adjusted p-value < 0.05 at 0- and 48-hours post-infection. The bubble plot depicts the enrichment of pathways on the mentioned time points post-infection, where the coordinate on the x-axis represents the gene ratio, the bubble size represents the gene count and colour represents the p-value. **(D and E).** Volcano plot depicting the fold change of different genes in 0 hours and 48 hours post-infection, the red dots represent the significantly upregulated and downregulated genes.

To understand the difference and similarities between the pathways that get affected in macrophages compared to hepatocytes during Mtb infection, we compared the gene expression analysis data from Mtb-infected HepG2, 48 hours post-infection with THP-1 infected macrophages, 48 post-infection, taken from Kalam et al.,[35] **(Fig S5 A, B).** The comparison provides insights regarding how Mtb modulates different pathways depending on the type of the host cell infected. In THP-1, most of the altered pathways are related to mounting an effective immune response to the bacteria like type 1 interferon signalling pathway, regulation of cytokine secretion, leukocyte chemotaxis, response to zinc, etc while in HepG2 the pathways that are getting altered are related to metabolism like response to xenobiotic stimulus, macroautophagy, glycerophospholipid and glycerolipid metabolism, and cellular respiration. This comparative analysis is indicative of Mtb’s ability to harness the metabolic richness of hepatocytes probably as a source for nutrients. Moreover, being a non-immune cell type, hepatocytes might lack a robust innate immune pathway like the macrophages and hence be less likely to clear mycobacterial infections.

### Increased fatty acid synthesis drives Mtb growth in hepatocytes

Mtb survival in foamy macrophages is driven by nutrient acquisition from the lipid droplets[36]. Transcriptomic studies of Mtb-infected cells also showed upregulated pathways for lipid metabolism. Examination of Lipid droplets in both PHCs and HepG2 revealed an increase in the number of lipid droplets at 24 hours post-infection **(Fig 5A and B).** Time kinetic analysis of BODIPY intensity in the infected HepG2 at different days post-infection indicated a concomitant increase in lipid droplets with the progress of infection **(Fig 5C).** Moreover, in PHCs, GFP labeled Mtb showed a high degree of colocalization with the lipid droplets **(Fig 5D).** Lipid droplets are single membrane-bound depots consisting mainly of neutral lipids like diacylglycerols (DAGs), triacylglycerols (TAGs), and cholesterol esters (CEs)[38].

**Figure 5:**
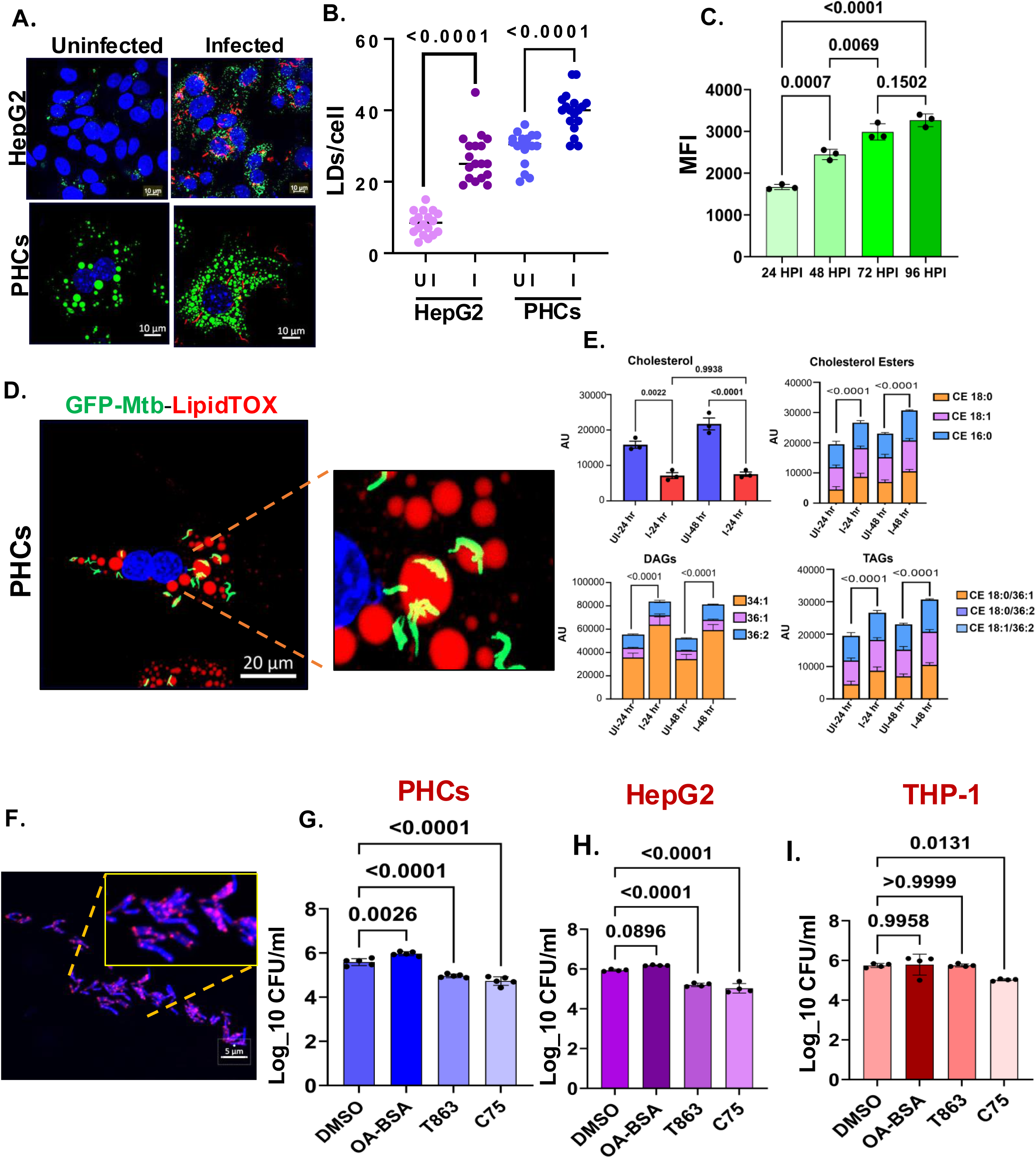
Increased fatty acid biogenesis drives Mtb growth in hepatocytes: **(A)**. Increased number of LDs/cells in HepG2 and PHCs post-Mtb infection as compared to the uninfected cells **(B).** Quantification of the number of lipid droplets in the infected HepG2 and PHCs with their uninfected control, 50-70 cells were analyzed from 3 independent experiments. **(C).** Increase in the BODIPY intensity at different days post-infection in infected hepatocytes **(D)**. A high degree of colocalization of Mtb-*H37Rv-*GFP (green) with lipid droplets (red) within PHCs **(E)** Mass spectrometric quantification of the different species of DAGs, TAGs, and cholesterol esters in the Mtb infected hepatocytes. **(F)**. Confocal images showing puncta of fluorescently labelled fatty acid in Mtb derived from metabolically labelled (with BODIPY 558/568 C_12_) HepG2. Relative CFU of Mtb under the administration of different inhibitors of the lipid metabolic pathway in **(G)**. PHCs **(H)**. HepG2 and **(I)**. THP-1. Representative data from 3 independent experiments. Data were analyzed using the two-tailed unpaired Student’s t-test in B and one-way ANOVA in C, F, G, and H.*p < 0.05, **p < 0.005, ***p < 0.0005, ****p < 0.0001. ns=non-significant.

Mass-spectrometric analysis of the infected and uninfected HepG2 cells at 24 hours post-infection, showed an increase in both TAGs and DAGs and CEs with a decrease in the levels of free cholesterols, indicating Mtb-induced changes in the neutral lipid biosynthesis **(Fig 5E).** To understand whether Mtb utilizes host lipid droplets as a source of nutrients in hepatocytes, we treated Mtb-infected hepatocytes with specific inhibitors of de-novo fatty acid biosynthesis (C75) and TAG biosynthesis (T863) **(Fig S6 A).** Interestingly, inhibiting both de-novo fatty acid biosynthesis as well as TAG biosynthesis reduced the bacterial load in both PHCs and HepG2 by almost 1.0-1.5 log fold **(Fig 5G and H)**. In THP-1 macrophage, although C75 reduced bacterial load by 0.5 log fold but T863 did not affect the bacterial load **(Fig 5I).** To get better insights into the nutritional dependency of intracellular Mtb on hepatocyte lipid source, we metabolically labelled HepG2 with 7.5 μg/ml of fluorescently tagged fatty acid (BODIPY 558/568 C^12^) which subsequently accumulated into host lipid droplets. After 16-20 hours, the labelled cells were treated with T863 and C75. Although T863 significantly reduced the load of the intracellular TAGs, C75 had little effect on the level of accumulated TAGs and showed a phenotype like the DMSO control **(Fig S6 C and D).** We infected these three set of cells with Mtb (MOI:10) for 24 hours. The Mtb isolated from DMSO and C75 treated HepG2 became distinctly labelled with highly fluorescent lipid bodies, but the fluorescence signal in Mtb derived from T863 treated cells was quite low and sparse **(Fig 5F and S7).** Considering that host-derived fatty acids are converted to triacylglycerols (TAGs) in the bacterial cytoplasm as a source of carbon and energy, we wanted to check whether the TAG synthesis machinery was modulated in the hepatocyte-infected Mtb. To this end, we isolated Mtb from HepG2, 48 hours post-infection, and analysed for key genes involved in TAG biosynthesis. Mtb grown in DMEM was used as a control. Interestingly, *Tgs1, Tgs4, Rv 1760*, etc were upregulated by 6-8-fold, indicating an active Mtb transcriptional change in Mtb to utilize and store host-derived lipids **(Fig S6 B).** Cumulatively, our studies thus demonstrate fatty acid biosynthesis and TAG formation to be important for Mtb growth in hepatocytes.

To assess the status of the neutral lipids in the liver, BODIPY staining was conducted in the liver cryosections of uninfected and infected mice (8 weeks post-infection). The liver of Mtb-infected mice bears more lipid bodies **(Fig S8 A).** Co-immunostaining of BODIPY and Ag85B also showed abundant lipid droplets and specific signals of Ag85B **(Fig S8 B).** Next, we stained the liver of Mtb-infected mice at 2-, 4- and 8-weeks post-infection with LipidTOX neutral red dye. Surprisingly we found an elevated signal intensity of the dye at 8 weeks post-infection. This shows that in mice, Lipid droplets might correlate with Mtb burden **(Fig S8 C, D).** Our data comprehensively establishes infection-induced lipid droplet accumulation as a pathogenic outcome of Mtb involvement in the liver during the chronic stage of infection.

### PPARγ upregulation in Mtb-infected hepatocytes leads to augmented lipid biogenesis

To understand the molecular mechanism behind the accumulation of lipid droplets in the Mtb-infected liver at 8 weeks post-infection, we investigated the expression patterns of the transcription factors involved in the regulation of the genes of lipid biogenesis in our RNA-seq data. At 48 hours post-infection, the transcript levels of Peroxisome proliferator-activated receptor-gamma (PPARγ) were upregulated by 5-6-fold. To validate, that we performed quantitative real-time PCR analysis of the *PPAR γ* gene in the infected HepG2 at 48 hours post-infection. With respect to the uninfected control, *PPAR γ* was upregulated by 3-4-fold. Besides, downstream adipogenic genes that are directly or indirectly controlled by *PPARγ* like *MGAT1*, *FSP27*, *FASN*, *DGAT1*, *DGAT2*, *ACAT1, ACAT2, ADIPSIN* etc were all upregulated by more than 2-fold in the infected cells. **(Fig S9 A).** Immunoblot also revealed a greater level of PPARγ protein in primary mouse hepatocytes at 24 hours and 48 hours post-infection in the Mtb infected cells **(Fig 6C, D).** In the in-vivo Mtb aerosol infection model, *Pparγ* expression levels showed an intriguing trend, at week post-infection, the expression level was comparable to the uninfected control, while at week 4 post-infection, the expression level spiked to 1.5-2-fold, reaching 3-4 fold at week 8 **(Fig 6B).** The pattern of expression of *Pparγ* in the infected liver correlated with the bacterial load in the liver, where we see the induction at week 4 when Mtb reaches the liver and maximum expression at week 8 when the load of Mtb is considerably high. Besides, PPARγ, some of the critical enzymes involved in fatty acid biosynthesis, TAG biosynthesis, cholesterol esterification like *Fasn, Dgat1, Dgat2, Mgat1, Acat1, Acat2* etc showed a temporal increase in expression levels at the later time points post-infection **(Fig 6A).** We then used a specific inhibitor of PPARγ, GW9662 (20 µM) and agonist of PPARγ, rosiglitazone (20µM) in infected HepG2 and quantified the bacterial load after 48 hours. Surprisingly, inhibition of PPARγ decreased the bacterial load by almost 2-fold, while chemically inducing PPARγ increased the bacterial load considerably **(Fig 6 E and F, and S9B)**. Moreover, LD number in cells also directly correlated with the levels of PPARγ **(Fig 6G and H).** To investigate whether, PPARγ expression was also induced in the liver of infected mice, we examined PPARγ in the liver post infection. Interestingly, we found enhanced expression PPARγ in the liver of the mice at 8 weeks post infection **(Fig S9 C and D).** Moreover, PPAR-γ intensity in hepatocyte was also high in the infected liver **(Fig S9 E).** Thus, PPARγ activation resulting in lipid droplets formation by Mtb might be a mechanism of prolonging survival within hepatocytes.

**Figure 6:**
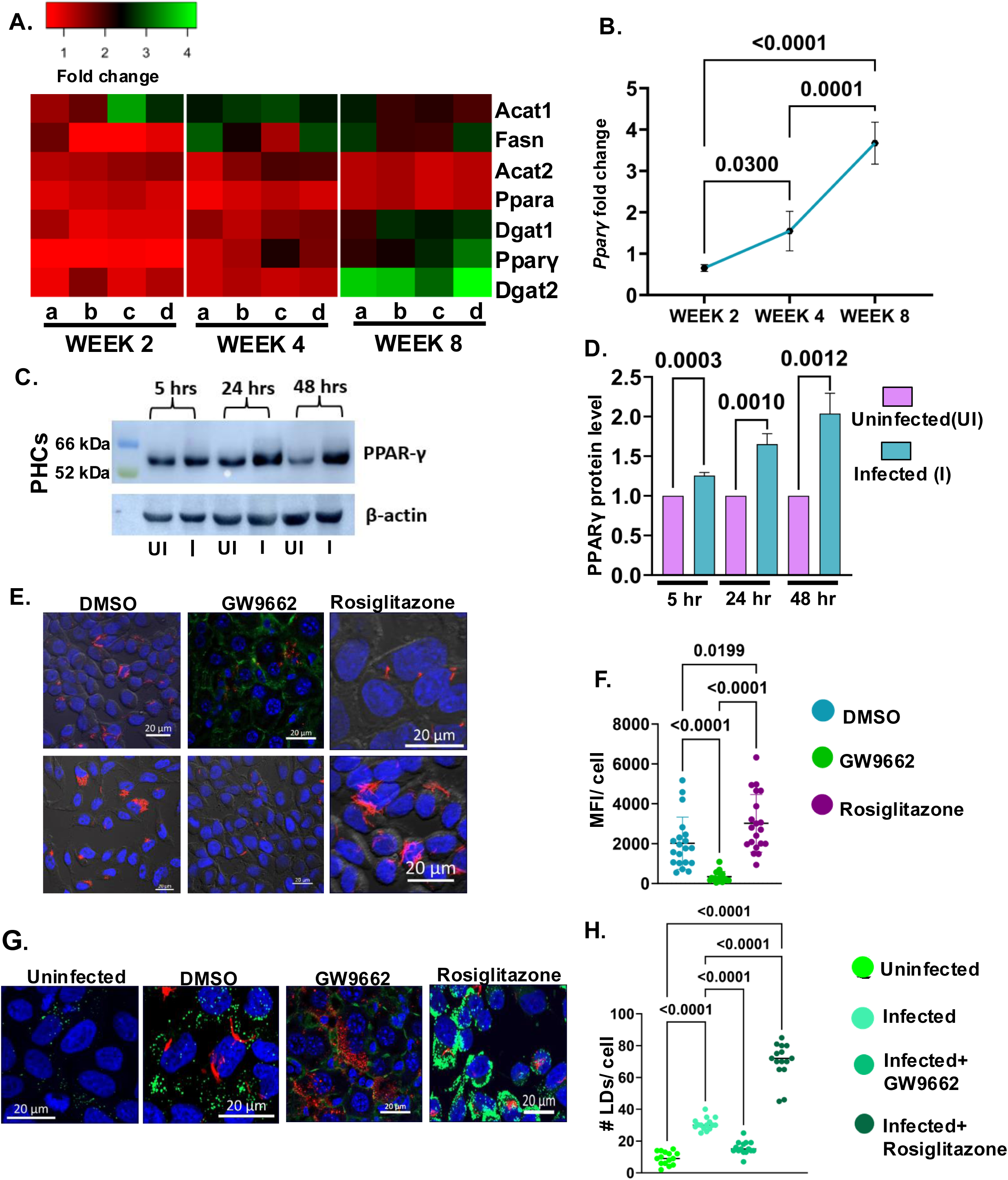
PPARγ driven lipid biogenesis drives Mtb growth in hepatocytes: **(A)** Heat map showing the fold change of the genes involved in the lipid biosynthesis and LD biogenesis in liver across weeks 2, 4 and 8, post infection. **(B)** Kinetic increase in the gene expression of *Pparγ* transcript levels across different weeks post infection **(C)** Immunoblot showing increased PPARγ in PHCs (MOI: 10) at 5-, 24- and 48-hours post-infection. **(D)** Bar plot showing the increased band intensity of PPARγ in the infected PHCs at the mentioned time points. **(E)** Representative confocal microscopy images showing HepG2 infected with Mtb-mCherry-*H37Rv* and treated with GW9662 and rosiglitazone **(F)** MFI/cell of in Mtb-mCherry-*H37Rv* DMSO treated, GW9662 and rosiglitazone treated HepG2 cells. **(G)** Representative confocal images of uninfected, infected, GW9662 and rosiglitazone treated infected HepG2 showing changes in the number of LDs. **(H)** Plot depicting the quantification of LDs/cell in the mentioned conditions. Representative data from n=4 biological replicates. Data were analysed by using the two tailed unpaired Student’s t test in D and by one way ANOVA in B, F and H. *p < 0.05, **p < 0.005, ***p < 0.0005, ****p < 0.0001, ns=non-significant

### Hepatocyte resident Mtb displays a drug-tolerant phenotype

The success of Mtb as a formidable pathogen depends on its ability to tolerate various anti-TB drugs[39]. Drug tolerance is a phenomenom of Mtb surviving drug treatment of longer durations in the absence of any resistance mechanisms. It is a property exhibited by the bacteria but is influenced by both the host and the bacterial factors[23]. Interestingly, hepatocytes are a unique cell type that contain phase I and phase II drug metabolising enzymes[40, 41]. The pharmacological potency of lipophilic drugs is determined by the rate at which these drugs are metabolized to inactive products.

Cytochrome P450 monooxygenases (cyp450s) system is the key phase I DMEs and known to interact with rifampicin[42]. We therefore analysed whether Mtb infection influences the cyp genes in hepatocytes. *CYP3A4* and *CYP3A43* respectively, both of which metabolize anti-TB drug rifampicin were upregulated in Mtb infected HepG2 by approximately 4-fold and 2-fold respectively **(Fig S10A).** Moreover, *NAT2* gene responsible for N-acetylation of isoniazid was also upregulated in the Mtb infected HepG2 by almost 2-fold) **(Figure S10 A).** We therefore argued that hepatocyte resident Mtb may display higher tolerance to rifampicin. Towards this, we treated Mtb-infected HepG2 and PHCs with different concentrations of Rifampicin (0.1, 0.5, 5 µg/ml) for 24 hours and CFU enumerated the bacterial after lysis. RAW 264.7 was kept as macrophage control with the similar experimental setup. The percentage of bacteria which survived the drugs was the drug tolerant population. Both HepG2 and PHCs resident Mtb were significantly tolerant to (25-30%) to rifampicin, as compared to the macrophages **(Fig 7A and B).** Almost 10 percent of the bacterial population in HepG2 display a tolerogenic phenotype at the highest antibiotic concentration **(Fig 7B).**

**Figure 7:**
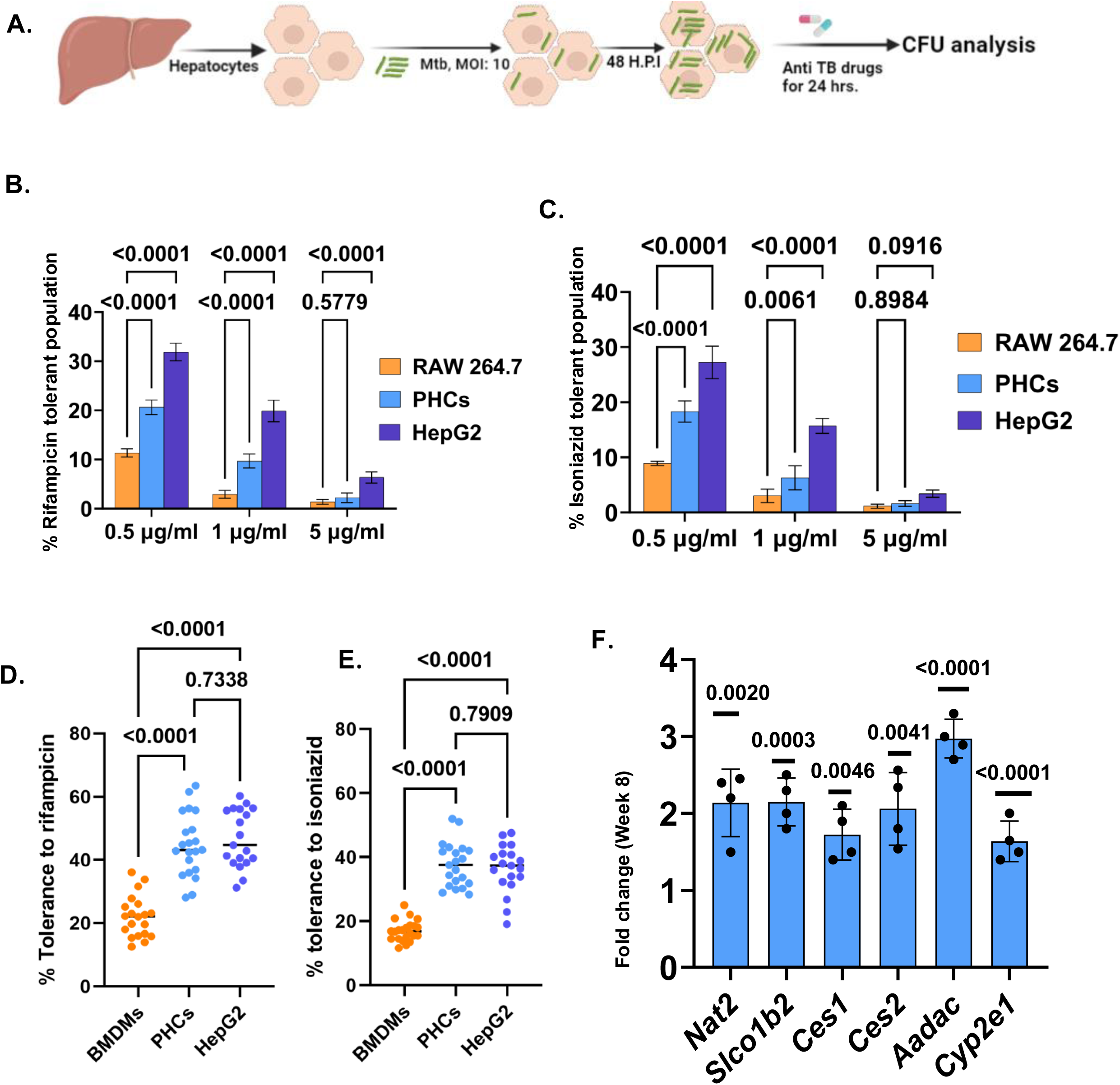
Hepatocytes provide a drug-tolerant niche to Mtb: **(A).** Experimental scheme of deducting the percentage drug tolerance in hepatocytes (generated with Biorender.com) **(B).** Percentage tolerance of Mtb-H37Rv against Rifampicin within RAW 264.7, PHCs and HepG2 at different time points post-infection. **(C).** Percentage tolerance of Mtb-H37Rv against Isoniazid within RAW 264.7, PHCs, and HepG2 at different time points post-infection. Representative data from 3 independent experiments. **(D).** Percentage tolerance of Mtb-*H37Rv* against rifampicin within BMDMs, PHCs, and HepG2 as measured microscopically (**E).** Percentage tolerance of Mtb-H37Rv against isoniazid within BMDMs, PHCs, and HepG2 measured microscopically. Representative data from 3 independent experiments. Each dot represents a single field having more than 4 infected cells. 20 such fields were analyzed **(F)**. Transcript levels of the various DMEs involved in Rifampicin and Isoniazid metabolism in mice liver, 8 weeks post-infection (n=4 mice), the fold change has been calculated by considering the expression in the uninfected mice to be 1. Data were analyzed by using 2-way ANOVA in (B) and (C), one-way ANOVA in (D) and (E) and the two-tailed unpaired Student’s t-test in (F) and Representative data from n=4 biological replicates *p < 0.05, **p < 0.005, ***p < 0.0005, ****p < 0. 0001.ns=non-significant.

We also examined Mtb susceptibility to isoniazid (INH), another first line anti-TB drug which is predominantly metabolized (50–90%) via N-acetylation of its hydrazine functionality by arylamine N-acetyltransferase 2 (*NAT2*)[43]. Interestingly, KEGG analysis of transcriptomic data suggested several genes in this pathway to be upregulated in hepatocytes infected with Mtb **(Fig S10 B).** Experimental studies indeed showed higher tolerance of Mtb to INH in both primary hepatocytes and HepG2 at different concentrations (0.1, 0.5, 5 µg/ml) **(Fig 7A and C).** To corroborate the CFU data, we repeated the same experiment **(Fig 7A)** and looked at the bacterial load within each of murine PHCs and BMDMs using high-resolution confocal microscopy. Mouse BMDMs were as a macrophage control. Here also we have calculated the % tolerance as a ratio by measuring the mean fluorescent intensity of GFP-Mtb per hepatocyte treated with drug to MFI of GFP-Mtb per hepatocyte treated with DMSO (control). Extensive analysis of more than 200 cells distinctly showed presence of more Mtb within PHCs treated with both rifampicin and isoniazid than in BMDMs treated with the same concentration of the drugs **(Fig 7D, E and S11A).**

To check whether some of these key DMEs are also altered in the Mtb infected mice liver, we did qRT-PCR of some key DMEs related top both rifampicin and isoniazid metabolism. Transcript levels of some of the key DMEs (were upregulated in the liver of the infected mice, 8 weeks post-infection like *Nat2, Cyp2e1, Slco1b2*, *Ces1, Ces2, and Aadac* (upregulated by almost 2-3-fold) **(Fig 7F).**

Drug tolerance is a bacterial property influenced by both bacterial and host-induced factors. To check whether Mtb derived from hepatocytes, post-drug exposure has altered drug sensitivity or not, we conducted a resazurin microtitre assay with different dilutions of both rifampicin and isoniazid with Mtb derived from PHCs treated with DMSO, Mtb derived from PHCs post-exposure to rifampicin and Mtb derived from PHCs post-exposure to isoniazid. Mtb from all three different setups did not show any alterations in their MIC values, displaying no change in drug sensitivity (**Fig S12 A**). The increased tolerance of Mtb within the hepatocytes can also be attributed to the increased activity of the drug efflux pumps. Drug efflux pumps like Rv1258c, Rv1410, and Rv1819c are known to get upregulated under drug stress conditions, preventing the accumulation of the drugs and thereby decreasing drug sensitivity. To this end, we isolated intracellular Mtb RNA from HepG2 treated with DMSO, 0.5 μg/ml rifampicin, and 0.5 μg/ml isoniazid and checked for the expression of key drug efflux pumps like *Rv 1258c. Rv1410, Rv1819* etc in these three Mtb populations **(Fig S12 B, C and D).** Interestingly, quantitative RT-PCR analyses of these Mtb genes did not show any upregulation of these transporters in Mtb derived from HepG2 exposed to rifampicin and isoniazid, indicating that drug tolerance phenotype can be better attributed to host intrinsic factors rather than Mtb efflux pumps. Our data show that Mtb-infection of hepatocytes induces DMEs, and this host-specific extrinsic activation may result in decreased bioavailability or increased inactivation of anti-TB drugs.

## Discussion

Over the years, various anatomical and cellular environments conducive to Mtb infection have been identified, particularly concerning its latency[44, 45]. Pulmonary TB typically presents with classical symptoms such as persistent coughing and mucus production due to lung involvement[7]. While systemic manifestations like weight loss, fatigue, and loss of appetite are commonly linked to TB progression, the underlying factors driving these metabolic derangements remain unexplored. Our study sheds light on hepatocytes as a previously unrecognized site for Mtb survival and replication, thereby extrapolating the current understanding of TB beyond its pulmonary focus. By combining *in vitro*, *ex vivo*, *in vivo*, and clinical data, we demonstrate that Mtb not only persists within hepatocytes but also induces important metabolic reprogramming, with significant implications for disease progression, symptomatic manifestations, and drug tolerance.

A central finding of our work is the Mtb-mediated activation of peroxisome proliferator-activated receptor gamma (PPARγ) in hepatocytes. PPARγ activation leads to the accumulation of cholesterol esters (CE 16:0, 18:0, 18:1), diacylglycerols (DAGs 36:1, 36:2, 34:1), and triacylglycerols (TAGs 18:1/36:2, 18:0/36:2, 18:0/36:1). These lipid pools contribute to the formation of lipid droplets, which colocalize with intracellular Mtb, serving as a nutrient reservoir that facilitates bacterial persistence and proliferation. The upregulation of key TAG biosynthesis genes such as *Tgs1*, *Tgs4*, and *Rv1760* in Mtb residing within hepatocytes underscores a transcriptional rewiring within the pathogen to assimilate lipids as a source of their nutrients. Our pharmacological studies using PPARγ agonist and antagonist demonstrated that modulation of PPARγ directly impacts Mtb burden in hepatocytes, confirming the pivotal role of PPARγ in Mtb survival in hepatocytes. The lipid remodeling induced by Mtb infection in hepatocytes is recapitulated in the murine aerosol model, where an increased number of lipid droplets was observed at week 8, accompanied by localized accumulation of immune cells and granuloma-like structures. These findings are consistent with previous reports in macrophages where PPARγ activation enhances lipid biosynthesis, regulates immune responses, and impedes host cell apoptosis[46, 47]. Recent findings have further underscored heightened levels of PPARγ in peripheral blood mononuclear cells (PBMCs) from TB patients with elevated cortisol levels and increased disease severity[48]. However, hepatocytes are uniquely positioned in systemic lipid regulation, engaging in de novo fatty acid synthesis, TAG synthesis, β-oxidation, and lipoprotein metabolism[22, 49]. Mtb-mediated perturbation of these vital processes can lead to a sequela of metabolic disorders such as non-alcoholic fatty liver disease (NAFLD), dyslipidemia and insulin resistance. In macrophages, heightened intracellular lipid levels have been observed to impede autophagy and the acidification of phagolysosomes, both crucial for bacterial eradication[50]. Despite these parallels, comparative transcriptomic analyses of infected HepG2 cells with THP1 cells highlight differences in pathways such as vacuolar and vesicular transport, xenobiotic metabolism, macroautophagy, and cellular respiration. Furthermore, hepatocytes secrete hepatokines, a class of proteins serving as signaling molecules, with diverse roles in metabolism, inflammation, and energy homeostasis, thereby influencing the host-pathogen interplay[13]. Intriguingly, a recent investigation demonstrated that hepatic PPARγ activation induces the expression of growth differentiation factor 15 (GDF-15), a crucial regulator of weight loss observed in ketogenic diets[51].

Additionally, we suggest that Mtb infection of hepatocytes creates a drug-tolerant environment in the liver due to activation of DMEs, many of which are earlier shown to metabolize the two frontline drugs, isoniazid and rifampicin. Moreover, lipid accumulation in the liver might also indirectly alter the levels of drug metabolizing enzymes as reported in previous studies[42, 52]. Upregulation of transcripts is not only observed in infected cells *in vitro*, but also in the livers of mice eight weeks after Mtb infection. Particularly noteworthy is the upregulation of NAT-2, which controls the rate-limiting step of acetylating isoniazid into acetylisoniazid. This metabolite is further processed into acetylhydrazine and isonicotinic acid, ultimately reducing the drug’s effectiveness against Mtb. Similarly, key rifampicin-metabolizing esterase genes such as Ces1, Ces2, Aadac, and transporter Slco1b2 exhibit upregulation, potentially influencing the drug’s distribution and metabolism in the body. The activation of DMEs can significantly modify the pharmacokinetics and pharmacodynamics of anti-TB medications, resulting in suboptimal drug concentrations and the emergence of drug-resistant strains[53].

Pulmonary tuberculosis (TB) can lead to miliary through hematogenous dissemination, where Mtb spreads from the infected lungs into blood vessels either from a primary lung focus, reactivated TB or caseous necrosis. Once in blood vessels, the bacteria seed multiple organs, forming tiny granulomas, characteristic of miliary TB. The liver becomes involved either through direct hematogenous spread or extension from nearby infected lymph nodes, leading to hepatic TB, which presents with granulomas and liver dysfunction. This systemic spread underscore the severity of untreated pulmonary TB and the need for early intervention[54-57]. Our murine and guinea pig aerosol infection models demonstrated progressive liver involvement starting at week 4, with the presence of lipid-laden hepatocytes, localized immune infiltrates, and granuloma-like structures. Importantly, liver biopsy samples from TB patients revealed ectopic granulomas and Mtb antigen (Ag85B) localization within hepatocytes, highlighting the clinical relevance of our findings. While previous research has explored liver infection by Bacillus Calmette-Guérin (BCG) using an intra-venous model in mice, the bacilli were predominantly found within tissue-resident macrophages or Kupffer cells at 15 minutes and 2 days post-infection[58]. However, our study consistently identified the presence of Mtb within hepatocytes in a murine aerosol infection model after the fourth week. Similarly, in a guinea pig aerosol infection model, liver infection was evident, marked by distinct granulomas by week 4. Furthermore, analysis of human biopsy liver samples from pulmonary TB patients revealed ectopic granuloma-like structures within the liver hepatocytes, with the presence of Mtb-specific Ag85B signals within hepatocytes. Together, these findings underscore hepatocytes as a novel niche for Mtb persistence, shedding new light on the pathogenesis of TB. Interestingly, in infected armadillos, *M. leprae* is shown to infect hepatocytes and hijack liver homeostatic, regeneration pathways to promote *de novo* organogenesis[59].

Although animal models offer valuable insights into Mtb infection and TB pathology, they fall short of fully replicating the complexities of human TB. Several recent studies in COVID-19 patients have shown that hepatocytes get infected with SARs-CoV-2, thereby increasing gluconeogenesis and hyperglycemia[60, 61]. With compelling reports associating hepatic steatosis with the onset of type 2 diabetes mellitus (T2DM), we speculate that TB-induced perturbations in lipid metabolism might predispose chronic TB patients to T2DM and vice versa[62, 63]. Through our investigation, we propose that future studies in human TB patients might scrutinize this metabolically rich hepatocyte niche to understand multiple organ-wide derangements in TB pathogenesis.

## Materials and Methods

### Ethical clearances

#### Human studies

Paraffinized non-Mtb infected, and Mtb-infected human autopsied liver specimens were obtained from the Postgraduate Institute of Medical Education and Research, Chandigarh, India. The Institutional Ethics Committee of CSIR-IMTech (Council of Scientific and Industrial Research-Institute of Microbial Technology) approved all research experiments involving human samples (Approval no [IEC (May 2021) #6]). The samples utilized in the study are from a library of autopsied human specimens in PGIMER (Postgraduate Institute of Medical Education and Research). The consent is provided by the relatives of the deceased person while its submission to the PGIMER.

#### Animal studies

All the mice (C57BL/6) were housed in the animal house at the National Institute of Immunology (NII) before being transported to the TACF-ICGEB (Tuberculosis Aerosol Challenge Facility), for subsequent animal infection studies. The animal experiments were performed adhering to the institutional guidelines (Approval number: Institutional Animal Ethical Committee, IAEC 543/20.). Female Hartley Dunkan guinea pigs were used for infection studies in TACF-ICGEB in accordance with the institutional guidelines (Approval number: Institutional Animal Ethical Committee, IEAC #440/17)

#### Confocal microscopy and Immunofluorescence measurements

Lipid droplets were stained by BODIPY 493/503 dye. Primary mouse hepatocytes and HepG2 cells were grown on 12 mm coverslips at respective densities as per the experimental requirement. Cells were washed with 1X PBS (HiMedia M1452-500G), fixed by 4% paraformaldehyde, and incubated at room temperature for 15 minutes. After incubation, the cells were washed thrice with 1X PBS followed by staining with 10 μM BODIPY dye in 1XPBS for 45 minutes at room temperature. After the staining, the excess dye was removed by washing thrice with 1X PBS. To check for the acidified compartments within the cells, LysoTracker Red DND-99 (L7528) was added to the cells at a concentration of 500 nM for 30 minutes at 37° Celsius, followed by washing thrice with 1X PBS to remove the residual dye. The cells were fixed with 4% paraformaldehyde as previously mentioned. For staining with various antibodies, the cells were permeabilized with 0.2% Triton-x 100 (X-100-1L) for 15 minutes, followed by proper washing with 1X PBS. The cells were then blocked by 2% BSA in 1X PBST for 1 hour at room temperature. Post blocking, the cells were treated with primary antibody overnight at 4°Celsius, followed by proper washing with 1X PBS. Secondary antibody (1:500 dilution) was added to the cells for 1 hour at room temperature. Three washes with 1X PBS were given. The nucleus was stained with DAPI (Sigma D9542-5MG) at 1ug/ml concentration for 20 minutes at room temperature. The excess DAPI was washed with 1X PBS followed by mounting with ProLong Gold Antifade mountant (P36930). Images were acquired by Zeiss LSM 980 Laser scanning confocal microscope.

#### Image Analysis

Analysis was done using Image J and Zen blue software. Mean fluorescent intensity/ cell was calculated by corrected total cell fluorescence (CTCF) = Integrated Density – (Area of Selected Cell x Mean Fluorescence of Background readings). Signal intensity in tissue sections were normalized to respective areas. For colocalization of Mtb with various intra-cellular markers, each data point represents a single field with 6-8 infected cells. More than 100 cells across 4 independent experiments were analyzed. Serial confocal sections were acquired with Z-stacks spanning 10-15 uM. The percentage of Mtb in the respective markers were calculated by calculating the Mtb voxels colocalizing in the respective markers. The image was analyzed by Zen Blue software (Zeiss) and Image J.

For percentage tolerance of Mtb within hepatocytes, we have calculated the ratio by measuring the mean fluorescent intensity of GFP-Mtb per hepatocyte treated with drug to MFI of GFP-Mtb per hepatocyte treated with DMSO (control). BMDMs were used as the macrophage control. More than 20 fields, each consisting of more than 4 infected cells have been used for analysis. MFI/ cell was calculated using the Zeiss ZEN blue image analysis software

### Tissue Immunofluorescence staining

#### Paraffin sections

Five-micron thick sections of paraffin-embedded tissue sections were taken in poly-L-Lysine coated slides (P0425-72EA). Deparaffinization was performed by heating the slides at 50°C for 20 seconds (3 times) till the wax melts, followed by the subsequent steps, 100% xylene (Merck, CAS-1330-20-7) for 10 minutes (3 times), xylene and absolute ethanol (Merck, CAS-64-17-5) for 10 minutes, 100% ethanol for 10 minutes, 70% percent ethanol for 5 minutes (2 times), 50% ethanol, distilled water for 5 minutes (2minutes) and a final wash in 1x PBS for 5 minutes (2 times). Antigen retrieval was performed in an antigen retrieval buffer (10mM Sodium Citrate, 0.05% tween-20, pH: 6) by heating the slides at 60°C for 15 minutes. After antigen retrieval, permeabilization was performed with 0.4% Triton-X 100 in 1X PBS for 20 minutes followed by proper washing with 1x PBS. Blocking was done with 5% BSA for 1 hour. Sequential addition of primary antibody was performed at 1:100 dilution at 4°C overnight. Primary antibody was washed with 1X PBS followed by counter stain with DAPI nuclear stain at 1 ug/ml concentration. Mounting was done with a drop of vectashield (sigma-aldrich, F6182-20ml). The slides were visualised in Zeis LSM 980 confocal microscopy at 40X (oil) magnification.

#### Cryosections

Seven-micron thick cryosections were taken in poly-L-Lysine coated slides (P0425-72EA). The sections were washed with 1X PBS, 3 times for 5 mins each. Permeabilisation was done with 0.25% Triton-X 100 in 1X PBS for 15 minutes followed by washing with 1X PBS. Blocking was done with 5% BSA for 1 hour followed by incubation with primary antibody overnight at 4°C. After that the slides were washed with 1X PBS 2 times for 5 minutes each followed by incubation with fluorophore conjugated secondary antibody for 45 minutes. The slides were washed with 1X PBS followed by counterstain with DAPI at 1ug/ml concentration. Mounting was done with a drop of Vectashield (Sigma-Aldrich, F6182-20ml). The dilution of the dyes and antibodies used in the staining are mentioned in the table.

For staining with BODIPY, the sections were incubated with 15 ums of BODIPY for 40 minutes at room temperature. The slides were visualized in Zeis LSM 980 confocal microscopy at 40X (oil) magnification.

#### Fluorescence in-situ hybridization

FISH was used to detect Mtb in infected human liver following published protocols (1-3). Briefly, the paraffinized human liver tissue sections were initially deparaffinized using a serial washing step with xylene and ethanol, following which the sections were treated with 1 mg/ml Proteinase K and 10 mg/ml Lysozyme in 10 mM Tris (pH 7.5) at 37°C for 30 min. Next, the samples were incubated in the Prehybridization buffer at 37°C for 1 h. Prehybridization buffer is composed of 20% 2X Saline sodium citrate (SSC), 20% Dextran sulfate, 30% Formamide, 1% 50X Denhardt’s reagent, 2.5% of 10 mg/ml PolyA, 2.5% of 10 mg/ml salmon sperm DNA, 2.5% of 10 mg/ml tRNA. The slides were thoroughly washed with a 2X SSC buffer. The sections were then incubated in hybridization buffer at 95°C for 10 min and then chilled on ice for 10 min. Further hybridization was allowed at 37°C overnight. Hybridization buffer is composed of prehybridization buffer plus 16S Mtb-H_37_ Rv probe (5′ FITC – CCACACCGCTAAAG – 3′), which is specific for the 16S rRNA of Mtb at a final concentration of 1 ng/μl. The liver tissue sections were next subjected to a series of washing steps with 1X SSC at room temperature for 1 min, 1X SSC at 55°C for 15 min, 1X SSC at 55°C for 15 min, 0.5X SSC at 55°C for 15 min, 0.5X SSC at 55°C for 15 min, 0.5X SSC at room temperature for 10 min. Coverslips were mounted on glass slides and visualized using Nikon A1R confocal microscope with a 488 nm laser.

#### Acid Fast Staining and Auramine O and Rhodamine B staining in liver sections

Acid-fast staining was performed using a ZN Acid Fast Stains-Kit (K005L-1KT, HIMEDIA). Prior to staining, the paraffinized samples were deparaffinized using a serial washing step with xylene and ethanol 1. 100% Xylene for 6 min, 2. Xylene: Ethanol 1:1 for 3 min, 3. 100% Ethanol for 3 min, 4. 95% Ethanol for 3 min, 5. 70% Ethanol for 3 min, 6. 50% Ethanol for 3 min, 7. Distilled water. The glass slides were flooded with Carbol Fuchsin stain and heated to steam for 5 min with a low flame. The glass slides were allowed to stand for 5 min without further heating. The glass slides were then washed in running tap water. The glass slides were decolorized with acid-fast decolourizer for 2 min. 5. Washed with tap water. 6. Counterstain for 30 sec with Methylene Blue Washed with tap water, dried in air, and examined under 100x objective with oil immersion. The presence or absence of bacteria in infected and uninfected samples was checked through staining with Phenolic Auramine O-Rhodamine B dye (4) (1/3 dilution of stock solution) (Auramine O-861020-25gm, Sigma) (Rhodamine B-R6626-100gm, Sigma). Coverslips were mounted on glass slides and visualized using CLSM with a 488 nm laser.

#### Bacterial cultures and in-vitro experiments

Virulent laboratory strains of H37Rv, BCG, and GFP-H37Rv bacterial cultures were cultivated on 7H9 medium (BD Difco) supplemented with 10% Oleic Acid-Albumin-Dextrose-Catalase (OADC, BD, Difco),0.05% glycerol and 0.05% tween 80 under shaking at 37°C conditions for in vitro assays. The cultures were then incubated in an orbital shaker at 100 rpm and 37 °C until the mid-log phase. pMN437-GFPm2 vector (Addgene, 32362) was used to electroporate the virulent H37Rv strain to create GFP-H37Rv, which was then maintained in 50 μg/ml hygromycin 7H9-OADC medium. pMSP12: mCherry plasmid (Addgene No. 30169) was electroporated in H37RV to generate Mtb-H37Rv-mCherry.To prepare the single-cell suspension needed for infection tests, bacterial cultures were passed through a sequence of different gauge needles five times through 23-gauge, 26-gauge, and three times through 30-gauge.

Human monocytic cell line THP-1 were obtained from American Type Culture Collection (ATCC) and cultured in RPMI-1640 medium with 10% FBS at 37 °C and 5% CO2 incubator. THP-1 derived macrophages were obtained by incubating THP-1 cells with 20 ng/ml phorbol 12-myristate 13-acetate (PMA, sigma) for 24 hours followed by washing and maintenance in complete media. The cells were kept in non-PMA containing complete media for 16-20 hours before infection with Mtb. HepG2, Huh-7, RAW 264.7 and PHCS were grown in DMEM under with 10% FBS at 37 °C and 5% CO2 incubator.

AML-12 cells were cultured in DMEM media containing 1X insulin-transferrin-selenium supplement (ITS-G), under the above-mentioned conditions.

In-vitro and ex-vivo infection experiments in primary cells and different cell lines were performed at a Multiplicity of infection of 10 (MOI: 10) for both CFU enumeration and confocal microscopy. The macrophage experiments involving THP-1 and RAW 264.7 involved incubating the cells with Mtb-H37Rv for 5-6 hours followed by washing with 1X PBS and amikacin treatment (200 ug/ ml for 2 hours) to remove the extracellular bacteria. The cells were kept for the designated time points for 24 hours, 48 hours and so on and then lysed with lysis buffer (0.05 % SDS in 1X PBS) followed by plating in 7H11 plates.

For primary mouse hepatocytes and AML-12 cells, the cells were infected with the Mtb at a multiplicity of 10 for 8 hours followed by amikacin treatment (200 ug/ ml for 2 hours). For HepG2 and Huh-7, the time of incubation was 5-6 hours followed by washing with 1X PBS and amikacin treatment as previously mentioned to remove the extracellular bacteria.

The percentage drug tolerant population was calculated using the following formula (A/B X 100) where A is the CFU in the drug treated group, B is the CFU in the untreated group. The cells were morphologically checked for signs of cell death before proceeding with the plating. For the inhibitor and inducer experiments, the cells were treated with the respective drugs for 48 hours post uptake of Mtb. After that they were lysed and Mtb CFU was enumerated.

#### Standardization of Multiplicity of infection (MOI) for PHCs and HepG2

To standardise the MOI, HepG2 and BMDMs were infected with the following MOI of GFP-Mtb as per the standardized protocol: 1, 2.5, 5 and 10. 6 hours post-infection, the cells were trypsinised and fixed with 2% PFA. The percentage uptake was calculated by flow cytometry. For, PHCs, the cells were infected at the above-mentioned MOI and percentage infection was calculated by microscopy as (# of cells with GFP-Mtb/Total # of cells *100)

#### C57BL/6 aerosol challenge

The mice infection experiments were conducted in the Tuberculosis Aerosol Challenge Facility (TACF, ICGEB, New Delhi, India). C57BL/6 mice were placed in individual ventilated cages within the enclosure, maintaining a temperature of 20-25°C, 30-60 % humidity and 12h-12h of light-dark cycle. Following the standardized protocol, mice were infected with 200 CFUs of H37Rv in a Wisconsin-Madison chamber. To ensure proper establishment of infection, two animals were euthanized 24 hours post aerosol challenge. The lungs were harvested and homogenised in 1X PBS and plated in Middlebrook 7H11 agar plates (Difco) supplemented with 10% OADC and 0.5% glycerol. CFU enumeration was done three weeks post plating.

#### C57BL/6 peritoneal infection

The mice were injected with 10^6^ CFUs of H37Rv in 0.2 ml of 1X PBS. Following infection, on different days post infection, the lung, spleen, and the liver was harvested and homogenized and plated in Middlebrook 7H11 agar plates (Difco) supplemented with 10% OADC and 0.5% glycerol. CFU enumeration was done three weeks post plating.

#### List of antibodies and dyes used in the study

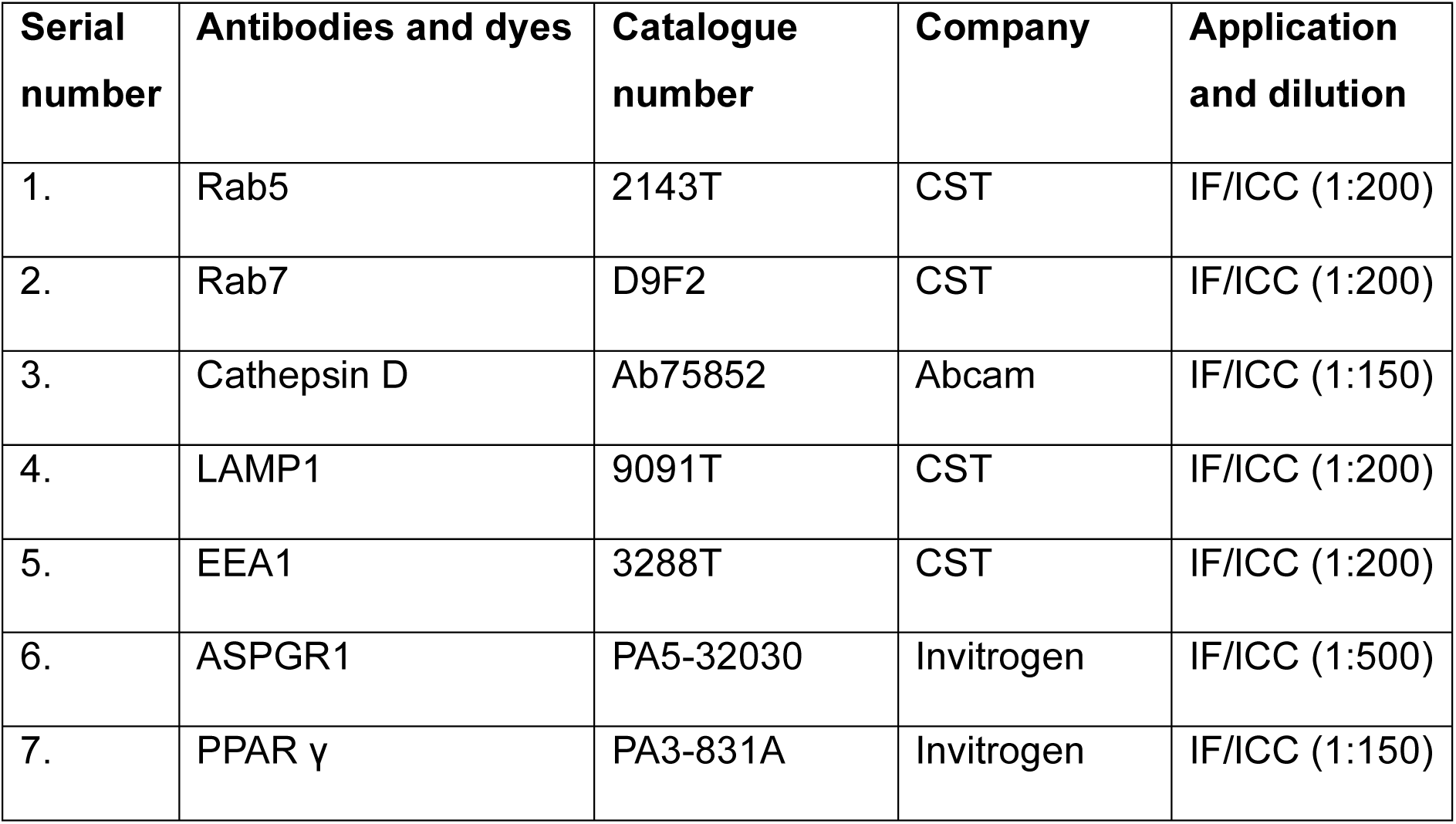
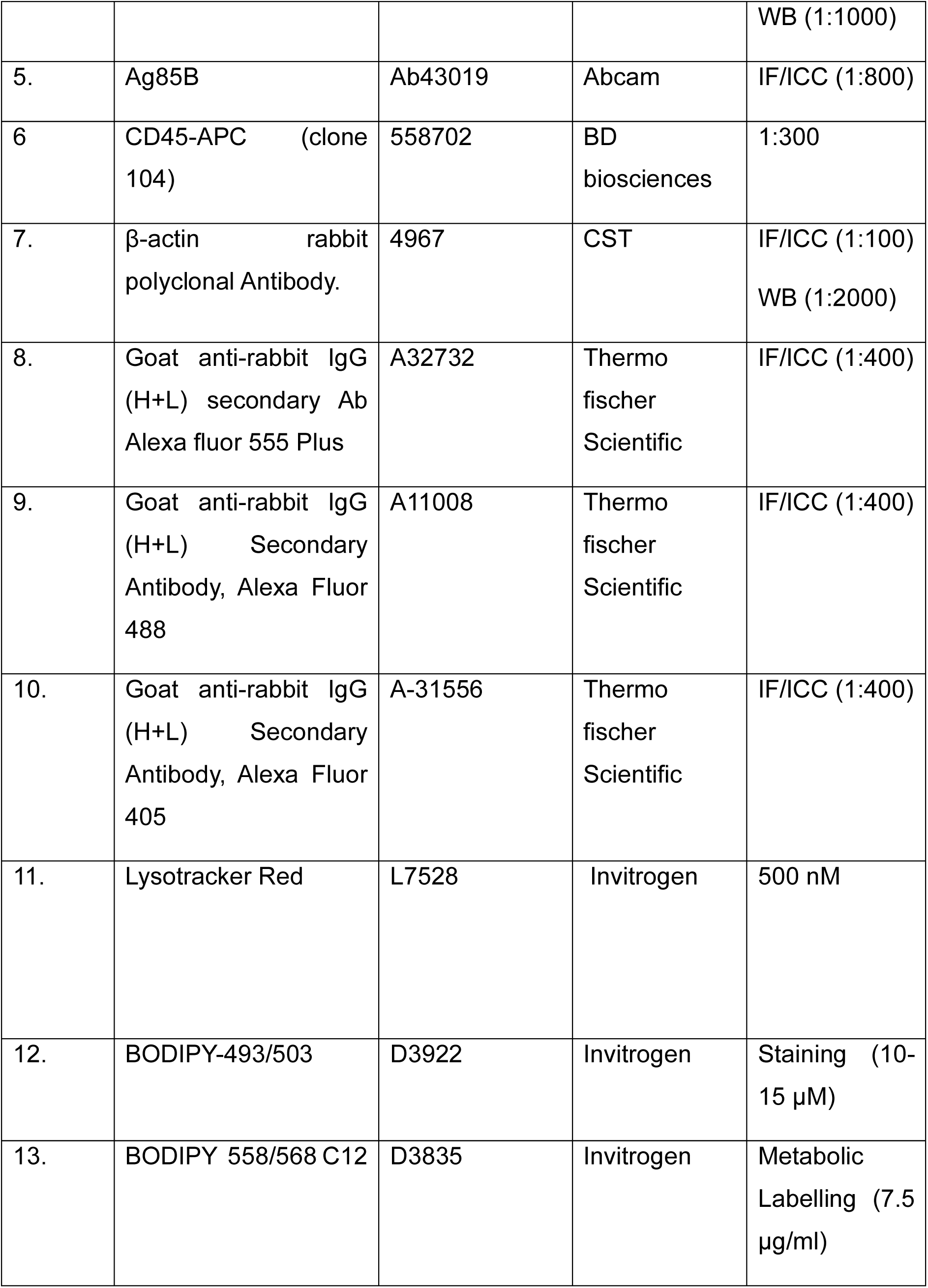
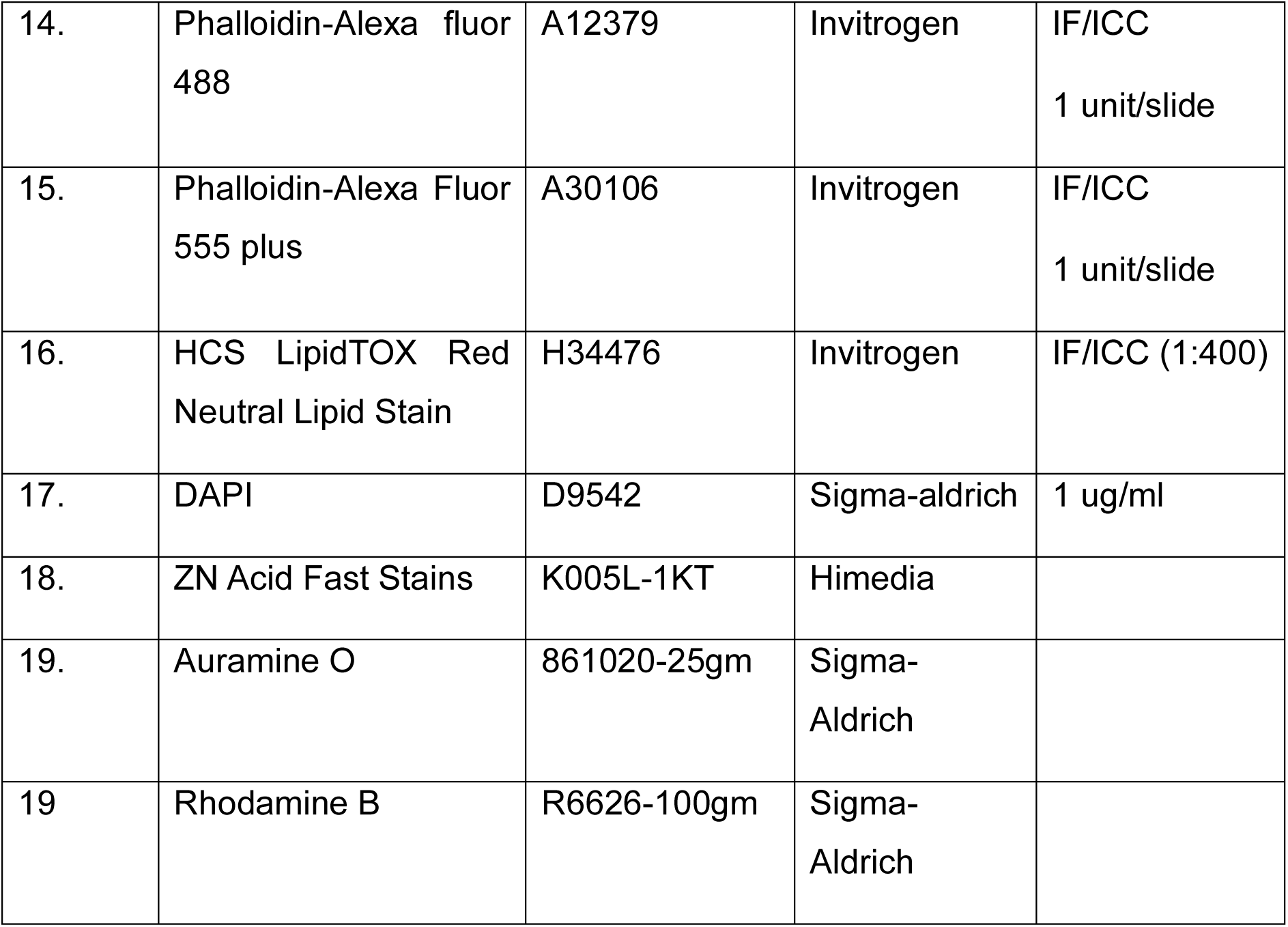

**List of antibodies and dyes used in the study:**
| Serial number | Antibodies and dyes | Catalogue number | Company | Application and dilution |
| --- | --- | --- | --- | --- |
| 1. | Rab5 | 2143T | CST | IF/ICC (1:200) |
| 2. | Rab7 | D9F2 | CST | IF/ICC (1:200) |
| 3. | Cathepsin D | Ab75852 | Abcam | IF/ICC (1:150) |
| 4. | LAMP1 | 9091T | CST | IF/ICC (1:200) |
| 5. | EEA1 | 3288T | CST | IF/ICC (1:200) |
| 6. | ASPGR1 | PA5-32030 | Invitrogen | IF/ICC (1:500) |
| 7. | PPAR $\gamma$ | PA3-831A | Invitrogen | IF/ICC (1:150) |
|  |  |  |  | WB (1:1000) |
| 5. | Ag85B | Ab43019 | Abcam | IF/ICC (1:800) |
| 6 | CD45-APC (clone 104) | 558702 | BD biosciences | 1:300 |
| 7. | β-actin rabbit polyclonal Antibody. | 4967 | CST | IF/ICC (1:100)<br>WB (1:2000) |
| 8. | Goat anti-rabbit IgG (H+L) secondary Ab Alexa fluor 555 Plus | A32732 | Thermo fischer Scientific | IF/ICC (1:400) |
| 9. | Goat anti-rabbit IgG (H+L) Secondary Antibody, Alexa Fluor 488 | A11008 | Thermo fischer Scientific | IF/ICC (1:400) |
| 10. | Goat anti-rabbit IgG (H+L) Secondary Antibody, Alexa Fluor 405 | A-31556 | Thermo fischer Scientific | IF/ICC (1:400) |
| 11. | Lysotracker Red | L7528 | Invitrogen | 500 nM |
| 12. | BODIPY-493/503 | D3922 | Invitrogen | Staining (10-15 μM) |
| 13. | BODIPY 558/568 C12 | D3835 | Invitrogen | Metabolic Labelling (7.5 μg/ml) |
| 14. | Phalloidin-Alexa fluor 488 | A12379 | Invitrogen | IF/ICC<br>1 unit/slide |
| 15. | Phalloidin-Alexa Fluor 555 plus | A30106 | Invitrogen | IF/ICC<br>1 unit/slide |
| 16. | HCS LipidTOX Red Neutral Lipid Stain | H34476 | Invitrogen | IF/ICC (1:400) |
| 17. | DAPI | D9542 | Sigma-aldrich | 1 ug/ml |
| 18. | ZN Acid Fast Stains | K005L-1KT | Himedia |  |
| 19. | Auramine O | 861020-25gm | Sigma-Aldrich |  |
| 19 | Rhodamine B | R6626-100gm | Sigma-Aldrich |  |

#### List of reagents in the study

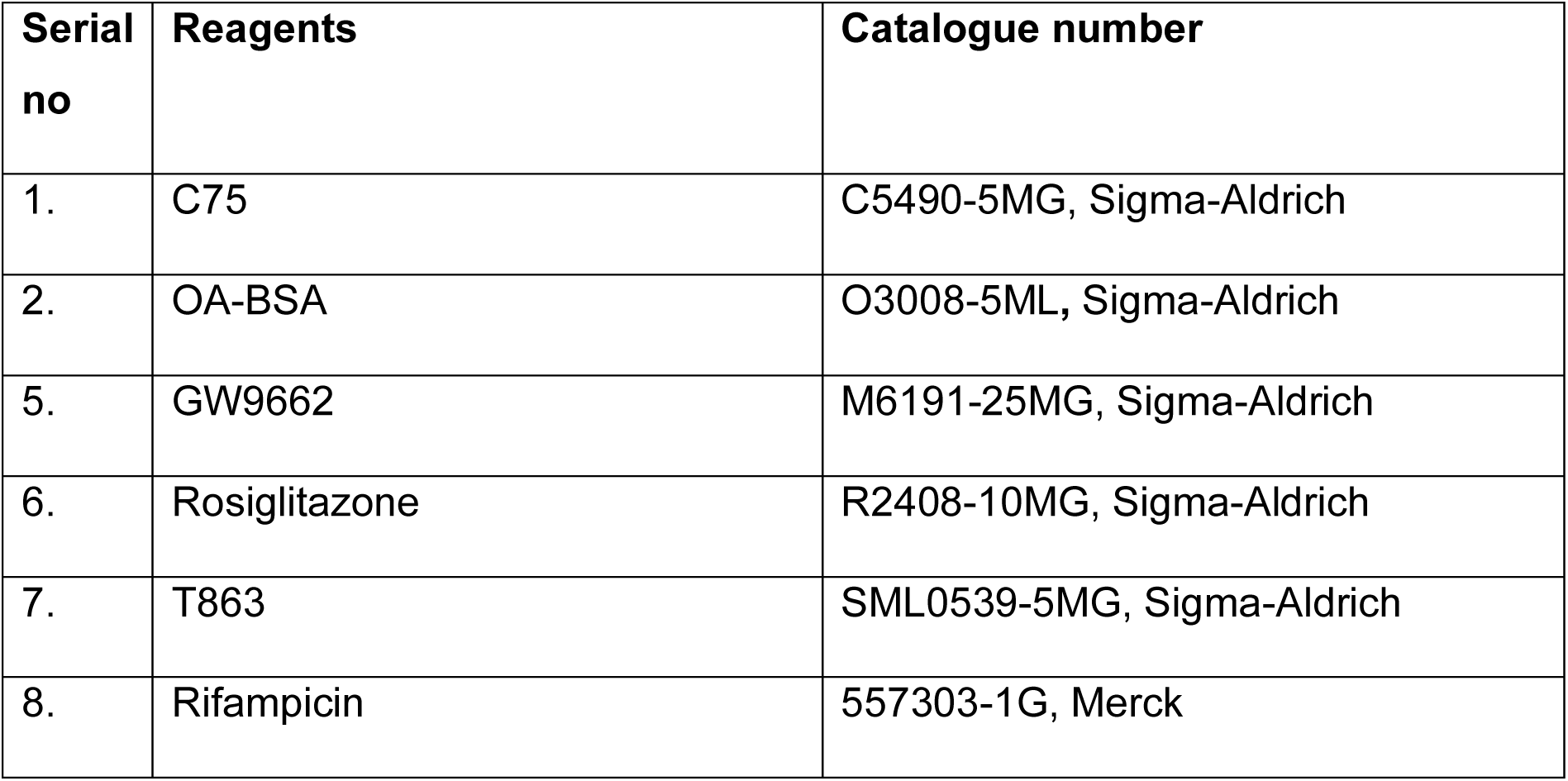
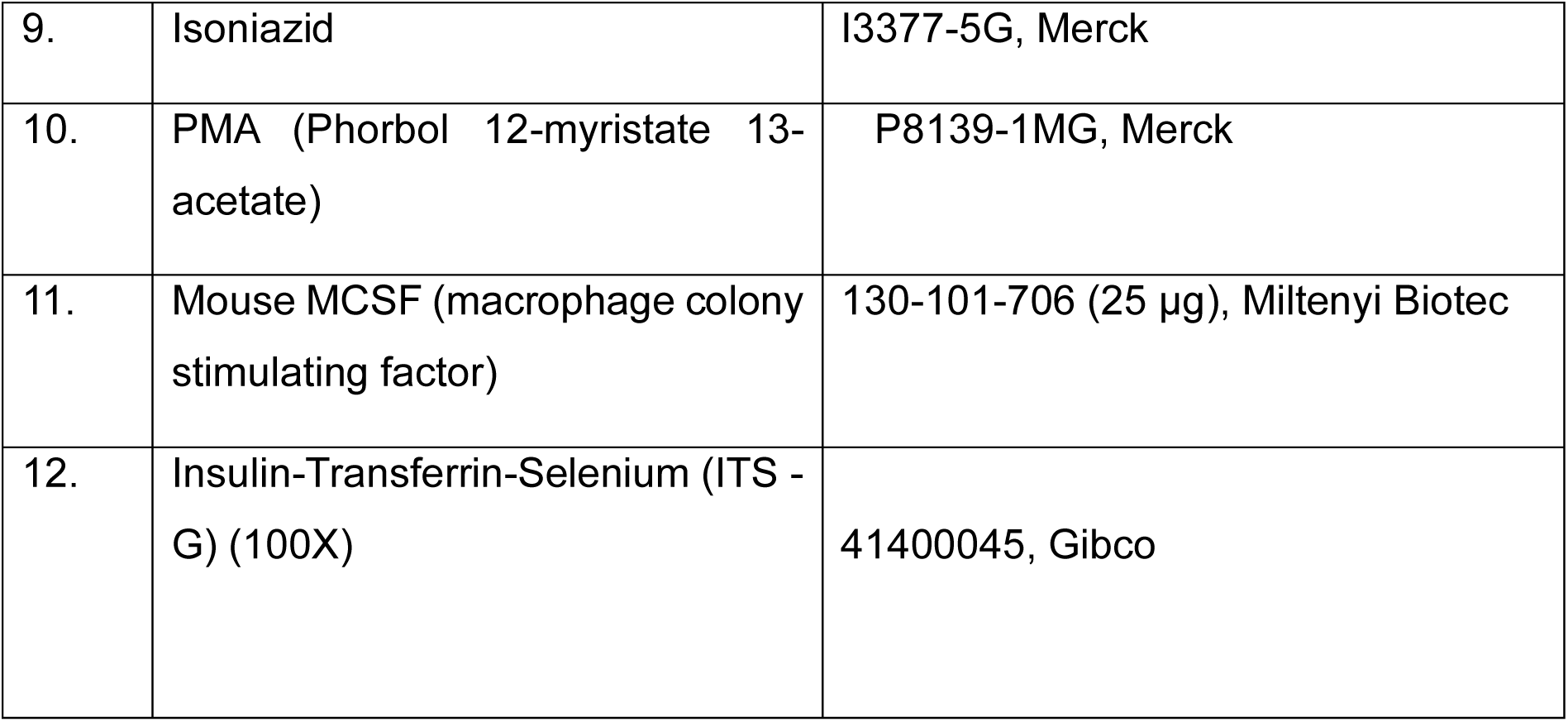

**List of reagents in the study:**
| Serial no | Reagents | Catalogue number |
| --- | --- | --- |
| 1. | C75 | C5490-5MG, Sigma-Aldrich |
| 2. | OA-BSA | O3008-5ML, Sigma-Aldrich |
| 5. | GW9662 | M6191-25MG, Sigma-Aldrich |
| 6. | Rosiglitazone | R2408-10MG, Sigma-Aldrich |
| 7. | T863 | SML0539-5MG, Sigma-Aldrich |
| 8. | Rifampicin | 557303-1G, Merck |
| 9. | Isoniazid | I3377-5G, Merck |
| 10. | PMA (Phorbol 12-myristate 13-acetate) | P8139-1MG, Merck |
| 11. | Mouse MCSF (macrophage colony stimulating factor) | 130-101-706 (25 µg), Miltenyi Biotec |
| 12. | Insulin-Transferrin-Selenium (ITS - G) (100X) | 41400045, Gibco |

#### Sorting of labelled H37Rv cells

The infected hepatocytes were trypsined and washed with 1X PBS followed by passing through 40µm cell strainer to make single cell suspension. After that the cells positive for mCherry signals were sorted using the BD FACS Aria at TACF, ICGEB. Approximately 1 million cells were used for RNA isolation per sample.

#### Primary hepatocyte Isolation

After euthanizing the mice, the peritoneal cavity was opened, and the liver was perfused with 1X HBSS solution till exsanguination was complete. After that collagenase solution (17 mg in 30 ml of 1X HBSS) was passed for digestion. The liver was cut into small pieces and kept in 25 ml of DMEM media followed by lysing the tissues with a glass pestle and sieve. The cells were then passed through a 70 µM cell strainer. 25 ml of Percoll was added to the cells with proper mixing. It was centrifuged at 1000 rpm for 5 minutes. The floating cells were removed, and a brownish layer of pure hepatocytes pelted at the bottom. The cells were counted and plated according to the experimental need on a collagen coated plate.

#### Isolation of Bone marrow derived macrophages (BMDMs) from mouse

BMDMs used for Mtb infection studies were isolated following the published protocol with minor modifications[64]. Briefly, the epiphysis of the femurs and tibia from C57BL/6 mice were cut and the bone marrow were gently flushed into BMDM supplemented with 10% FBS and 1% penicillin streptomycin. The bone marrow cells were centrifuged at 200 X g for 5 minutes at 4° C. The cell pellet was aspirated with 1X PBS and treated with RBC Lysis buffer (for 3 minutes on ice, followed by the addition of complete media. The cells were resuspended in 5 ml of the media and passed through a 70 μm cell strainer. The cells were seeded in complete DMEM supplemented with 10ng/ml of MCSF. On the third day, half of the media volume was replaced by fresh DMEM supplemented with 10ng/ml of MCSF. On the 7^th^ day, the cells were trypsinised, counted and seeded according to experimental requirements.

#### Lipid Extraction protocol and Mass spectrometry

5 million HepG2 cells were infected with Mtb-H37Rv at MOI 10 and kept for 24 hours and 48 hours post-infection. An equal number of uninfected cells were taken. The cells were scrapped, and the procedure of Bligh and Dyer was followed[65]. In brief, the cells were lysed in 1% Triton X-100 after being rinsed twice with 1X PBS. Following lysis, the lysate was vortexed and four volumes of methanol-chloroform (2:1) were added. After that, one volume each of water, chloroform, and 50 mM citric acid was added and vortexed. Following a 10-minute centrifugation at 10,000 rpm at 4° Celsius, the lower organic phase was collected and dried using liquid nitrogen. All semi-quantitative lipid measurements were done using previously reported high-resolution MS/MS methods and chromatographic techniques on an Agilent 6545 QTOF instrument. All sterols were resolved using a Gemini 5U C-18 column (Phenomenex) while DAGs/TAGs were resolved using a Luna 5U C-5 column (Phenomenex) using established solvent systems.

#### Cell lysate preparation and Western blotting

The cells were washed twice with 1X PBS followed by lysis with SDS-RIPA buffer (50 mM Tris-HCl pH 7.5, 150 mM NaCl, 1 mM EDTA, 0.1% SDS, 1% Triton-X 100, 1 mM DTT, 1X Proteinase inhibitor). The cells were incubated with the buffer for 30 minutes in ice followed by vortexing for 5 minutes. The supernantant fraction was collected by centrifuging at 10000 rpm for 20 minutes at 4°C. The protein concentration was determined by bicinchoninic acid (BCA) protein estimation kit (Thermo Fisher Scientific, Waltham, Massachusetts, USA, 23227) following the manufacturer’s protocol. 60-80 ug of protein was resolved in SDS-PAGE followed by transferring onto a PVDF membrane. Blocking was done in 5% skimmed milk in 1X TBST followed by incubation with primary antibodies overnight at 4°C. The membranes were washed 3 times with 1x TBST for 10 minutes each followed by incubation with the HRP-conjugated secondary antibody for 1 hour. Immobilon HRP substrate was used to develop the blots and ImageQuant Chemiluminescent imaging system (LAS 500) was used to acquire the images. Band intensities were measured by using ImageJ.

#### MIC determination of Mtb isolated from primary hepatocytes and primary hepatocytes treated with Rifampicin and isoniazid

Mtb was isolated from primary hepatocytes, primary hepatocytes treated with 0.5μg/ml of rifampicin and primary hepatocytes treated with 0.5μg/ml of isoniazid. These bacteria were grown to a logarithmic phase of OD_600_= 0.6 in Middlebrook 7H9 broth supplemented with 10% OADC, 0.05% glycerol and 0.05% tween 80 under shaking conditions. Single-cell suspension of the Mtb was made as previously described. Two-fold serial dilutions of both rifampicin and isoniazid was prepared in 0.1 ml 7H9-OADC (without Tween 80) (The concentration and the dilution of the drugs have been mentioned in the figure) in 96-well flat bottom microplates. Approximately 5X10^4^ Mtb cells were added to each well in a volume of 0.1 ml. Control wells containing (Mtb only, medium+ inhibitor and only medium) were included in the plate setup. The plate was incubated at 37°C 5 days, followed by the addition of 20 μl of 0.02% resazurin for 24 hours. Visually, minimum inhibitory concentration (MIC) was noted as the lowest drug concentration that prevented the change of colour from blue to pink.

#### qRT-PCR of intracellular Mtb isolated from hepatocytes

HepG2 cells were infected with Mtb at a MOI of 10 and at the respective time points post-infection, the intracellular Mtb was isolated. In all experiments, the cells were treated with amikacin to remove the extracellular bacteria, followed by subsequent washing by 1X PBS. 4M Guanidine isocyanate was added to the flasks in equal amount to the media and the cells were scrapped. The suspension was centrifuged at 3500 rpm for 7 minutes and the pellet was resuspended in 350 μl of the RNA Lysis buffer of MN Kit (740955.250), followed by beat beating. RNA was isolated as per the manufacturer’s protocol. Isolated RNA was treated with Dnase (PGM052, Puregene) for 1 hour to remove genomic DNA contamination. RNA integrity was checked by running it on a 2% agarose gel with RNA loading dye.

1 μg of RNA was reverse transcribed to cDNA using the Takara cDNA synthesis kit (6110A) as per the manufacturer’s protocol. Gene expression analysis by quantitative real-time PCR was performed PowerUp SYBR Green PCR (Thermo fischer scientific, (A25742) master mix in ABI 7500 FAST instrument. 16S rRNA was used as the normalizing control and comparative Ct method was used for quantification.

#### Fluorescent fatty acid labelling of HepG2 and isolation

HepG2 cells were incubated with 7.5 μg/ml of fluorescently tagged fatty acid (BODIPY 558/568 C_12_) for 24 hours. The unincorporated fatty acids were removed by washing it with 1X PBS, three times. After that the cells were treated with DMSO or inhibitors (T863 and C75) for 24 hours, followed by infection with Mtb at a MOI of 10, following the standardized protocol of infection. A separate set of cells before infection were analysed for the effect of the inhibitors on LD formation by microscopy.

After, 48 hours post infection, the cells were washed with 1X PBS and treated with Triton X -100 (0.05% v/v in water), probe sonicated, and Mtb from the cells were isolated by centrifuging at 3500 RPM for 8 minutes. Isolated Mtb cells were washed with 1X PBS and fixed with 4% PFA and stained with DAPI (1mg/ml). The cells were mounted on poly-L-lysine coated slides with antifade agent. The cells were visualised in Zeiss 980 LSM confocal microscope at 100X, magnification.

#### Gene expression studies RNA isolation

Total RNA was isolated from HepG2, PHCs and liver sections using the MN-NucleoSpin RNA isolation kit (740955.250) following the manufacturer’s protocol. For RNA sequencing, an equal number of cells was used during isolation. For Liver tissue, approximately 10mg was tissue was used. The quality of the RNA was verified by running it on a 1.5% agarose gel and by monitoring 260/280 ratios. All the RNA samples were frozen together in -80°C. For RNA sequencing, 3-5 ug of RNA was shipped in sodium acetate buffer (3 M Sodium acetate, pH 5.2) with 100% Ethanol. 4 biological replicates from each time point were sent for sequencing. The RNA samples with RNA integrity number (RIN > 8.5) were used for library preparation.

RNA samples (for four biological replicates) were subjected to pair-end RNA sequencing after rRNA depletion on the Illumina platform Novaseq-6000 at CCMB, Hyderabad, India. Quality control and sequence trimming was performed using fastp (v0.23.2). The trimmed paired-end reads were aligned to the human genome (GRCh38) using the HISAT2 (v2.2.1) pipeline. Reads were assembled into transcripts using StringTie (v2.2.1). Annotation was conducted using aligned sequences and a GTF annotation file. The mapped reads were then used for generating the count table using StringTie (v2.2.1), genes lacking an Ensembl ID annotation were excluded. We arrived at a list of 62694 genes which were used for further analysis (available through GEO accession-GSE256184). The differential count was performed by DEseq2 (R package) using the uninfected samples of respective time points. Pathway enrichment was performed using the GO database and clusters were visualized using the R package ClusterProfiler. Further pathways of interest were analysed by GAGE and visualized using Pathview in KEGG view.

#### cDNA synthesis and RT-qPCR

1 ug of RNA was reverse transcribed to cDNA using the Takara cDNA synthesis kit (6110A) as per the manufacturer’s protocol. Gene expression analysis by quantitative real-time PCR was performed PowerUp SYBR Green PCR (Thermo fischer scientific, (A25742) master mix in ABI 7500 FAST instrument. Beta-actin was used as the normalizing control and comparative Ct method was used for quantification.

### List of primers used in the study

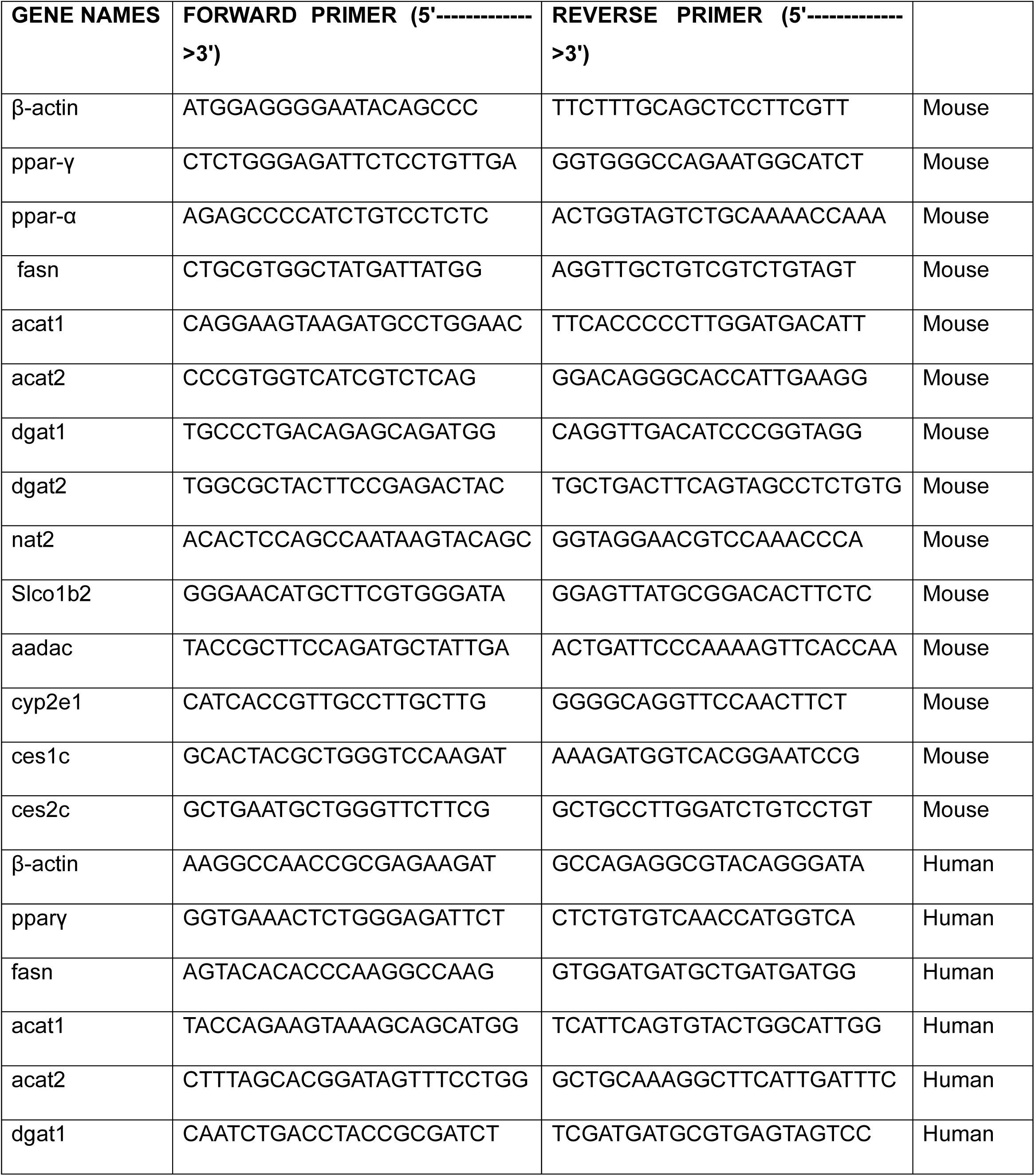
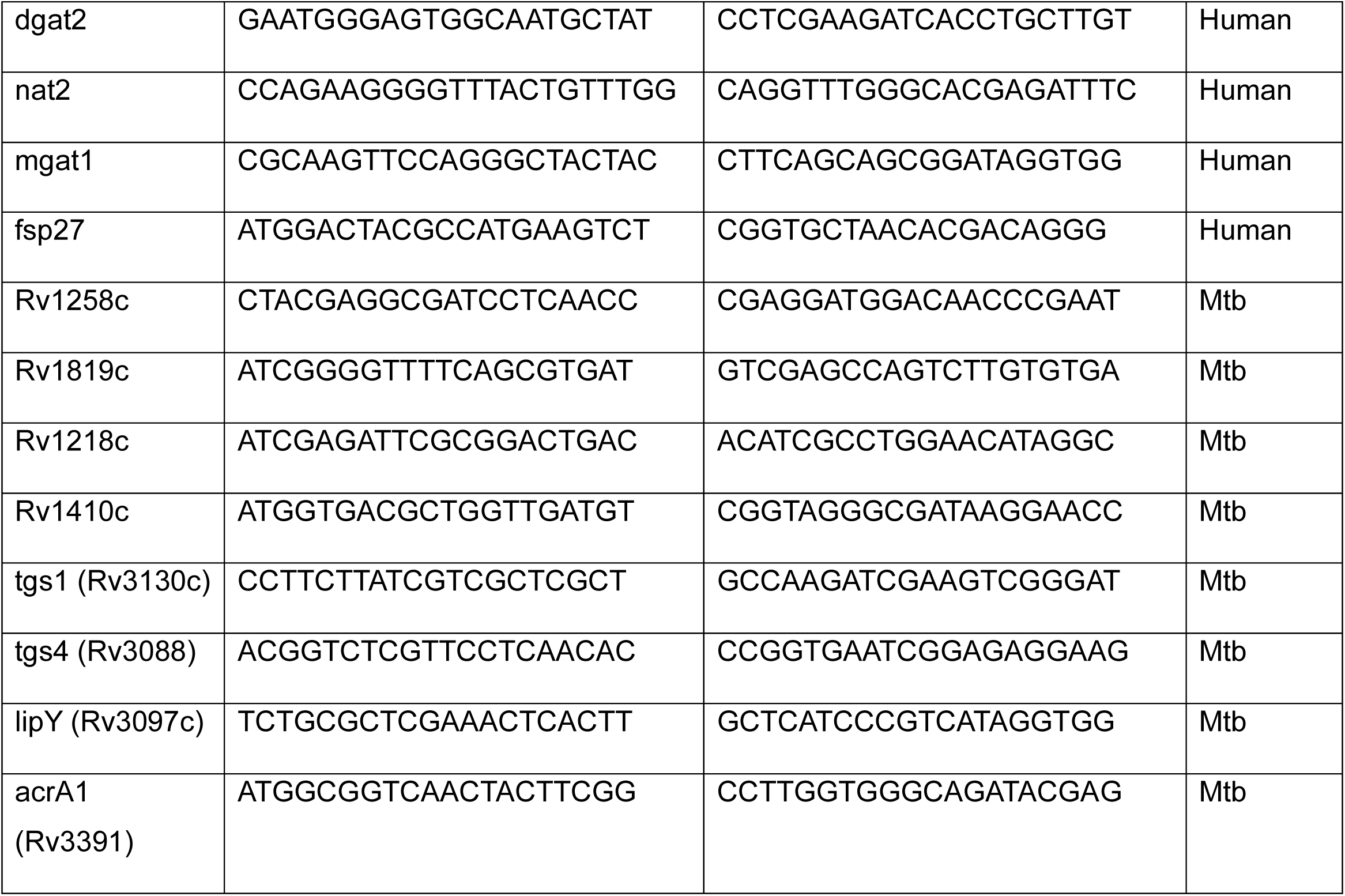

### List of reagents in the study

| Reagent | Company | Catalogue number |
| --- | --- | --- |
| Immobilon Forte Western<br>HRP substrate | Millipore | WBLUF0500 |
| Amersham Hybond Western blotting membranes, PVDF | Merck | GE10600023 |
| PowerUP SYBR | Thermo Fischer Scientific | A25742 |
| Vectashield | Sigma-Aldrich | F6182 |
| Primescript 1<sup>st</sup> strand cDNA synthesis kit | Takara Bio | 6110A |
| NucleoSpin RNA isolation Kit | Macherey-Nagel | 74106 |
| ProLong Gold antifade reagent | Thermo Fischer Scientific | P36930 |
| PolyFreeze Tissue Freezing medium | Sigma- Aldrich | SHH0026 |
| Microscope slides | Himedia | BG005 |
| Poly-Prep slides | Sigma- Aldrich | P0425-72EA |
| 2X RNA loading dye | Thermo Scientific | R0641 |

### Softwares

All the graphs have been generated using GraphPad Prism 10 and Microsoft Excel. The schematics have been drawn with the help of Biorender.com.

## Supporting information

Supplementary figures with legends

## Acknowledgements

We thank Dr. Lakshyaveer Singh, Tuberculosis Aerosol Challenge Facility (TACF), ICGEB for mice experiments. We thank Dr. Neerja Wadhwa for helping in the NII Central Confocal facility, Mr. Birendra Nath Roy for the preparation of the Cryosections, and the NII animal facility for providing us with the animals. We thank the Next Generation Sequencing (NGS) facility at CSIR-CCMB for transcriptomic support.

## Author Contribution

BS and RSG conceptualized the study and analysed the data. BS, DSG, JS, MY, PS, RDS performed the experiments. Human liver sections were stained by SS under the guidance from AK. Transcriptomic analysis was done by JS. Sorting was done by PS and RDS under the guidance from DK. Mass spectrometry analysis was conducted by AC under the supervision of SSK.KM conducted the guinea pig infection experiments. BS and RSG wrote the manuscript. BS, DM, DSG, AK, SSK, DK and RSG reviewed and edited the manuscript. RSG supervised the project.

## Supplementary Figure legends

**Figure S1:**
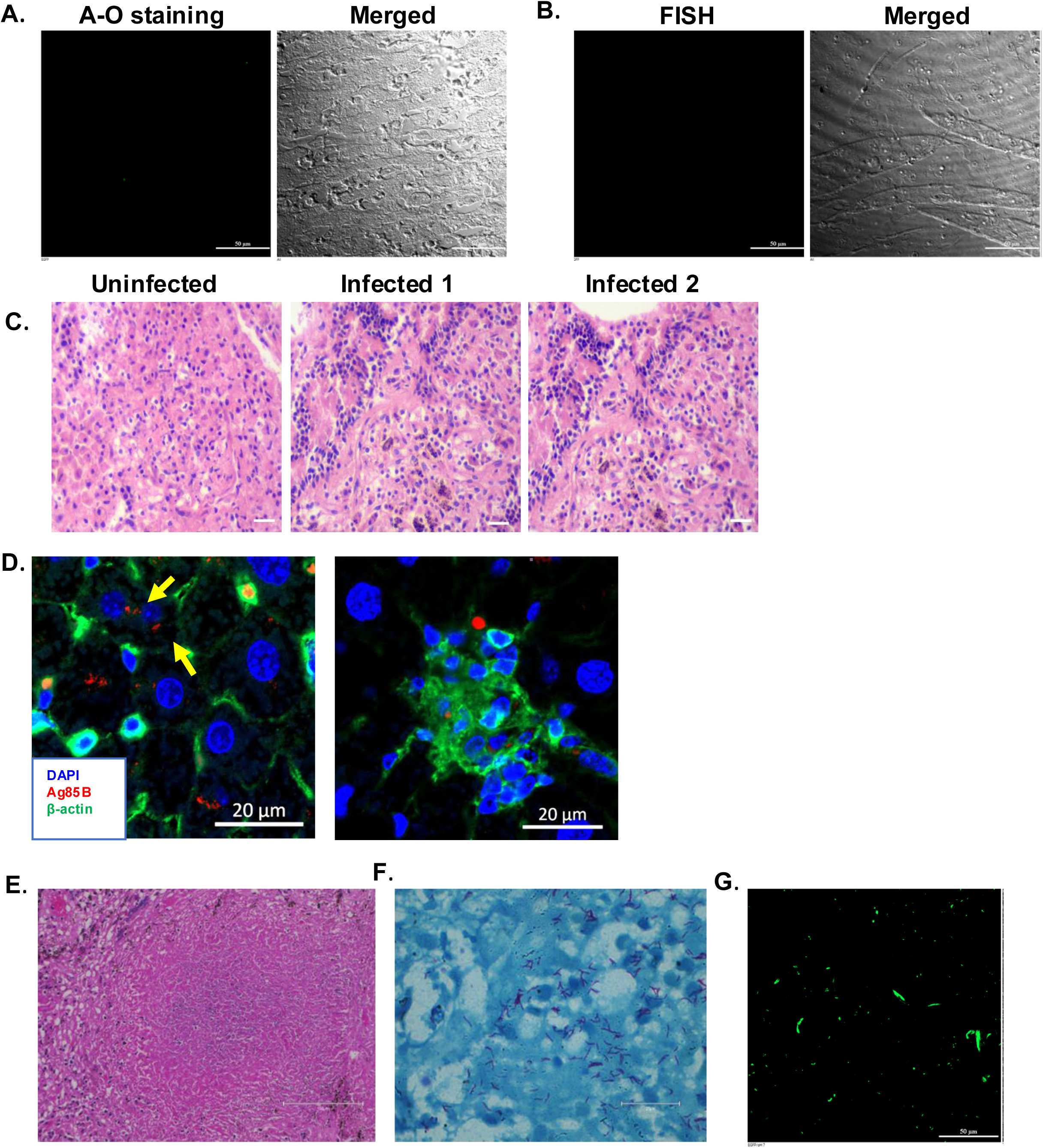
Histopathological changes in the liver biopsy samples of miliaryTB patients: **(A and B):** A-O staining and FISH in the liver biopsy of the uninfected (control) group shows no Mtb-specific signals. **(C).** H & E staining shows enhanced immune cell infiltration in the liver biopsy of the pulmonary TB patients compared to the uninfected control. **(D).** Dual staining of β-actin (green) and Ag85B (red) using respective antibodies shows the presence of Mtb in hepatocytes and other cells in liver biopsy sections (indicated by yellow arrows) **(E).** H&E staining shows distinct granuloma in the lung section of pulmonary TB patient and **(F).** Acid-fast staining in the same lung section shows a high bacterial load **(G).** A-O staining in the same lung section further validates the presence of high Mtb load. Scale bars in A, B and C are 50 µm and in D is 20 µm. and E and F are 150 µm. Data shown in this figure is representative of 5 patients with pulmonary TB.

**Figure S2:**
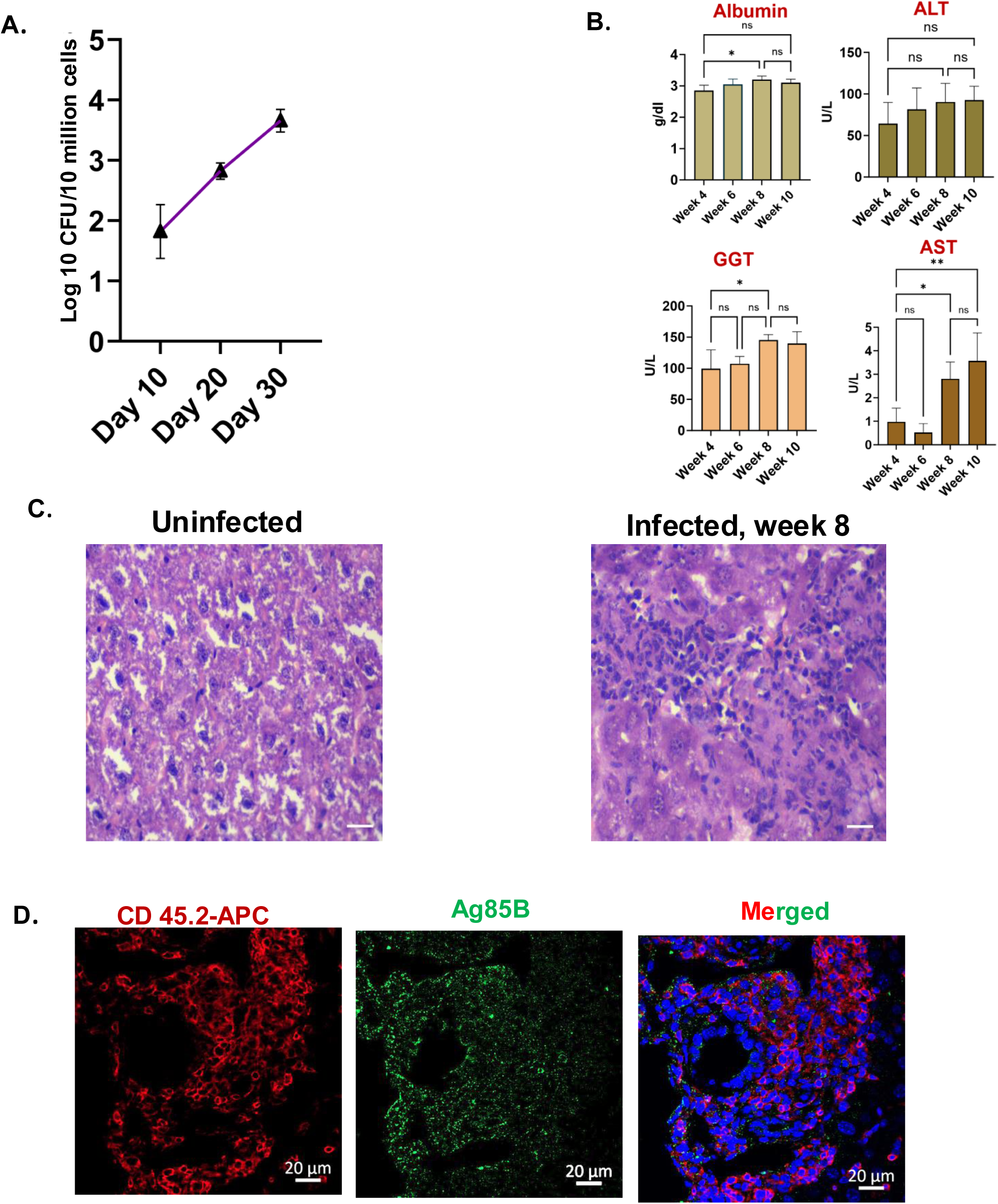
Histopathological changes in the murine liver post Mtb infection: **(A).** 10^5^ number of H37Rv were administered intra-peritoneally (IP) and the bacterial burden was enumerated in primary hepatocytes that were isolated from the infected mice. **(B).** Analysis of the serum parameters of albumin, ALT, AST and GGT in the uninfected and infected mice at the mentioned time points, mice were infected through the aerosol route with 200 CFU. **(C).** Representative H and E-stained liver sections show enhanced immune cell infiltration in Mtb-infected mice. **(D).** Multiplex immunostaining of infected mice liver with CD 45.2 and Ag85B indicates the presence of Ag85B positive signals in CD 45.2 cells. Sample size (n)= 5/6 mice in each group.

**Figure S3:**
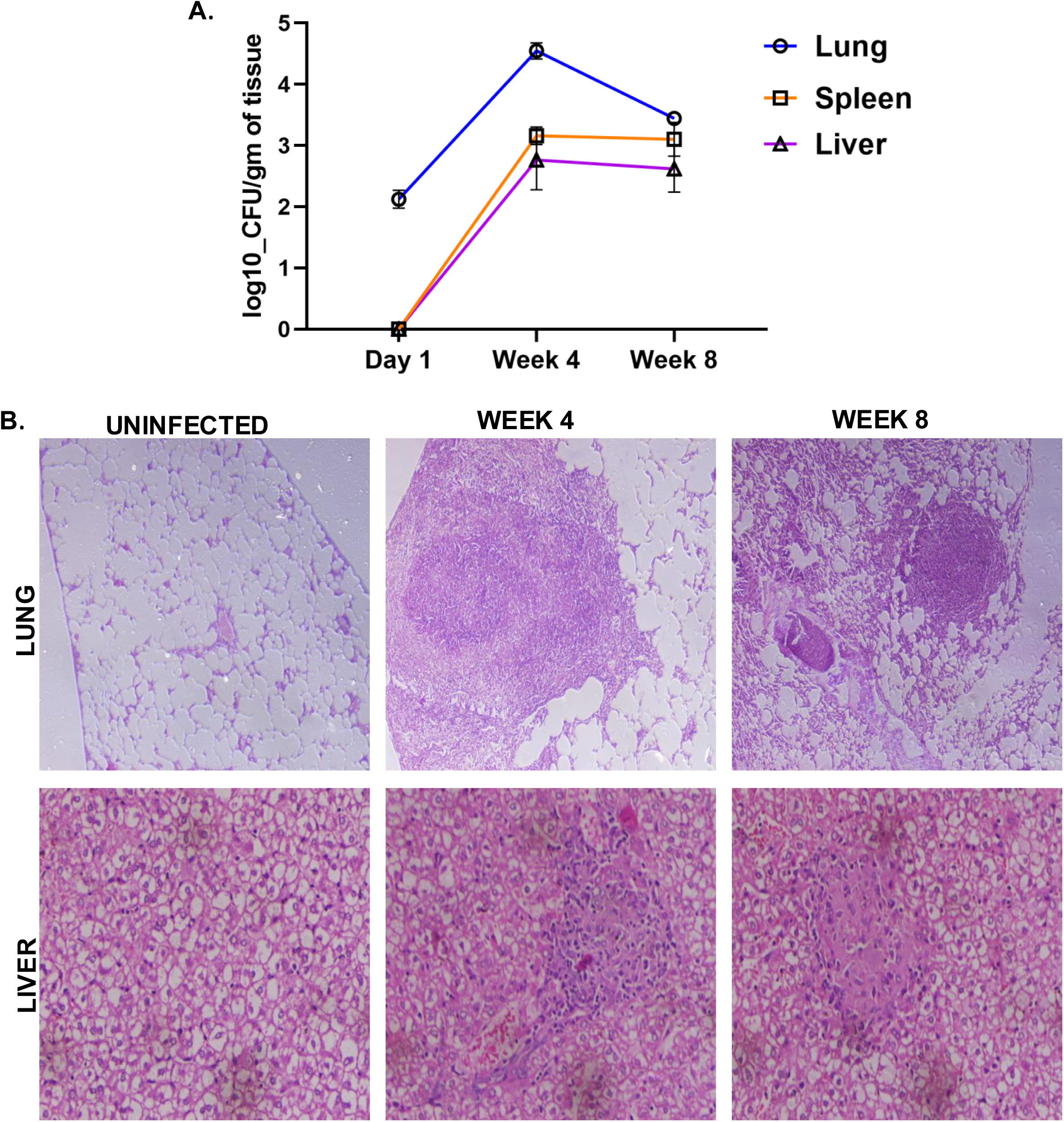
Involvement of the liver in the guinea pig aerosol infection model: **(A).** Guinea pigs were infected with 200 CFU of H37Rv through the aerosol route and the bacterial burden of the lungs, liver and spleen was enumerated at different time points post-infection **(B).** H and E images of the lung and the liver at 4- and 8-weeks post infection showing distinct granulomas in both the organs. Sample size (n)= 3 guinea pigs in each group.

**Figure S4:**
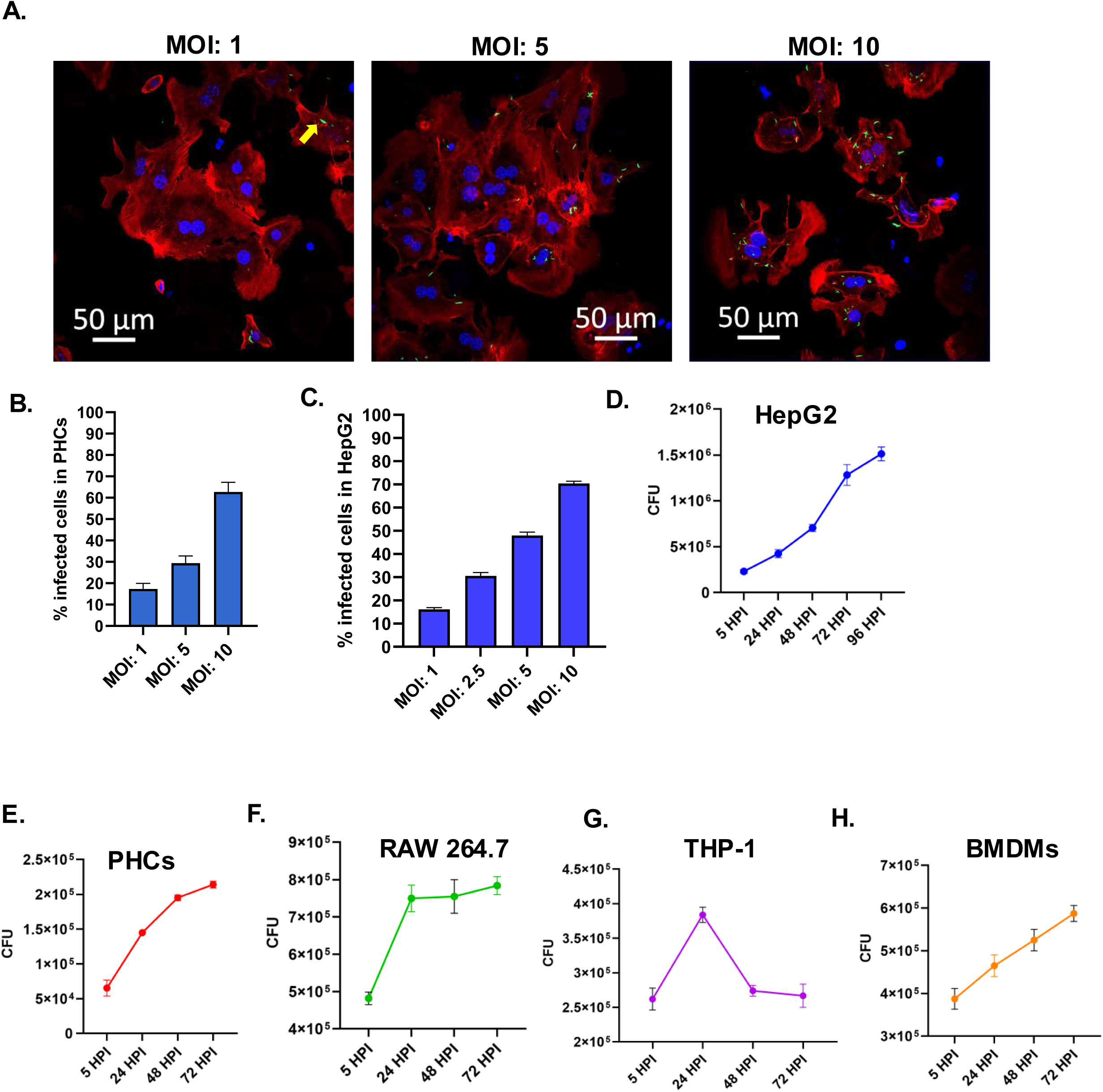
Standardization of multiplicity of infection (MOI) in PHCs and HepG2: **(A).** Confocal microscopic images of Phalloidin Alexa fluor 555 stained PHCs infected with different multiplicity of infection (MOI) of Mtb-H37Rv-GFP. **(B).** Quantification of % infectivity at different MOIs in primary hepatocytes (PHCs) **(C).** Quantification of % infectivity at different MOIs in HepG2. Relative CFU of Mtb in **(D).** HepG2, **(E).** PHCs**, (F).** RAW 264.7, **(G).** THP-1, **(H).** murine BMDMs at the respective time points post-infection.

**Figure S5:**
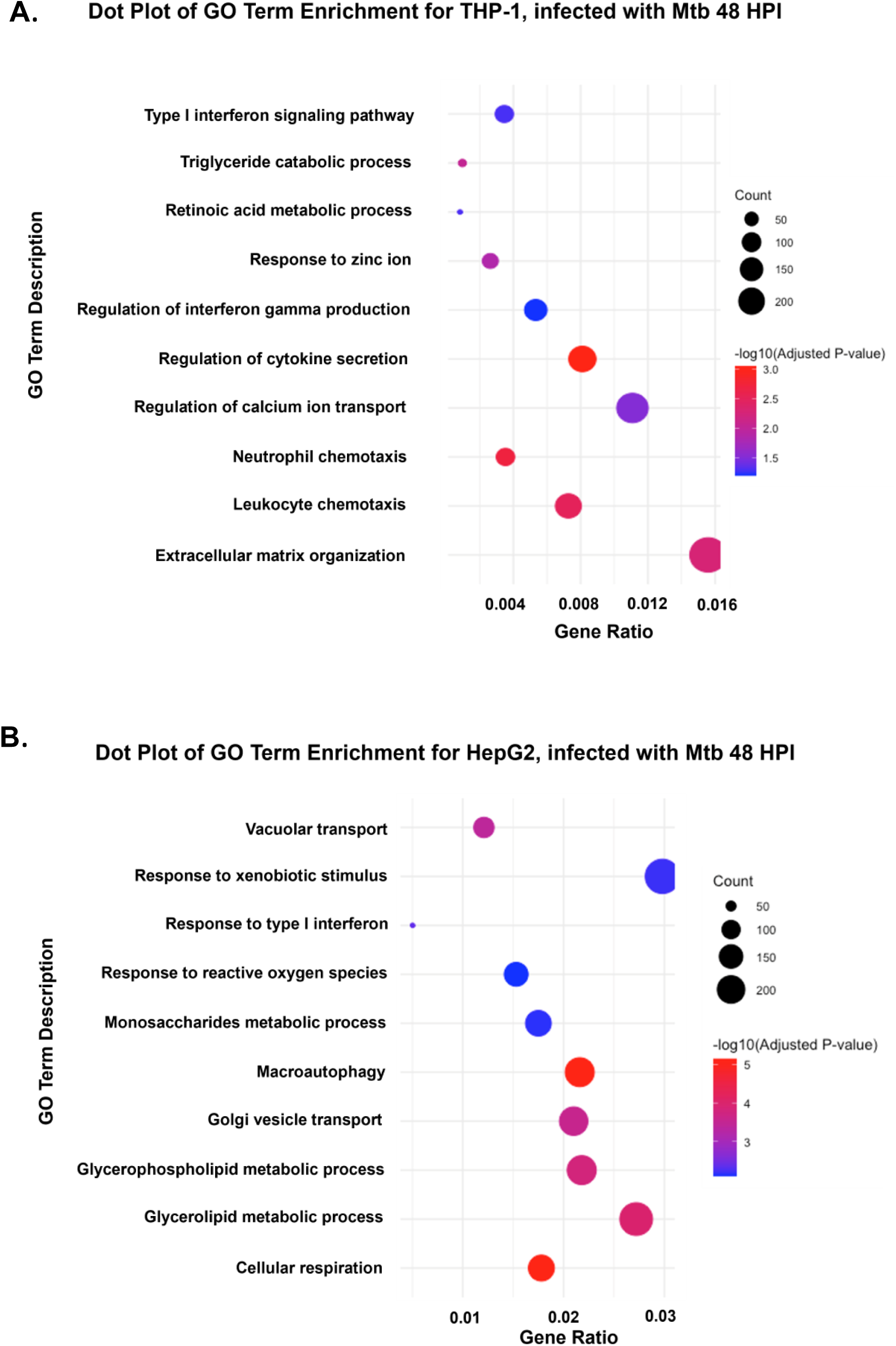
Comparison of significant pathways between Mtb infected THP-1 and HepG2, 48 HPI: (A). GO pathway enrichment analysis was done for DEGs with adjusted p-value < 0.05 at 48 hours post-infection for Mtb-infected THP-1. (B). GO pathway enrichment analysis was done for DEGs with adjusted p-value < 0.05 at 48 hours post infection for Mtb-infected HepG2. The bubble plot depicts the enrichment of pathways on the mentioned time points post-infection, where the coordinate on the x-axis represents the gene ratio, the bubble size represents the gene count and colour represents the p value The bubble plot depicts the enrichment of pathways on the mentioned time points post-infection, where the coordinate on the x-axis represents the gene ratio, the bubble size represents the gene count and colour represents the p-value.

**Figure S6:**
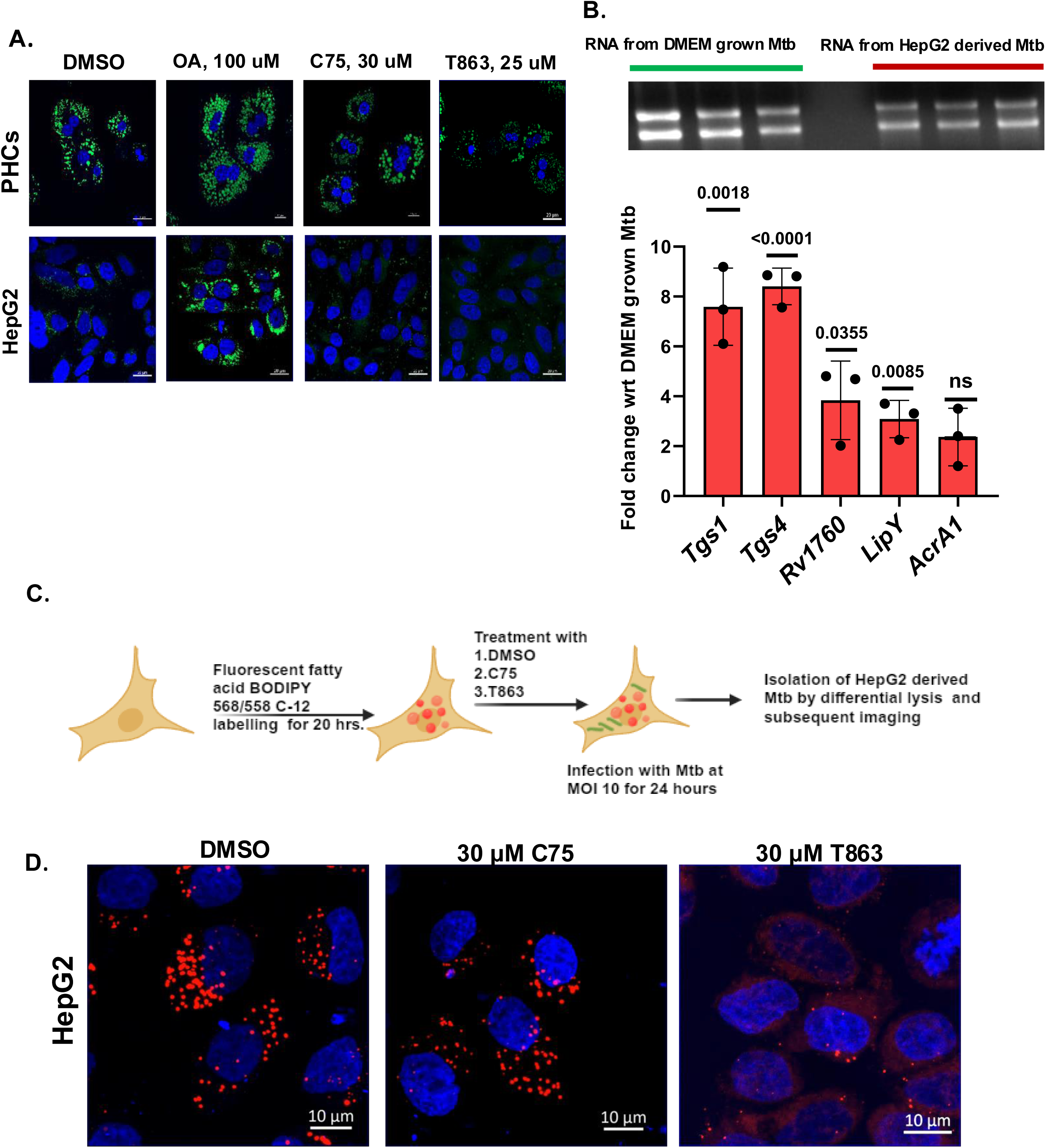
Increased lipid biogenesis provides a growth advantage to Mtb within hepatocytes: (A). Confocal microscopy showing the LDs in both PHCs and HepG2 under inhibitors of fatty acid biosynthesis and TAG biosynthesis. (B). Mtb RNA isolated from DMEM grown Mtb and HepG2-derived Mtb,48 HPI, with the RNA expression pattern of key lipid biogenesis genes and TAG biosynthesis genes in Mtb as determined by qRT PCR, the fold change has been calculated by considering the expression in the DMEM grown Mtb to be 1 (C). Experimental plan for metabolic labelling of Hepg2 and subsequent infection by Mtb (schematic of experimental procedure generated using Biorender.com) (D). Fluorescently labelled fatty acid (BODIPY 558/568 C12) staining in DMSO treated, C75 treated and T863 treated HepG2. Data were analyzed by using the two-tailed unpaired Student’s t-test in B. *p < 0.05, **p < 0.005, ***p < 0.0005, ****p < 0.0001, ns=non-significant.

**Figure S7:**
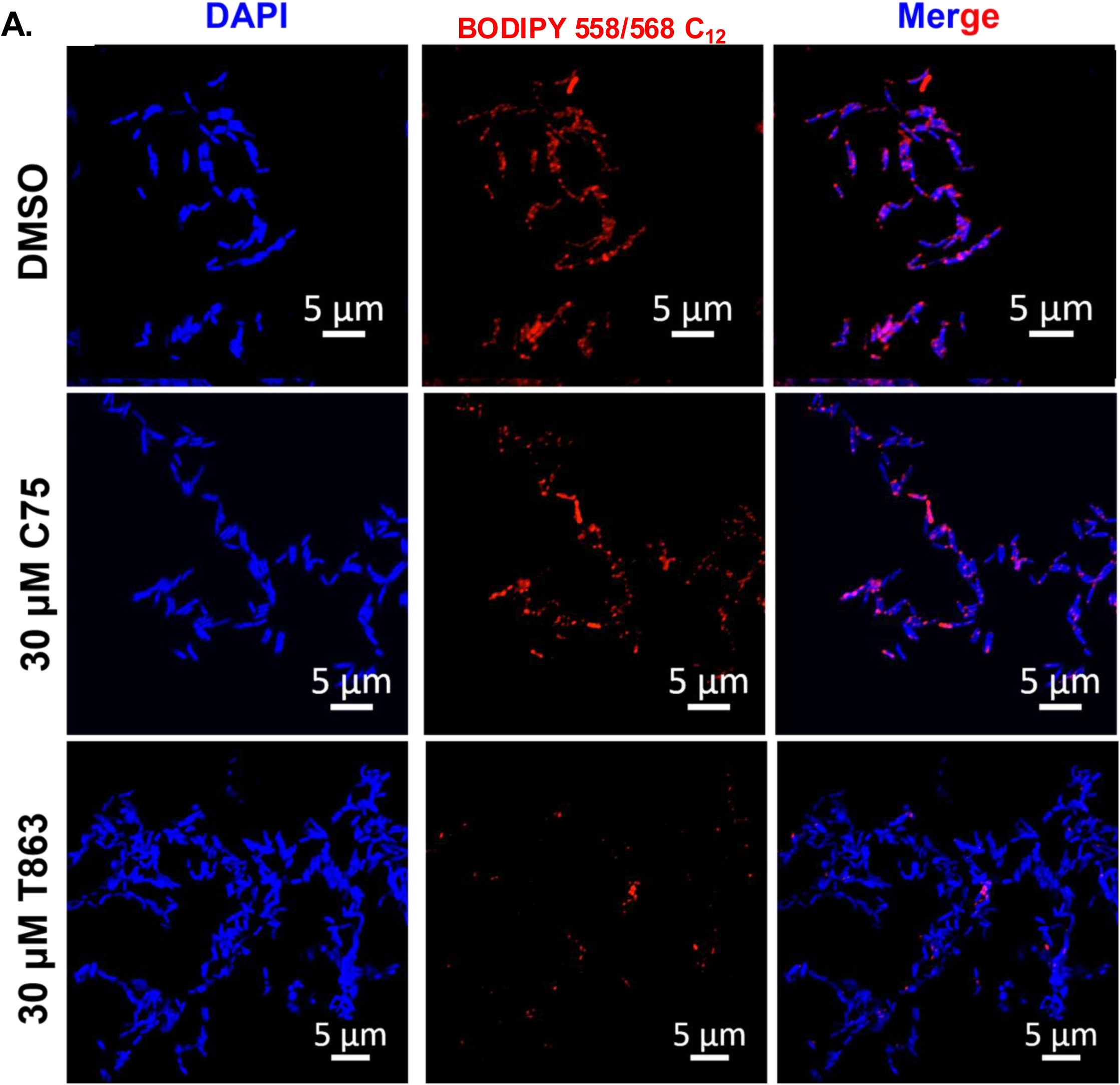
Confocal microscopy images showing the puncta of fluorescently labelled fatty acid (BODIPY558/568 C12) in Mtb isolated from metabolically labeled HepG2: (A). Confocal images showing puncta of labeled fatty acid in Mtb derived from DMSO, C75 and T863 treated metabolically labeled HepG2.

**Figure S8:**
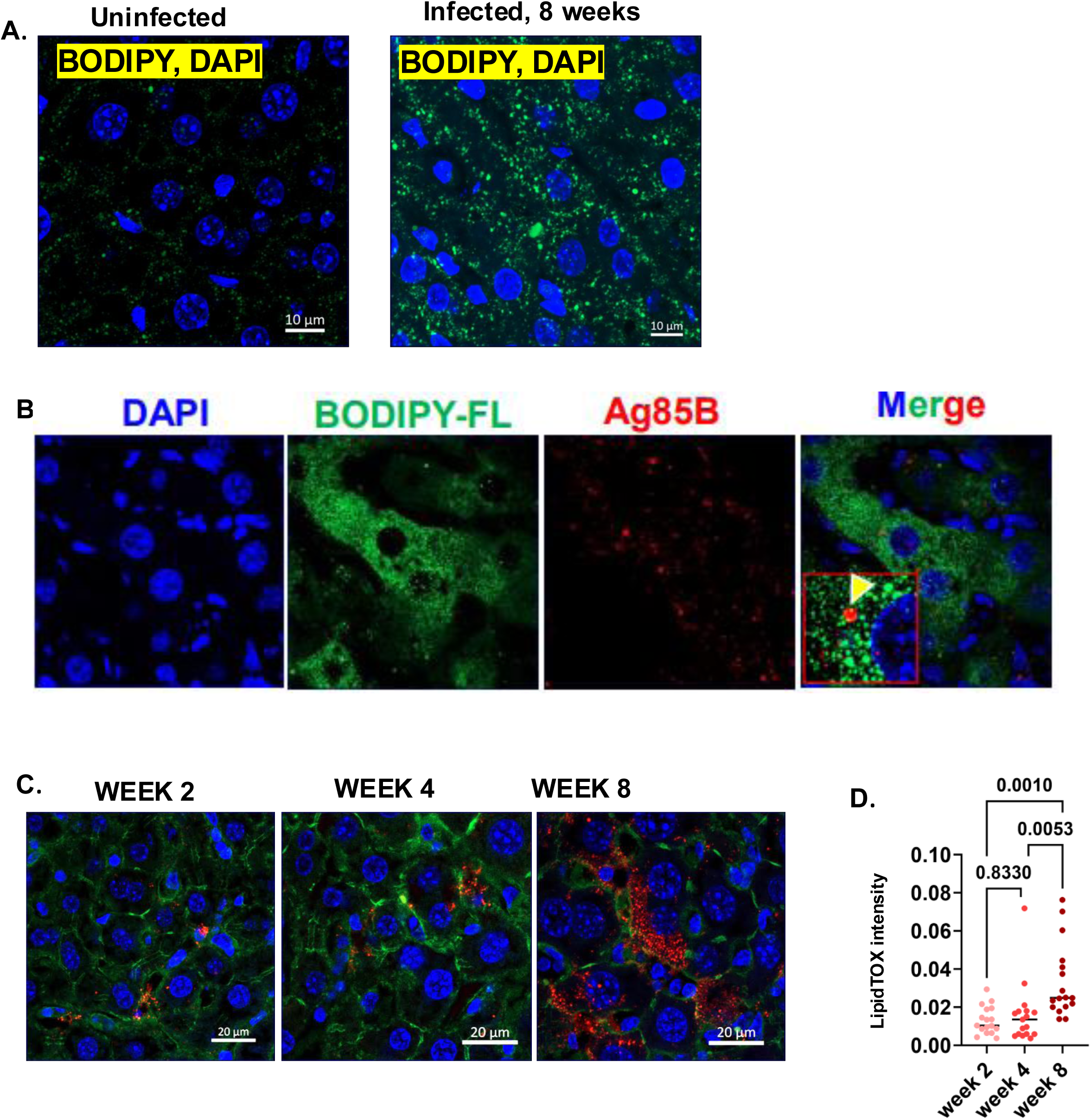
Enhanced lipid accumulation in the liver of Mtb infected mice: (A). BODPIY staining in the liver of the uninfected and 8 weeks post Mtb infected murine liver (B). Dual staining of BODIPY (green) and Ag85B (red) shows Ag85B colocalization with BODIPY (highlighted in the zoomed image). (C). Representative images showing the mean lipidTOX Neutral Red staining in infected liver, 2-, 4- and 8-weeks post infection (D). Plot depicting the mean lipidTOX intensity at the mentioned time points post-infection (normalized to the area. Data were analyzed one-way ANOVA. *p < 0.05, **p < 0.005, ***p < 0.0005, ****p < 0.0001. ns=nonsignificant in I. Representative data from experiments having n=4-6 mice/ group.

**Figure S9:**
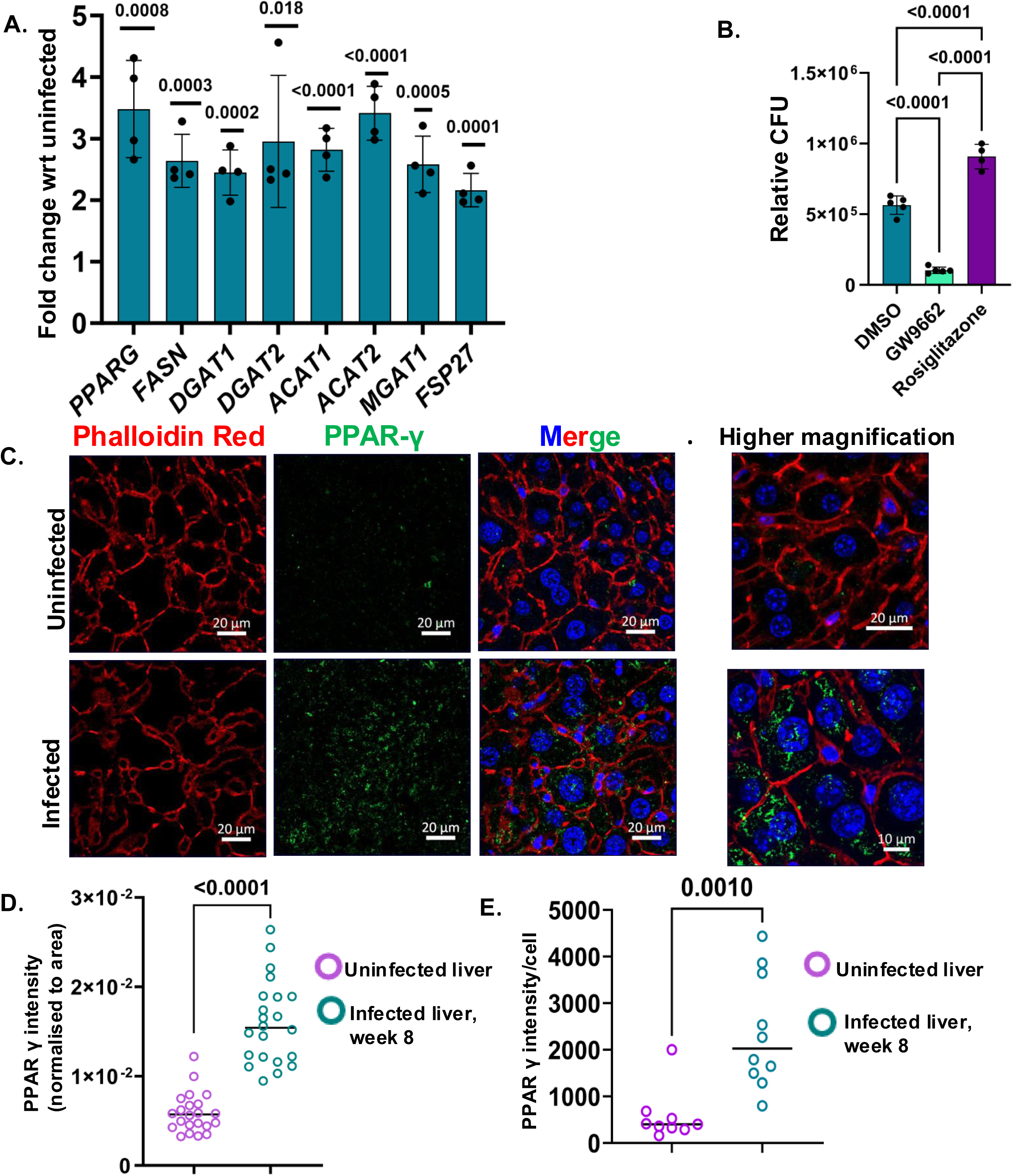
Mtb induced PPARγ expression in infected hepatocytes: (A). mRNA levels of PPARγ and other lipid biosynthetic genes in infected HepG2, the fold change has been calculated by considering the expression in the uninfected cell to be 1 (B). Relative bacterial burden in DMSO, GW9662 and rosiglitazone treated HepG2 infected with Mtb H37Rv. (C). Multiplex immunostaining showed increased expression of PPARγ in the liver of infected mice, 8 weeks post-infection. (D). Bar plot showing PPARγ intensity (normalized to the area) in the uninfected and infected liver, 8 weeks post-infection. (E). Intensity of PPARγ per hepatocyte in uninfected and infected liver, 8-week post-infection. Data were analyzed by using the two-tailed unpaired Student’s t-test in (A, D, and E) and by one-way ANOVA in B. Representative data from n=4 biological replicates *p < 0.05, **p < 0.005, ***p < 0.0005, ****p < 0.0001. ns=non-significant.

**Figure S10:**
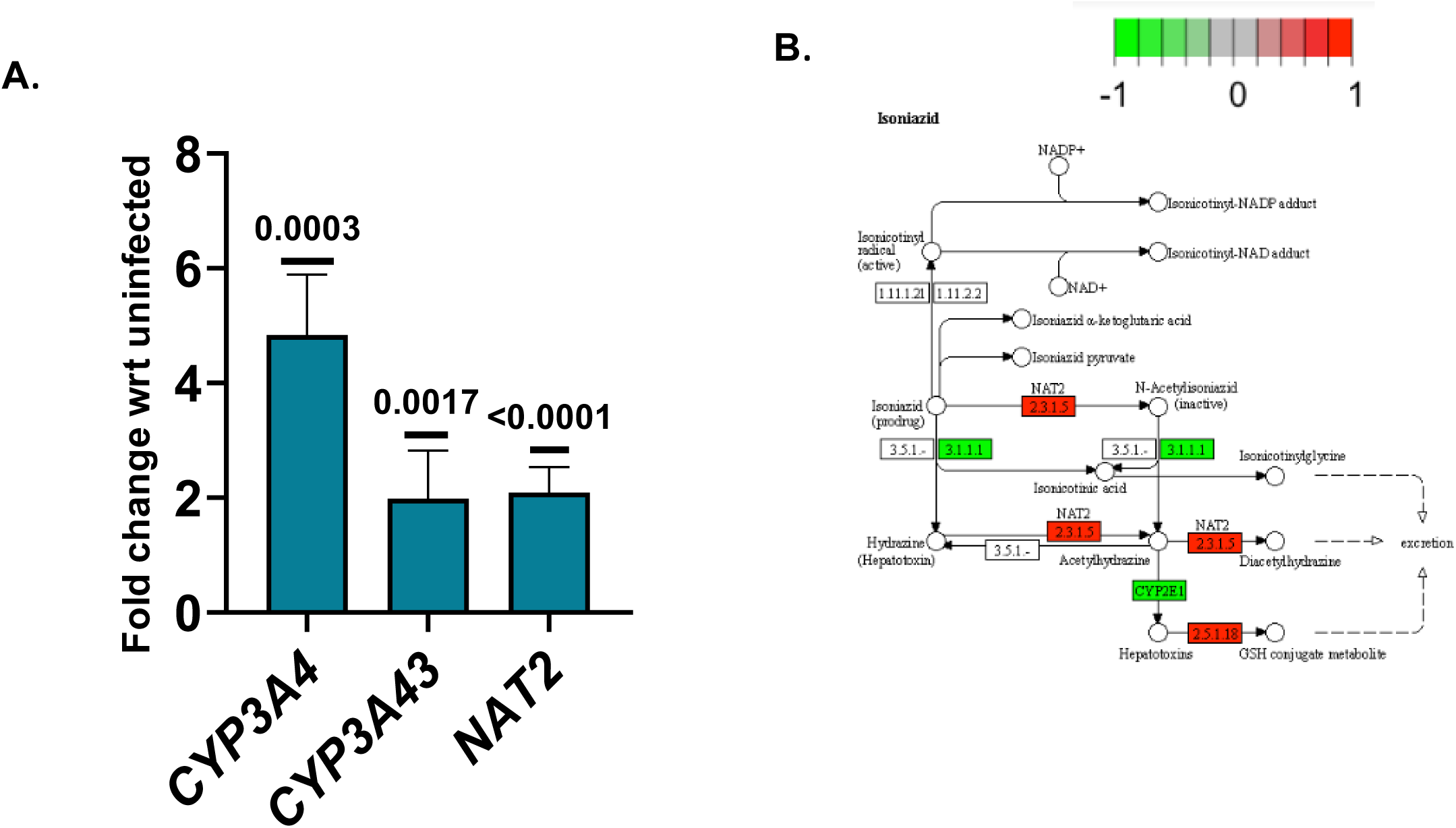
Gene expression analysis of key DMEs related to rifampicin and isoniazid metabolism: (A). Relative expression of CYP3A4, CYP3A43, and NAT2 genes in infected HepG2 post-Mtb infection, the fold change has been calculated by considering the expression in the uninfected cells to be 1. Data were analyzed using the two-tailed unpaired Student’s t test in C. *p < 0.05, **p < 0.005, ***p < 0.0005, ****p < 0.0001, ns=non-significant. (B). KEGG pathway analysis shows the upregulation of NAT2 gene (N-acetyltransferase 2) in Mtb infected HepG2

**Figure S11:**
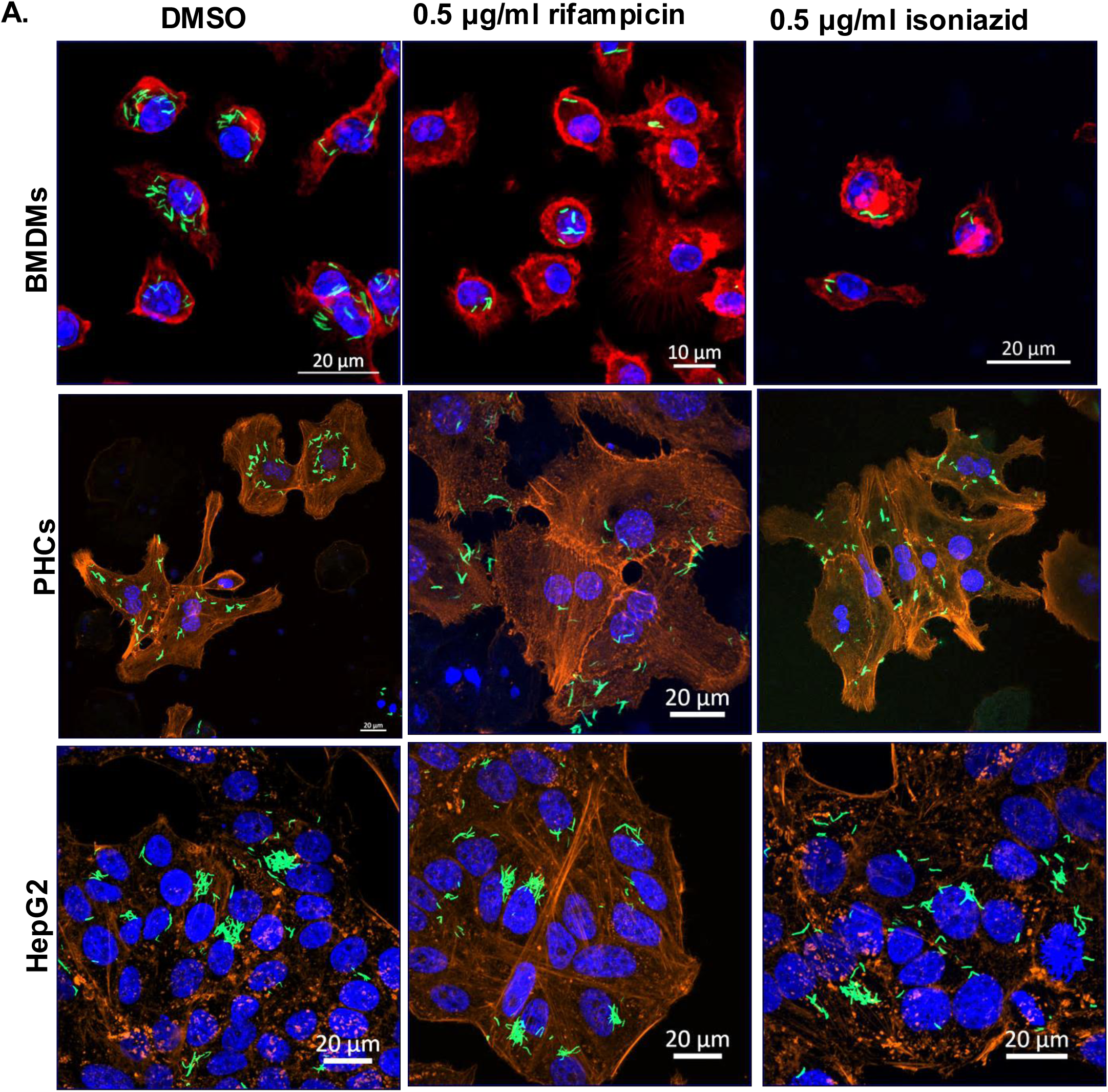
Microscopic images of Mtb infected BMDMs, PHCs and HepG2 treated with rifampicin and isoniazid: (A). Confocal microscopy images of Mtb-H37Rv-GFP infected BMDMs, PHCs, and HepG2. All the cells are treated with DMSO, 0.5 μg/ml rifampicin, and 0.5 μg/ml isoniazid post-infection. The cells were stained with Phalloidin Alexa fluor-555 to demarcate the cell boundary. Representative data from 3 independent experiments. Each dot represents a single field having more than 4 infected cells.20 such fields were analyzed. Figure

**Figure S12:**
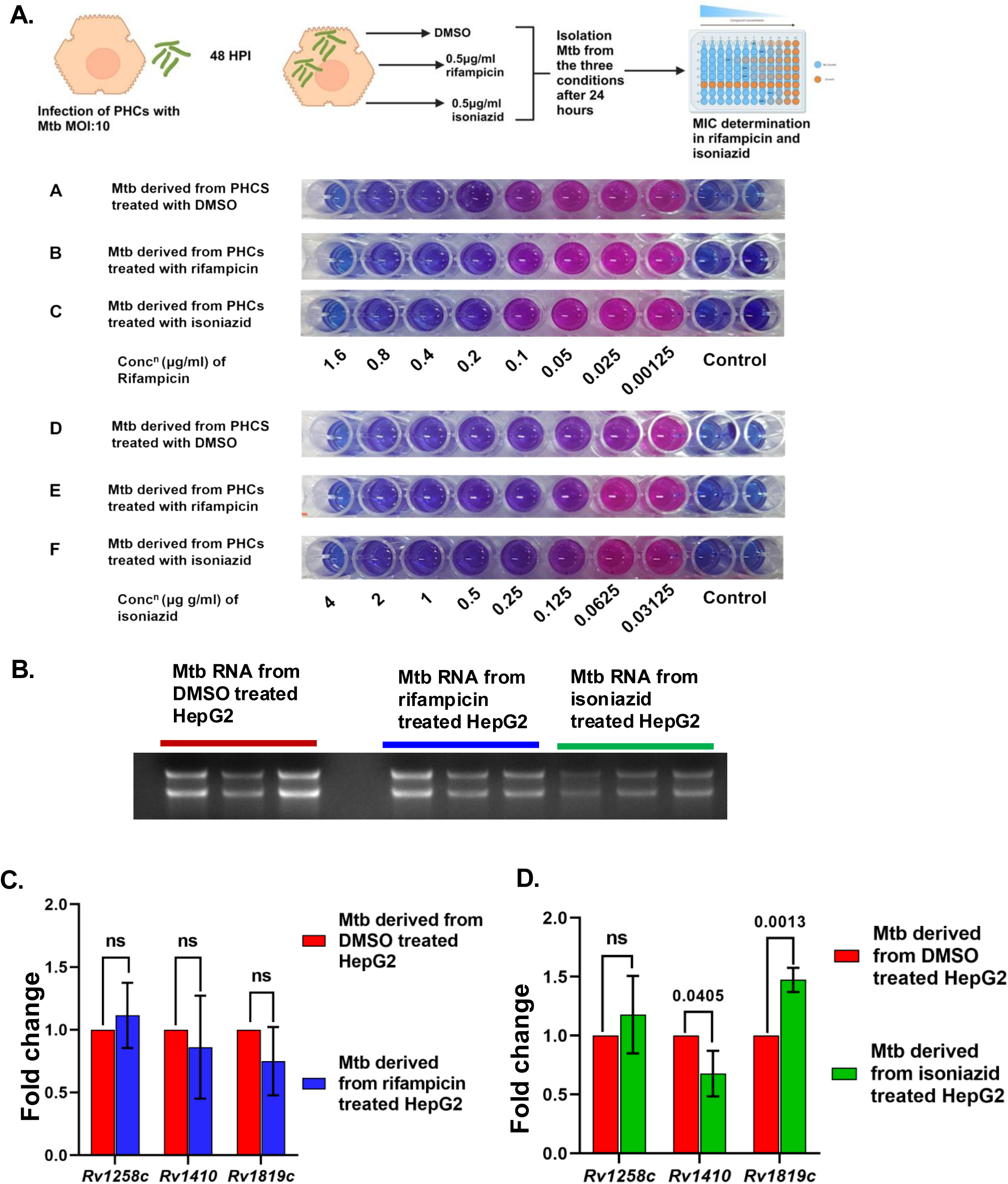
Mtb derived from hepatocytes show no changes in drug efflux pumps and antibiotic sensitivity: (A). Resazurin microtitre assay of the three groups of Mtb in different concentrations of isoniazid and rifampicin (schematic of experimental procedure generated using Biorender.com). (B). Mtb was obtained from DMSO-treated, 0.5 μg/ml rifampicin, and 0.5 μg/ml isoniazid-treated HepG2 by differential lysis and RNA was isolated from those Mtb populations (C and D). qRT-PCR of some of the key efflux pumps genes in these three Mtb groups. Data were analysed using the two-tailed unpaired Student’s t-test in C and D. *p < 0.05, **p < 112 0.005, ***p < 0.0005, ****p < 0.0001, ns=non-significant.

## References

1. *Global Tuberculosis Report* 2023, World health Organisation (WHO).

2. Seung, K.J., S. Keshavjee, and M.L. Rich, Multidrug-Resistant Tuberculosis and Extensively Drug-Resistant Tuberculosis. Cold Spring Harb Perspect Med, 2015. 5(9): p. a017863.

3. Husain, A.A., et al., Current perspective in tuberculosis vaccine development for high TB endemic regions. Tuberculosis (Edinb), 2016. 98: p. 149–58.

4. Eckhardt, M., et al., A systems approach to infectious disease. Nat Rev Genet, 2020. 21(6): p. 339–354.

5. Davis, H.E., et al., Long COVID: major findings, mechanisms and recommendations. Nat Rev Microbiol, 2023. 21(3): p. 133–146.

6. Lippi, G., F. Sanchis-Gomar, and B.M. Henry, COVID-19 and its long-term sequelae: what do we know in 2023? Pol Arch Intern Med, 2023. 133(4).

7. Luies, L. and I. du Preez, The Echo of Pulmonary Tuberculosis: Mechanisms of Clinical Symptoms and Other Disease-Induced Systemic Complications. Clin Microbiol Rev, 2020. 33(4).

8. Loddenkemper, R., M. Lipman, and A. Zumla, Clinical Aspects of Adult Tuberculosis. Cold Spring Harb Perspect Med, 2015. 6(1): p. a017848.

9. Morley, J.E., D.R. Thomas, and M.M. Wilson, Cachexia: pathophysiology and clinical relevance. Am J Clin Nutr, 2006. 83(4): p. 735–43.

10. Chang, S.W., et al., Gut hormones, appetite suppression and cachexia in patients with pulmonary TB. PLoS One, 2013. 8(1): p. e54564.

11. Bisht, M.K., et al., The cause-effect relation of tuberculosis on incidence of diabetes mellitus. Front Cell Infect Microbiol, 2023. 13: p. 1134036.

12. Niazi, A.K. and S. Kalra, Diabetes and tuberculosis: a review of the role of optimal glycemic control. J Diabetes Metab Disord, 2012. 11(1): p. 28.

13. Jensen-Cody, S.O. and M.J. Potthoff, Hepatokines and metabolism: Deciphering communication from the liver. Mol Metab, 2021. 44: p. 101138.

14. Kumar, N.P., et al., Acute Phase Proteins Are Baseline Predictors of Tuberculosis Treatment Failure. Front Immunol, 2021. 12: p. 731878.

15. Quinton, L.J., A.J. Walkey, and J.P. Mizgerd, Integrative Physiology of Pneumonia. Physiol Rev, 2018. 98(3): p. 1417-1464.

16. Shim, D., H. Kim, and S.J. Shin, Mycobacterium tuberculosis Infection-Driven Foamy Macrophages and Their Implications in Tuberculosis Control as Targets for Host-Directed Therapy. Front Immunol, 2020. 11: p. 910.

17. Park, J.H., et al., Understanding Metabolic Regulation Between Host and Pathogens: New Opportunities for the Development of Improved Therapeutic Strategies Against Mycobacterium tuberculosis Infection. Front Cell Infect Microbiol, 2021. 11: p. 635335.

18. Gago, G., L. Diacovich, and H. Gramajo, Lipid metabolism and its implication in mycobacteria-host interaction. Curr Opin Microbiol, 2018. 41: p. 36–42.

19. Wang, F., et al., Organ-organ communication: The liver’s perspective. Theranostics, 2021. 11(7): p. 3317–3330.

20. Alper, C.A., et al., Human C’3: evidence for the liver as the primary site of synthesis. Science, 1969. 163(3864): p. 286-8.

21. Zhou, Z., M.J. Xu, and B. Gao, Hepatocytes: a key cell type for innate immunity. Cell Mol Immunol, 2016. 13(3): p. 301–15.

22. Zhang, D., et al., Important Hormones Regulating Lipid Metabolism. Molecules, 2022. 27(20).

23. Datta, D., et al., Actionable mechanisms of drug tolerance and resistance in Mycobacterium tuberculosis. FEBS J, 2024. 291(20): p. 4433–4452.

24. Deb, C., et al., A novel in vitro multiple-stress dormancy model for Mycobacterium tuberculosis generates a lipid-loaded, drug-tolerant, dormant pathogen. PLoS One, 2009. 4(6): p. e6077.

25. Liu, Y., et al., Immune activation of the host cell induces drug tolerance in Mycobacterium tuberculosis both in vitro and in vivo. J Exp Med, 2016. 213(5): p. 809–25.

26. Samuels, A.N., et al., Understanding the contribution of metabolism to Mycobacterium tuberculosis drug tolerance. Front Cell Infect Microbiol, 2022. 12: p. 958555.

27. Santucci, P., et al., Intracellular localisation of Mycobacterium tuberculosis affects efficacy of the antibiotic pyrazinamide. Nat Commun, 2021. 12(1): p. 3816.

28. Wu, K.C. and C.J. Lin, The regulation of drug-metabolizing enzymes and membrane transporters by inflammation: Evidences in inflammatory diseases and age-related disorders. J Food Drug Anal, 2019. 27(1): p. 48–59.

29. Wu, Z., et al., Diagnosis and treatment of hepatic tuberculosis: report of five cases and review of literature. Int J Clin Exp Med, 2013. 6(9): p. 845–50.

30. Turhan, N., et al., Hepatic granulomas: a clinicopathologic analysis of 86 cases. Pathol Res Pract, 2011. 207(6): p. 359–65.

31. Hickey, A.J., et al., A systematic review of hepatic tuberculosis with considerations in human immunodeficiency virus co-infection. BMC Infect Dis, 2015. 15: p. 209.

32. Celton-Morizur, S., et al., Polyploidy and liver proliferation: central role of insulin signaling. Cell Cycle, 2010. 9(3): p. 460–6.

33. Jain, N., et al., Mesenchymal stem cells offer a drug-tolerant and immune-privileged niche to Mycobacterium tuberculosis. Nat Commun, 2020. 11(1): p. 3062.

34. Lerner, T.R., et al., Mycobacterium tuberculosis cords within lymphatic endothelial cells to evade host immunity. JCI Insight, 2020. 5(10).

35. Kalam, H., M.F. Fontana, and D. Kumar, Alternate splicing of transcripts shape macrophage response to Mycobacterium tuberculosis infection. PLoS Pathog, 2017. 13(3): p. e1006236.

36. Peyron, P., et al., Foamy macrophages from tuberculous patients’ granulomas constitute a nutrient-rich reservoir for M. tuberculosis persistence. PLoS Pathog, 2008. 4(11): p. e1000204.

37. Beigier-Bompadre, M., et al., Mycobacterium tuberculosis infection modulates adipose tissue biology. PLoS Pathog, 2017. 13(10): p. e1006676.

38. Wolk, M. and M. Fedorova, The lipid droplet lipidome. FEBS Lett, 2024. 598(10): p. 1215–1225.

39. Day, N.J., P. Santucci, and M.G. Gutierrez, Host cell environments and antibiotic efficacy in tuberculosis. Trends Microbiol, 2024. 32(3): p. 270–279.

40. Almazroo, O.A., M.K. Miah, and R. Venkataramanan, Drug Metabolism in the Liver. Clin Liver Dis, 2017. 21(1): p. 1–20.

41. Ahmed, S., et al., Pharmacogenomics of Drug Metabolizing Enzymes and Transporters: Relevance to Precision Medicine. Genomics Proteomics Bioinformatics, 2016. 14(5): p. 298–313.

42. Fisher, C.D., et al., Hepatic cytochrome P450 enzyme alterations in humans with progressive stages of nonalcoholic fatty liver disease. Drug Metab Dispos, 2009. 37(10): p. 2087–94.

43. Sotsuka, T., et al., Association of isoniazid-metabolizing enzyme genotypes and isoniazid-induced hepatotoxicity in tuberculosis patients. In Vivo, 2011. 25(5): p. 803–12.

44. Cosma, C.L., D.R. Sherman, and L. Ramakrishnan, The secret lives of the pathogenic mycobacteria. Annu Rev Microbiol, 2003. 57: p. 641–76.

45. Bussi, C. and M.G. Gutierrez, Mycobacterium tuberculosis infection of host cells in space and time. FEMS Microbiol Rev, 2019. 43(4): p. 341–361.

46. Arnett, E., et al., PPARgamma is critical for Mycobacterium tuberculosis induction of Mcl-1 and limitation of human macrophage apoptosis. PLoS Pathog, 2018. 14(6): p. e1007100.

47. Rajaram, M.V., et al., Mycobacterium tuberculosis activates human macrophage peroxisome proliferator-activated receptor gamma linking mannose receptor recognition to regulation of immune responses. J Immunol, 2010. 185(2): p. 929–42.

48. Diaz, A., et al., Studies on the contribution of PPAR Gamma to tuberculosis physiopathology. Front Cell Infect Microbiol, 2023. 13: p. 1067464.

49. Bechmann, L.P., et al., The interaction of hepatic lipid and glucose metabolism in liver diseases. J Hepatol, 2012. 56(4): p. 952–64.

50. Lovewell, R.R., C.M. Sassetti, and B.C. VanderVen, Chewing the fat: lipid metabolism and homeostasis during M. tuberculosis infection. Curr Opin Microbiol, 2016. 29: p. 30–6.

51. Lu, J.F., et al., GDF15 is a major determinant of ketogenic diet-induced weight loss. Cell Metab, 2024. 36(2): p. 454–456.

52. Rey-Bedon, C., et al., CYP450 drug inducibility in NAFLD via an in vitro hepatic model: Understanding drug-drug interactions in the fatty liver. Biomed Pharmacother, 2022. 146: p. 112377.

53. Dookie, N., et al., Evolution of drug resistance in Mycobacterium tuberculosis: a review on the molecular determinants of resistance and implications for personalized care. J Antimicrob Chemother, 2018. 73(5): p. 1138–1151.

54. Shiloh, M.U., Mechanisms of mycobacterial transmission: how does Mycobacterium tuberculosis enter and escape from the human host. Future Microbiol, 2016. 11(12): p. 1503–1506.

55. Coleman, M., et al., Mycobacterium tuberculosis Transmission in High-Incidence Settings-New Paradigms and Insights. Pathogens, 2022. 11(11).

56. Wei, X., et al., The clinical features and prognostic factors of miliary tuberculosis in a high tuberculosis burden area. Ann Med, 2024. 56(1): p. 2356647.

57. McMullan, G.S. and J.H. Lewis, Tuberculosis of the Liver, Biliary Tract, and Pancreas. Microbiol Spectr, 2017. 5(1).

58. Seiler, P., et al., Limited mycobacterial infection of the liver as a consequence of its microanatomical structure causing restriction of mycobacterial growth to professional phagocytes. Infect Immun, 2001. 69(12): p. 7922–6.

59. Hess, S., et al., In vivo partial reprogramming by bacteria promotes adult liver organ growth without fibrosis and tumorigenesis. Cell Rep Med, 2022. 3(11): p. 100820.

60. Barreto, E.A., et al., COVID-19-related hyperglycemia is associated with infection of hepatocytes and stimulation of gluconeogenesis. Proc Natl Acad Sci U S A, 2023. 120(21): p. e2217119120.

61. Mercado-Gomez, M., et al., The spike of SARS-CoV-2 promotes metabolic rewiring in hepatocytes. Commun Biol, 2022. 5(1): p. 827.

62. Hazlehurst, J.M., et al., Non-alcoholic fatty liver disease and diabetes. Metabolism, 2016. 65(8): p. 1096–108.

63. Dharmalingam, M. and P.G. Yamasandhi, Nonalcoholic Fatty Liver Disease and Type 2 Diabetes Mellitus. Indian J Endocrinol Metab, 2018. 22(3): p. 421–428.

64. Toda, G., et al., Preparation and culture of bone marrow-derived macrophages from mice for functional analysis. STAR Protoc, 2021. 2(1): p. 100246.

65. Bligh, E.G. and W.J. Dyer, A rapid method of total lipid extraction and purification. Can J Biochem Physiol, 1959. 37(8): p. 911–7.

