## Supplementary figures with legends for "PPARγ mediated enhanced lipid biogenesis fuels *Mycobacterium tuberculosis* growth in a drug-tolerant hepatocyte environment"

Supplementary data for the manuscript **‘’PPARγ mediated lipid biogenesis fuels *Mycobacterium tuberculosis* growth in a drug-tolerant hepatocyte environment”** **(REVISED)**

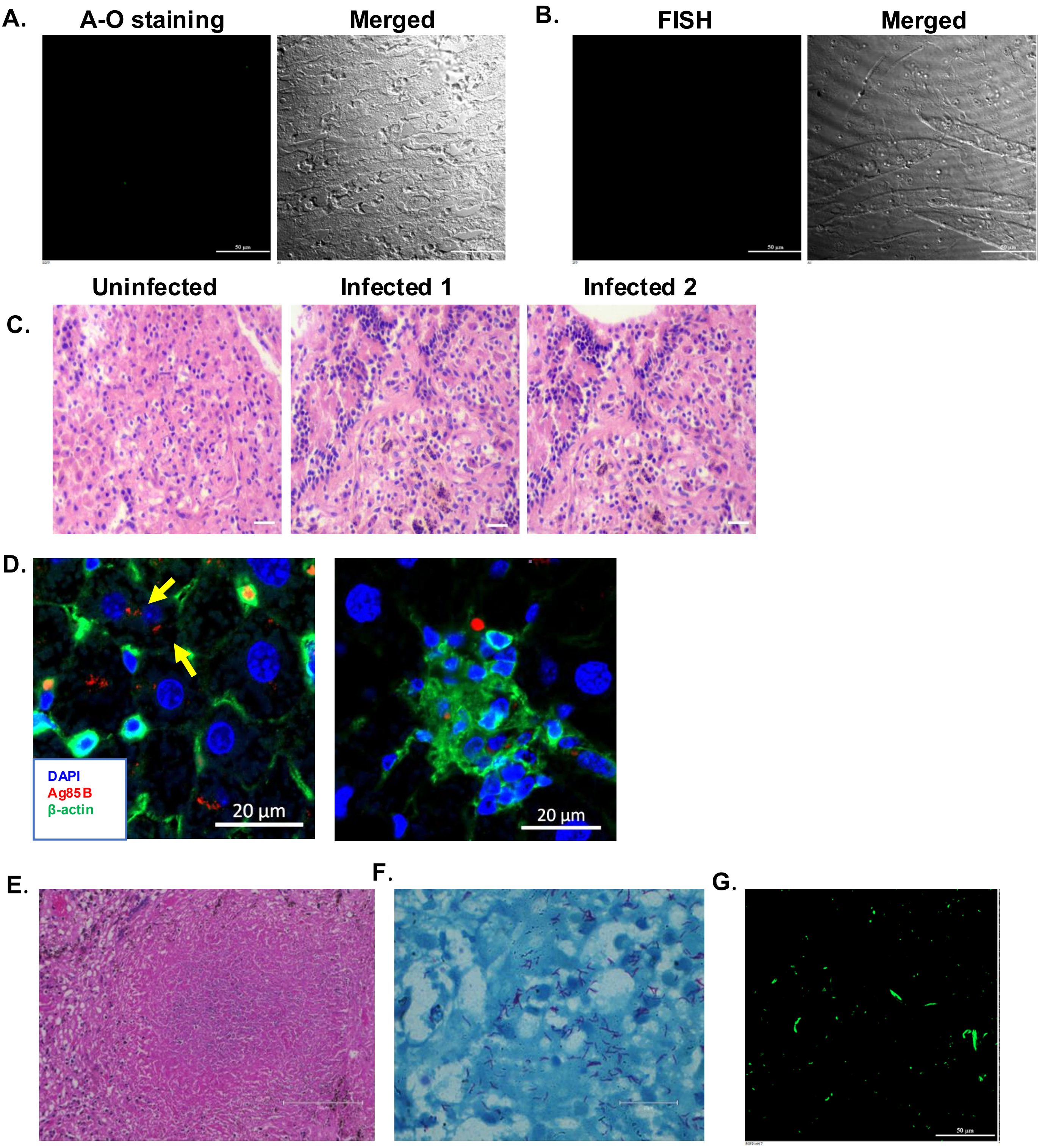


**Figure 1, Supplementary**

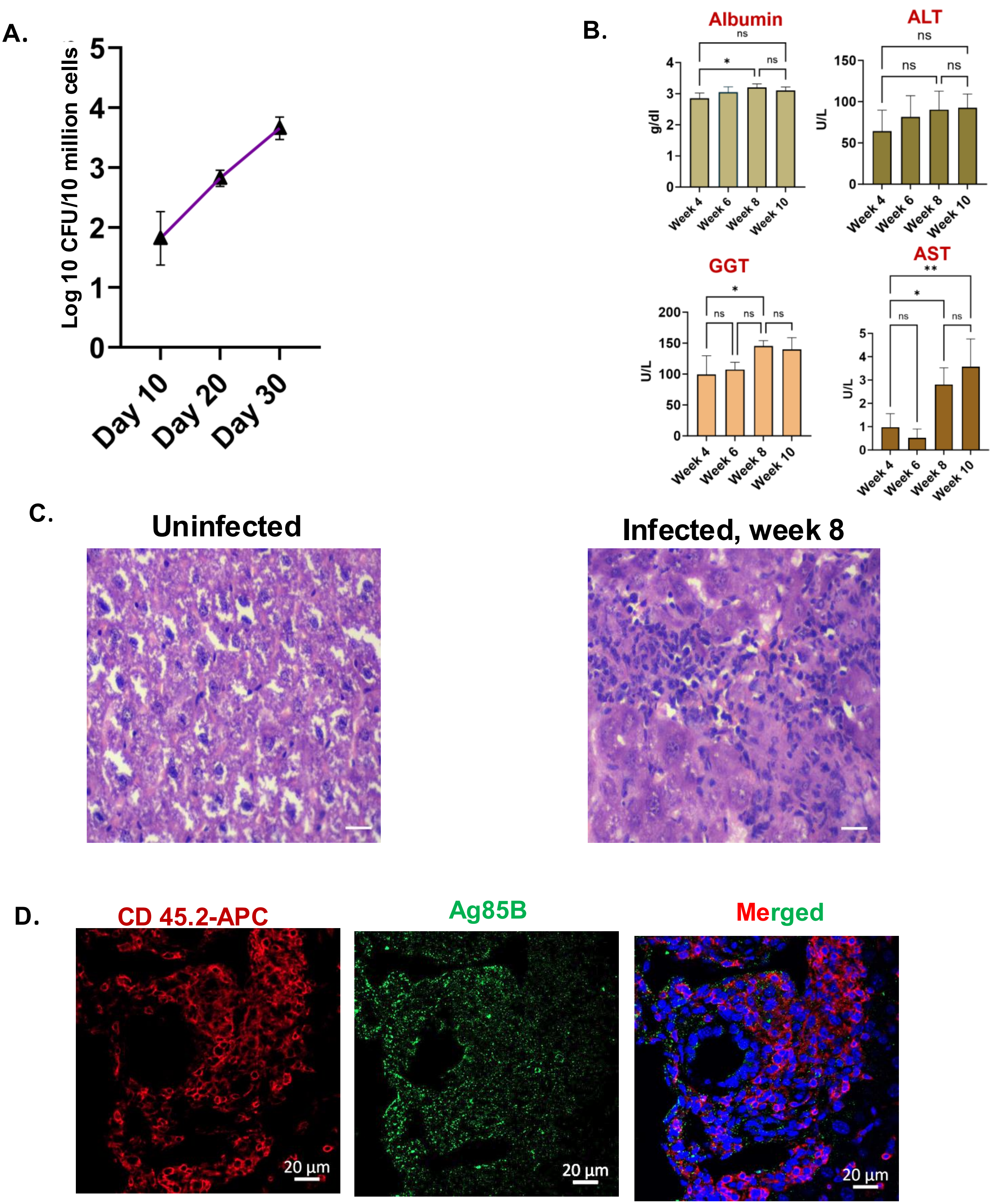


Figure 2, Supplementary

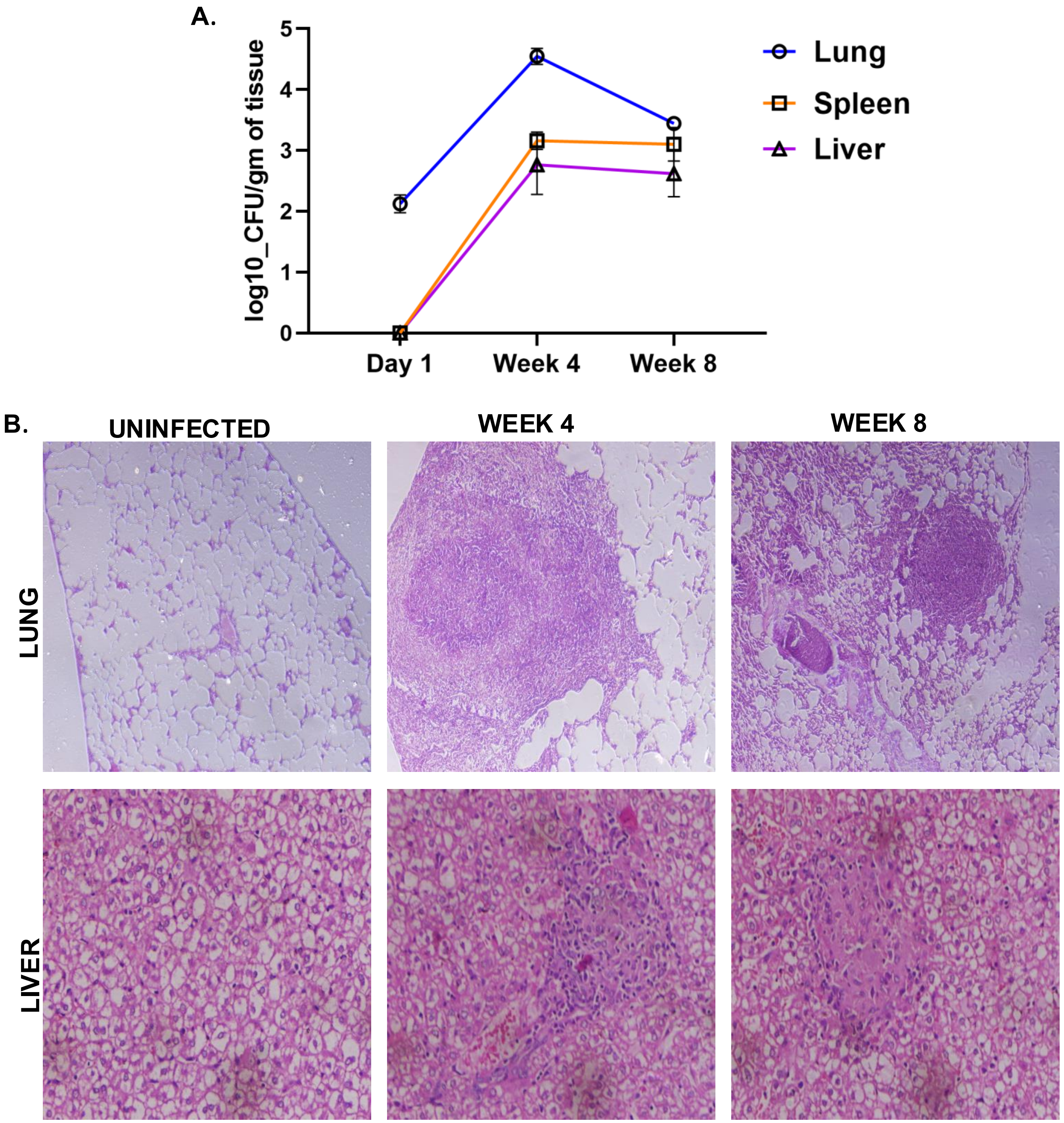


Figure 3, Supplementary

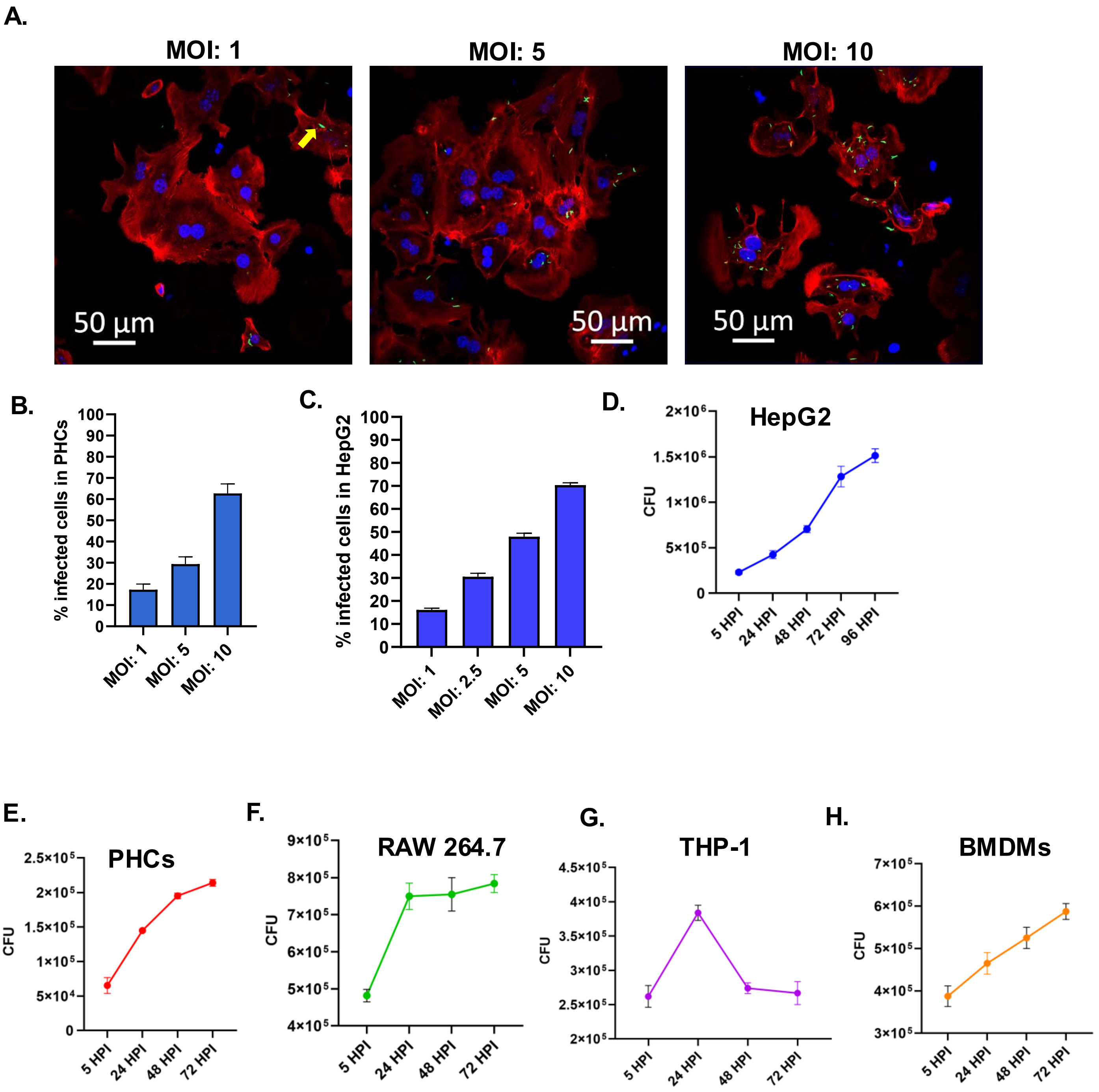


Figure 4, Supplementary

**A.** Dot Plot of GO Term Enrichment for THP-1, infected with Mtb 48 HPI

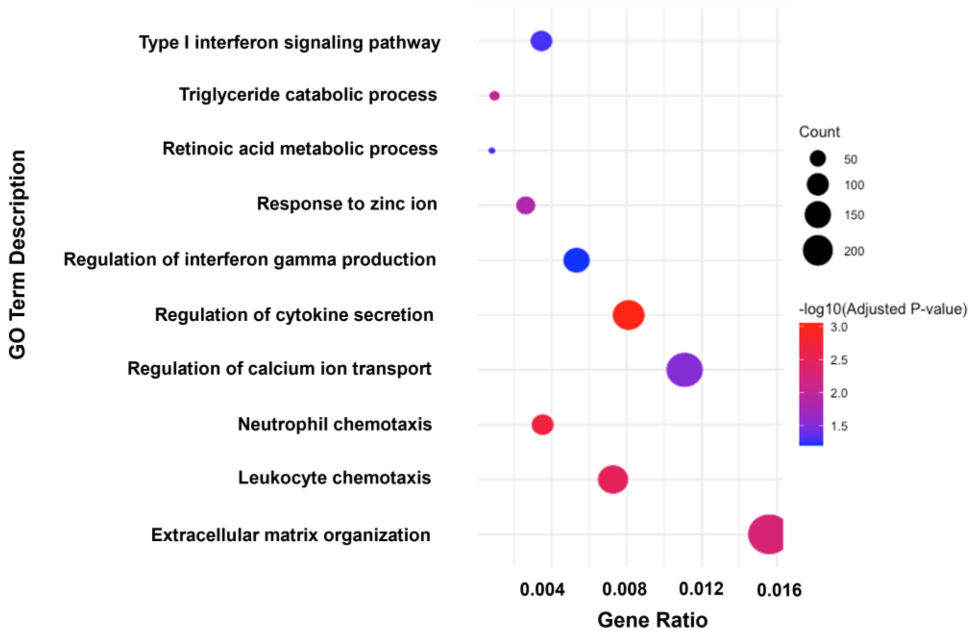


**B.** Dot Plot of GO Term Enrichment for HepG2, infected with Mtb 48 HPI

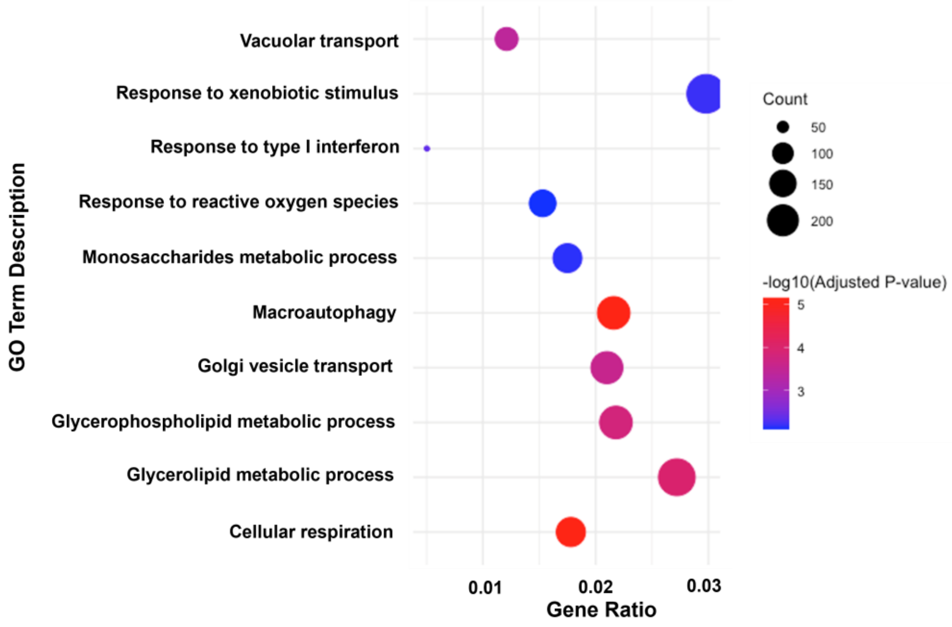


**Figure 5, Supplementary**

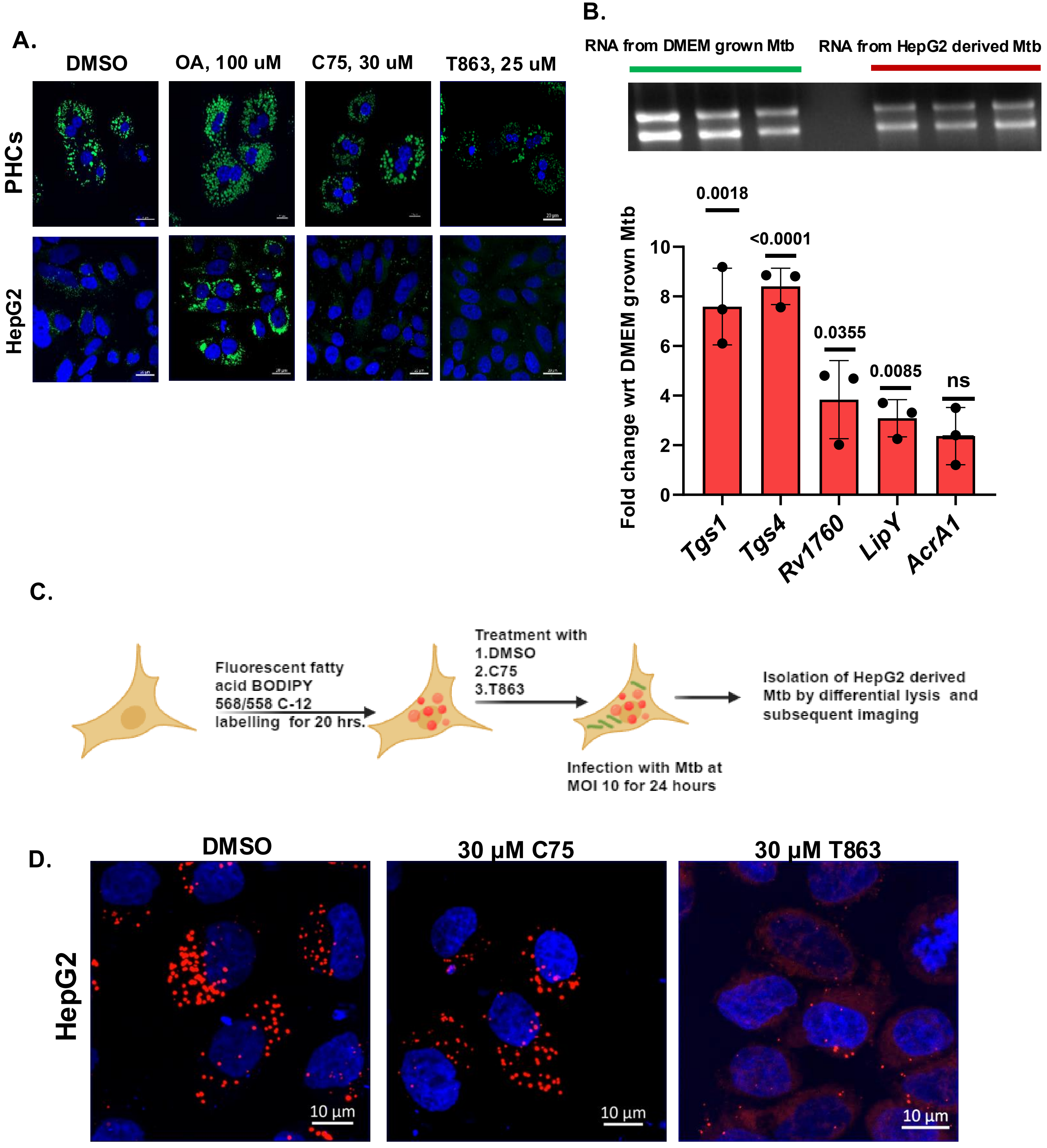


Figure 6, Supplementary

A.

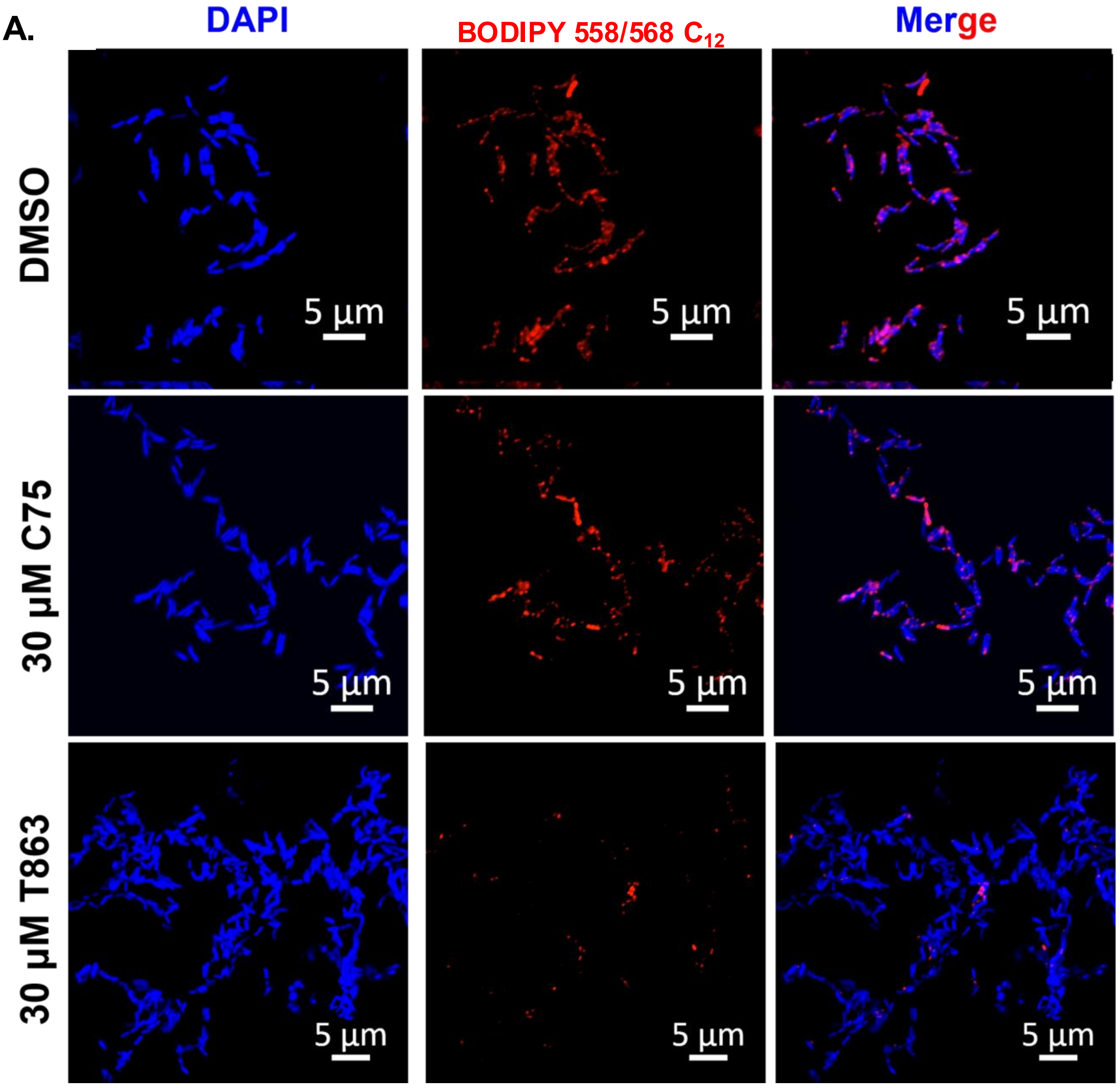


**Figure 7, Supplementary**

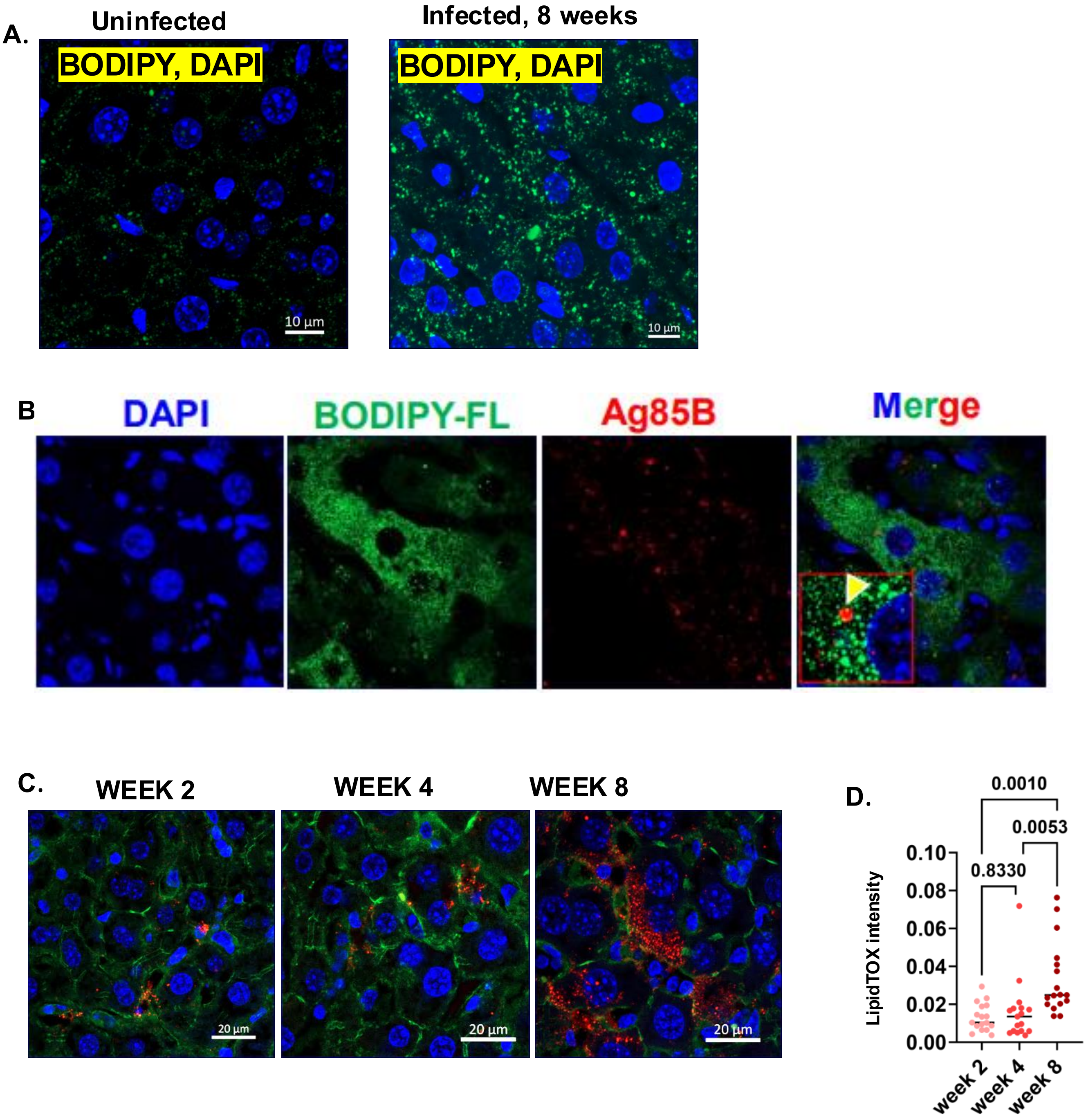


Figure 8, Supplementary

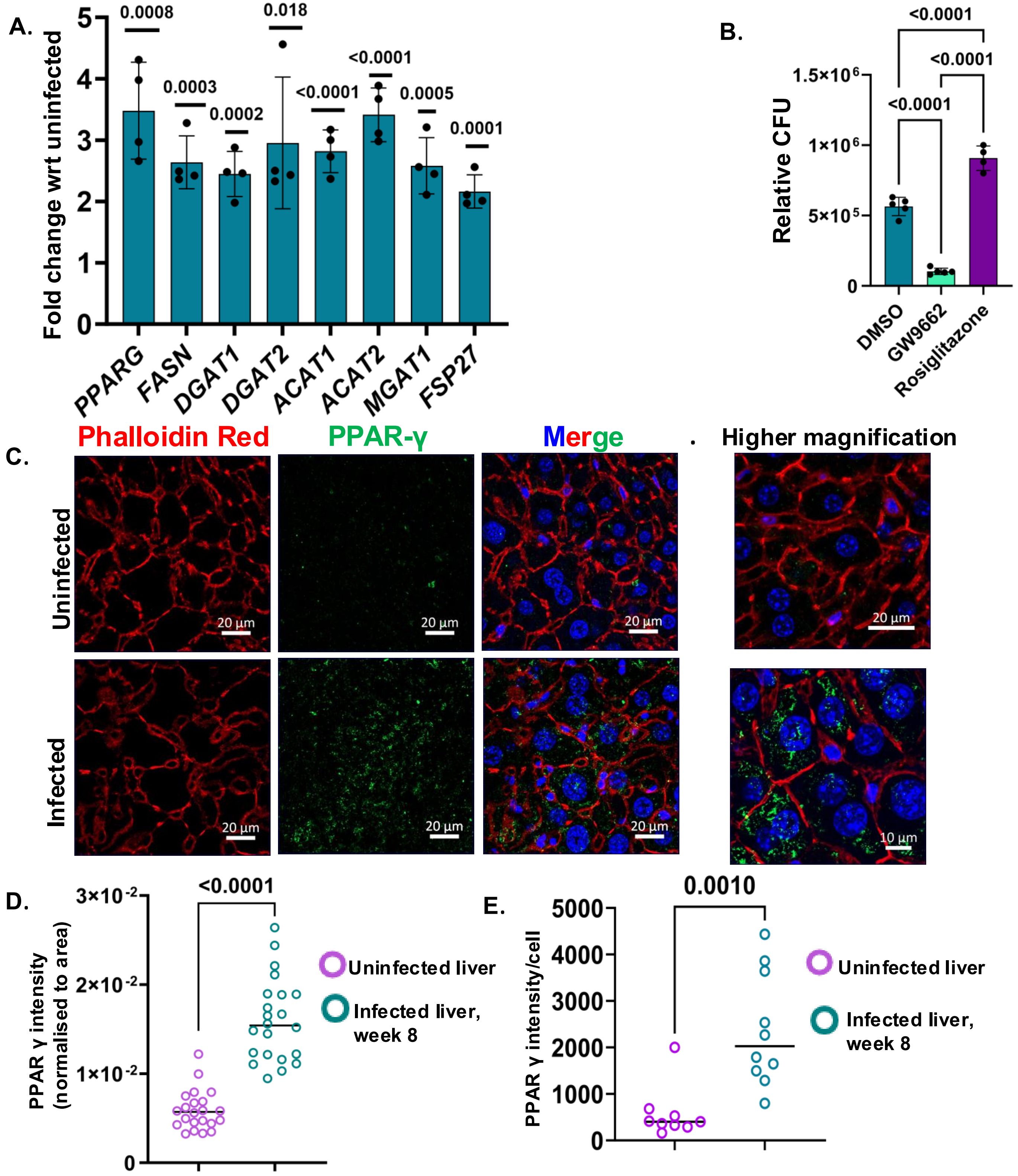


Figure 9, Supplementary

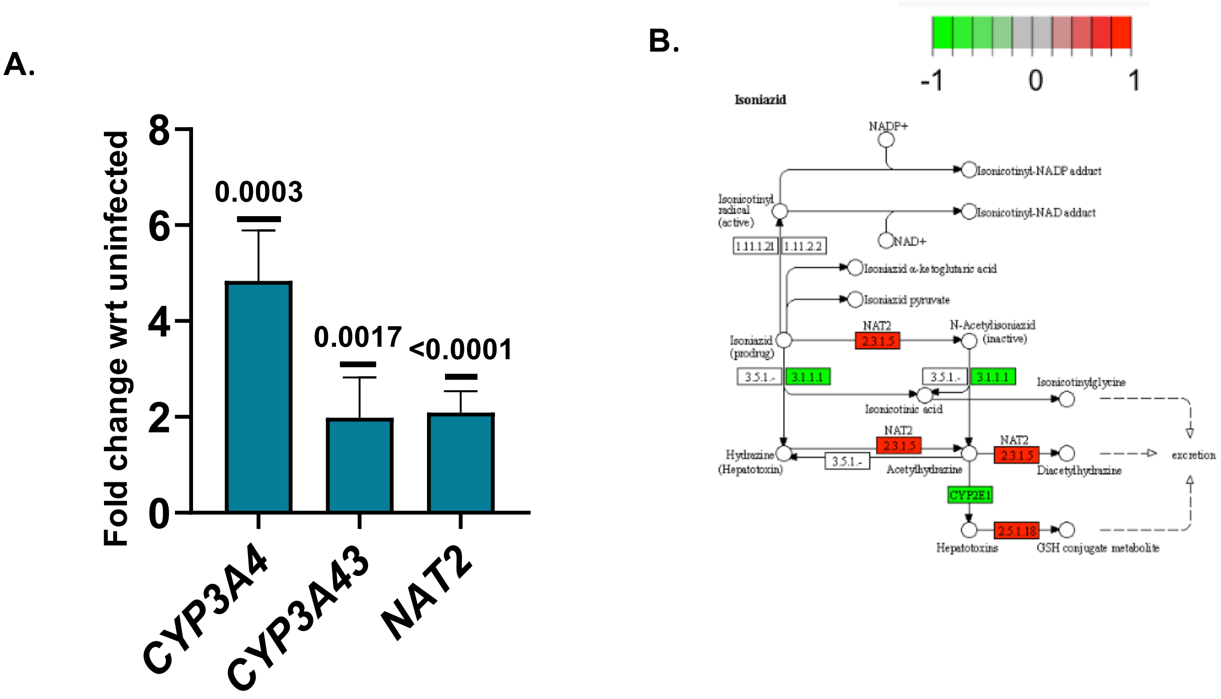


**Figure 10 ,Supplementary**

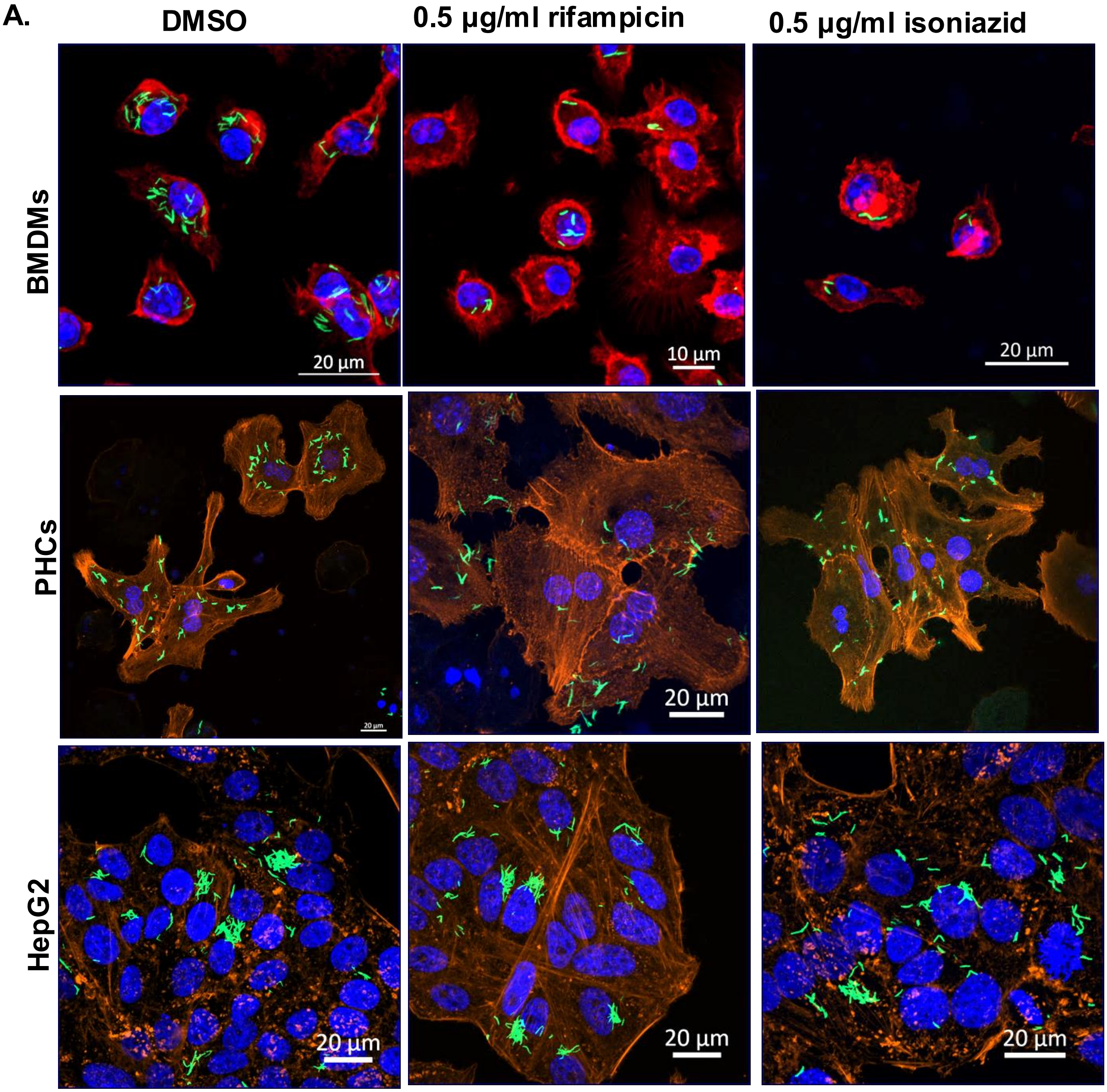


**Figure 11,Supplementary**

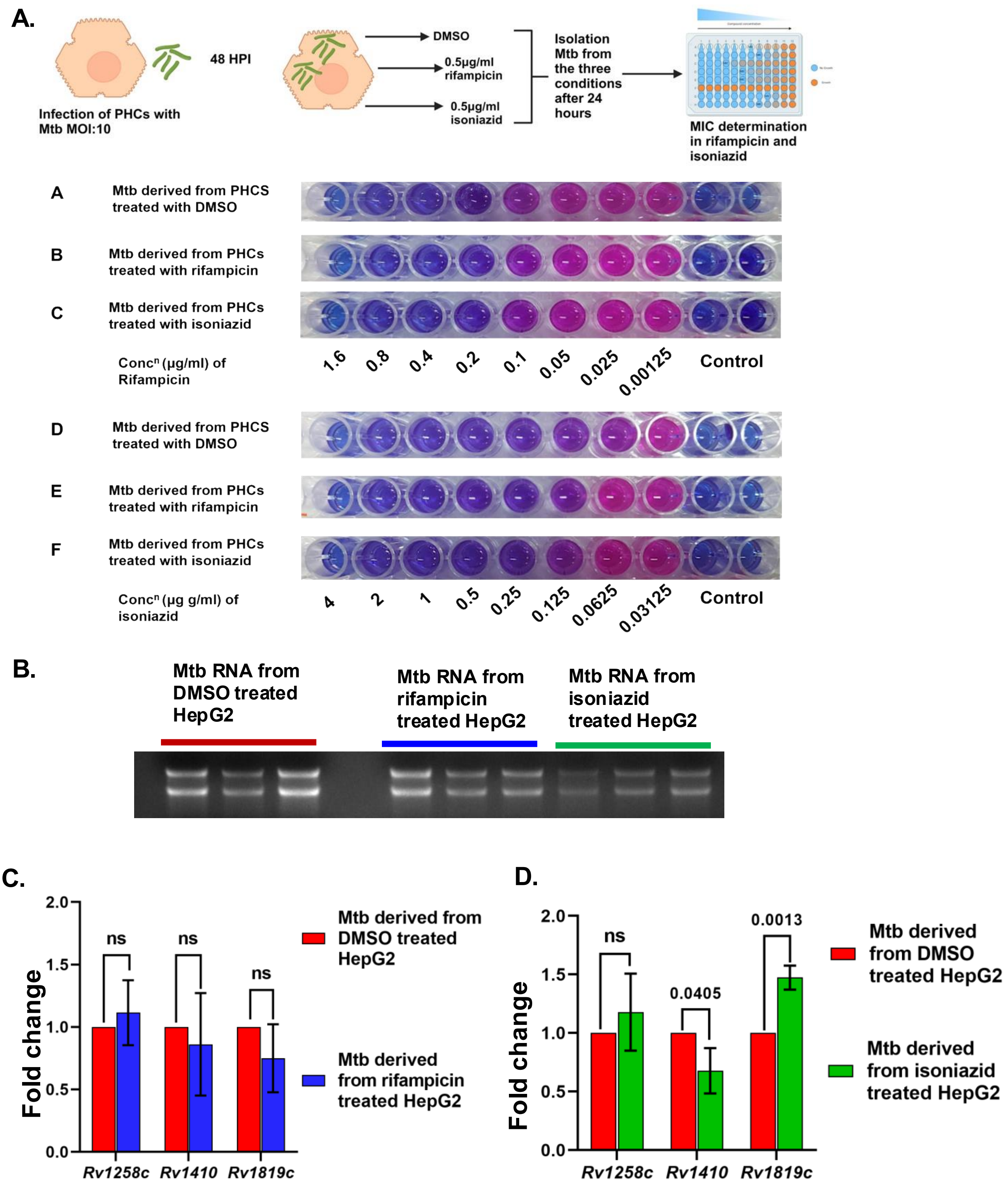


Figure 12 ,Supplementary

**Supplementary Figure legends:**

**Figure S1: Histopathological changes in the liver biopsy samples of miliaryTB patients: (A and B):** A-O staining and FISH in the liver biopsy of the uninfected (control) group shows no Mtb-specific signals. **(C).** H & E staining shows enhanced immune cell infiltration in the liver biopsy of the pulmonary TB patients compared to the uninfected control. **(D).** Dual staining of β-actin (green) and Ag85B (red) using respective antibodies shows the presence of Mtb in hepatocytes and other cells in liver biopsy sections (indicated by yellow arrows) **(E).** H&E staining shows distinct granuloma in the lung section of pulmonary TB patient and **(F).** Acid-fast staining in the same lung section shows a high bacterial load **(G).** A-O staining in the same lung section further validates the presence of high Mtb load. Scale bars in A, B and C are 50 μm and in D is 20 μm. and E and F are 150 μm. Data shown in this figure is representative of 5 patients with pulmonary TB.
